# Gene network module changes associated with the vertebrate fin to limb transition

**DOI:** 10.1101/2021.01.28.428646

**Authors:** Pasan C Fernando, Paula M Mabee, Erliang Zeng

## Abstract

Evolutionary phenotypic transitions, such as the fin to limb transition in vertebrate evolution, result from changes in associated genes and their interactions, often in response to changing environment. Identifying the associated changes in gene networks is vital to achieve a better understanding of these transitions. Previous experimental studies have been typically limited to manipulating a small number of genes. To expand the number of analyzed genes and hence, biological knowledge, we computationally isolated and compared the gene modules for paired fins (pectoral fin, pelvic fin) of fishes (zebrafish) to those of the paired limbs (forelimb, hindlimb) of mammals (mouse) using quality-enhanced gene networks from zebrafish and mouse. We ranked module genes according to their weighted-degrees and identified the highest-ranking hub genes, which were important for the module stability. Further, we identified genes conserved during the fin to limb transition and investigated the fates of zebrafish-specific and mouse-specific module genes in relation to their involvements in newly emerged or lost anatomical structures during the aquatic to terrestrial vertebrate transition. This paper presents the results of our investigations and demonstrates a general network-based computational workflow to study evolutionary phenotypic transitions involving diverse model organisms and anatomical entities.

## 1. Introduction

Phenotypes, such as fin development and limb development, are the result of multiple genes working together in complex biological pathways [1, 2]. Evolutionary modifications in phenotypes due to environmental or other changes involve rewiring gene interactions and their involvements in pathways [2, 3]. Most often, it is likely the network of multiple protein interactions rather than the contribution of a single protein that determines the resulting phenotype [1, 4]. Therefore, investigating the collection of genes and their interactions, i.e., modular gene structure [1], underlying phenotypes is important in evolutionary biology to understand the evolutionary mechanisms that drive phenotypic changes. Gene module analysis has become common in bioinformatics, and the concept of modular evolution has emerged to explain the changes in groups of genes rather than a single gene when studying the evolution of organisms [5–7]. However, most of these studies have focused on smaller protein complexes (typically containing less than 20 proteins) that determine molecular functions and biological pathways [8–10]. Phenotypes, such as fin and limb development, are resulted by a large number of proteins having diverse molecular functions and belonging to several biological pathways. Even the few protein network studies that focus on phenotypes have targeted human diseases [11, 12], and to our knowledge, there have been no evolutionary studies of modules to understand evolutionary phenotypic transitions. As there have been important anatomical changes associated with the vertebrate evolution, such as the fin to limb transition, module evolution studies for anatomical changes are essential to unravel new evolutionary information, which serves as the motivation for our work.

The fin to limb transition is an iconic anatomical change associated with the evolution of terrestrial vertebrates from aquatic fish-like ancestors [13, 14]. According to fossil record, the transformation of fishes into land vertebrates began in the Devonian, 365-408 million years ago [13, 15]. This transformation is associated with many phenotypic changes in addition to the fin to limb transition, including changes in the cranial and axial skeleton [13]. The relationship between homologous anatomical structures of land and aquatic vertebrates is evident from several similar characteristics. For instance, the pectoral fin endoskeleton of panderichthyid fish fossils shows significant similarities with the limb skeletons of terrestrial vertebrates (tetrapods) such as the presence of a proximal humerus and two distal bones [14]. Such evidence indicates that forelimbs and hindlimbs of tetrapods are homologous to pectoral and pelvic fins of fishes, respectively.

Identifying the genetic changes associated with the fin to limb transition is a prominent topic in evolutionary biology [16, 17]. Many wet lab experiments have demonstrated the evolutionary importance of genes such as *shh* [14, 16]. Few computational studies, however, have been targeted on the fin to limb transition [17]. The recent availability of large PPI networks and the ability to perform module analysis through the advancement of network algorithms provide an opportunity for a new perspective on genetic changes associated with the fin to limb transition.

Graph theoretic methods are critical to the study of networks in biology. These methods are enabled by biological knowledge that is represented as computational graphs, such as protein-protein interaction (PPI) networks and biological ontologies. In graph theory, a module is defined as a set of nodes that are highly connected internally and sparsely connected with external nodes [1]. These network modules usually correspond to biological functions that contribute to phenotypes; hence, they are often referred to as ‘functional modules’ in biological vocabulary [6, 18].

There are a number of functional module detection algorithms that can be used to detect modules in a graph [5]. Some methods, such as graph partitioning [19], only consider the network structure and do not require any prior information. For modules that are known to involve a large number of genes in complex phenotypes, it is beneficial to perform module detection using prior knowledge as computational constraints [1, 4, 20]. These methods start from a set of known genes for a given phenotype and expand the module based on the network structure. For example, one of the simplest ways to isolate a functional module by expansion is to assume all the immediate neighbors of the genes associated with the known phenotype are included in the module [1]. However, this method has proven to yield many false positives [1]. Therefore, network-based candidate gene prediction algorithms such as the Hishigaki method [21] and label propagation algorithm [22], which have been shown to be more accurate [1, 4, 21], are often used to predict new candidate genes for inclusion in a module.

One purpose of network analysis is the identification of hub genes, which are defined as important genes that are central to the stability of the module [23, 24]. Hub genes have a higher number of interactions (degrees) than other genes in the module. Their removal is most likely to disrupt the module organization, and thus the biological function(s) or phenotype(s) that is governed by the module. Using network analysis, a set of genes for a function or a phenotype can be transformed into a ranked list that is sorted based on their importance in the module. Usually, the number of interactions a gene forms within the module (degree) is used for the ranking [24].

The quality of PPI network data has been a problematic issue in previous network analyses because of the large portion of spurious PPIs generated by experimental methods, such as high-throughput yeast two-hybrid assay [4, 25]. Therefore, in our previous work [25], we improved the quality of the PPI networks retrieved from the STRING database (STRING, RRID:SCR_005223) [26] by integrating existing experimental knowledge about gene-anatomy relationships available in literature using Uberon anatomy ontology [27]. First, semantic anatomy-based gene networks were generated by calculating the semantic similarity between anatomy terms annotated to different genes, and then, these semantic networks were integrated with the PPI networks for zebrafish and mouse, which improved the candidate gene prediction accuracy for anatomical entities [25]. In this study, we use these improved integrated networks to obtain the most accurate modules.

When considering the evolution of functional modules, most studies have focused on identifying the genes that are retained during evolution, i.e., conserved genes, and their organization in the respective modules [8, 10]. It has been hypothesized that gradual modular changes occur in evolution while maintaining the basic modular structure; this is because dramatic changes in gene interactions may destroy the proper function of an organism [7]. In support of this hypothesis, conserved genes are observed to play an important role in maintaining the stability of the gene modules during evolution [7, 8, 10]. The recruitment and the removal of other genes and the rewiring of biological pathways are often held together by the conserved genes. Performing module analysis allows identification of these important conserved genes, which are often also identified as hub genes [7, 8]. While such conserved module genes may play a role in maintaining gene module structure, species-specific module genes that have been recruited or removed during the evolution may play important roles that contribute to evolutionary transitions [28].

In this work, our goal is to compare PPI network modules associated with fins and limbs to identify the genetic changes, such as the changes in involved genes and their importance, which led to the anatomical changes that characterize the evolution of fins to limbs. From this analysis, we identify genes that are conserved between fins and limbs to understand their roles in the modular evolution, and we predict novel gene candidates with no previously known contributions to the development of paired fins or paired limbs. Further, we identify fin module-specific and limb module-specific genes and investigate their evolutionary roles. This work suggests some evolutionary hypotheses regarding the role of conserved genes versus fin or limb specific genes in the many evolutionary changes in these animals. Finally, this study demonstrates a general network-based computational model to perform gene module comparisons for evolutionary phenotypic transitions.

## 2. Methods

### (a) Selection of the integrated networks for module detection

Based on our previous work [25] of network-based candidate gene prediction using quality-enhanced PPI networks that were generated by four semantic similarity methods (Lin, Resnik, Schlicker, and Wang), the best performing gene networks for zebrafish and the mouse were selected for this project. These are referred to as ‘zebrafish integrated network’ and ‘mouse integrated network’ from herein.

### (b) Detection of network modules

For module detection, genes with direct annotations to the pectoral fin (UBERON:0000151), forelimb (UBERON:0002102), pelvic fin (UBERON:0000152), and hindlimb (UBERON:0002103) were used as prior information and their anatomical profiles were extracted from the Monarch Initiative repository (https://monarchinitiative.org/; RRID:SCR_000824) (06/20/2018)[29]. In addition, genes that were annotated to the parts (e.g., pectoral fin lepidotrichium and pectoral fin radial skeleton are parts of the pectoral fin) and the developmental precursors (pectoral fin bud, pelvic fin bud, forelimb bud, and hindlimb bud) of the above entities were extracted using the Uberon anatomy ontology relationships. The genes directly annotated to the anatomical entity of interest or annotated to a part or developmental precursor of the entity are collectively referred to as ‘genes with original annotations’.

Beginning with the genes with original annotations, gene modules for the anatomical entities of interest were identified by predicting novel genes using the Hishigaki network-based candidate gene prediction method [21, 25]. First, the network-based candidate gene prediction performance for each anatomical entity of interest was evaluated using leave-one-out cross-validation [25], and ROC and precision-recall curves were generated. Then, a prediction precision threshold was used to predict new candidate genes. A trial and error method was used to select the best precision threshold for each gene module.

After predicting the candidate genes, the modules were extracted for the pectoral fin and the pelvic fin from the zebrafish integrated network and for the forelimb and the hindlimb from the mouse integrated network. The extracted modules were visualized using the Cytoscape software [30] (Cytoscape, RRID:SCR_003032).

### (c) Validation of the predicted genes

The predicted candidate genes could be validated using either experimental methods, such as gene knockdown [31], or computational methods such as the one used in this work. First, the predicted genes for the pectoral fin and pelvic fin modules in zebrafish were compared with the orthologous genes in the forelimb and hindlimb modules in mouse and *vice versa* to determine whether they were annotated to a homologous anatomical entity.

Second, enrichment analyses were performed to confirm for each module, whether the predicted genes shared similar Biological Process terms from Gene Ontology (GO-BP) as the genes with original annotations. Enrichment analyses were also performed to confirm for each module, whether the predicted genes shared similar Uberon anatomy annotations as the genes with original annotations.

Third, the weighted degree distributions of the predicted genes were compared with the weighted degree distributions of the genes with original annotations in each module. If the predicted genes have a higher weighted degree distribution, it indicates that the predicted genes have a similar or a higher importance as genes with original annotations.

### (d) Comparison of the network modules

To study the fin to limb transition and identify the modular changes, the pectoral fin and pelvic fin modules of the zebrafish were compared with the forelimb and hindlimb modules of the mouse, respectively.

Teleost fishes, such as the zebrafish, have more genes than tetrapods, such as the mouse. A whole genome duplication event is proposed to have occurred at the origin of actinopterygian fishes, i.e., the teleost genome duplication [32]; hence, most of the mouse genes have duplicated copies in the zebrafish. To perform the module comparison, the gene ortholog mappings between mouse and zebrafish genes were retrieved from the Zebrafish Information Network [33] (ZFIN, 06/26/2018) (https://zfin.org/downloads) (Zebrafish Information Network, RRID:SCR_002560). During the comparison, if multiple zebrafish orthologs were present in a zebrafish module for a single mouse gene, all zebrafish orthologs were retained. By performing the module comparison, conserved genes (genes that are common to the two modules), zebrafish module-specific genes, and mouse module-specific genes were identified.

In network analysis, the degree of a gene (the number of interactions of the gene) is often used as an important metric [10, 24]. Genes with higher degrees in a module, i.e., hub genes, are considered more important because they have more interactions with other module genes and removal of such a gene from the module may significantly affect the integrity of the module [23]. When analyzing networks with weights assigned for interactions (weighted networks), such as the integrated networks used here, weighted degree is preferred over the degree because it considers the different interaction weights rather (equation 1) than counting the number of interactions for a specific node [24].

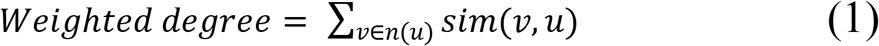

In equation 1, *n(u)* is the neighborhood of the gene of interest (*u*) and *v* iterates through all the neighbors of gene *u*. The gene similarity score for the interaction between genes *v* and *u*, which is represented by *sim(v,u)*, is used for the interaction weight. Weighted degree of gene *u* is the summation of all weights of interactions between gene *u* and all its neighbors.

The weighted degree for each gene in a module was calculated, and the genes were ranked accordingly. During the comparisons, the weighted degree of each zebrafish module gene was compared with the corresponding mouse ortholog. However, due to the size differences of the zebrafish and mouse modules, the weighted degree of each gene had to be normalized by the total number of genes in each module. Then, normalized weighted degree distributions for conserved genes, zebrafish module-specific genes, and mouse module-specific genes were compared for pectoral fin *versus* forelimb and pelvic fin *versus* hindlimb to study the relative importance of genes in each group.

The fate of the zebrafish module-specific genes in mouse was investigated by extracting mouse orthologs for the pectoral and pelvic fin module-specific genes and performing enrichment analyses using Uberon and GO-BP terms. Similarly, the roles of the mouse module-specific genes in zebrafish were investigated using zebrafish orthologs for the forelimb and hindlimb module-specific genes. The DAVID (https://david.ncifcrf.gov/) (DAVID, RRID:SCR_001881) online functional enrichment analysis tool was used to perform gene set enrichment analysis using GO-BP terms. DAVID uses Fisher’s exact test [34] to perform enrichment analyses. Although the GO is widely used for enrichment analysis, anatomy ontologies are rarely used. To perform enrichment analysis using the Uberon anatomy ontology and Fisher’s exact test, a Python program (Uberon enrichment analysis program) was developed and used. Ontology terms with p-values less than 0.05 were considered as enriched terms.

## 3. Results and discussion

### (a) Selection of the integrated networks for module detection

The integrated networks generated using the Lin and Schlicker methods were selected for module detection for zebrafish and mouse, respectively because they outperformed other integrated networks based on the results of our previous work [25]. The zebrafish Lin integrated network contained 17,394 genes and 730,855 interactions and the mouse Schlicker integrated network contained 18,002 genes and 613,671 interactions [25].

### (b) Detection of network modules

The statistics showing the number of genes with original annotations to each anatomical entity are given in electronic supplementary material, table S1. The total number of genes for the pectoral fin (198) and the forelimb (267) were comparatively similar than the total number of genes for the pelvic fin (15) and the hindlimb (777). Detection of the pelvic fin module was challenging because of the low number of original gene annotations. Unlike the limb development in the mouse, where forelimb and hindlimb buds emerge at the same timepoint, the pelvic fin buds emerge at a much later stage than the pectoral fin bud [35]. This may have been a potential reason for fewer annotations to the pelvic fin; the studied gene disruptions may have killed the larval zebrafish before the pelvic fin develops or the larvae may have been sacrificed at a pre-determined early stage.

The ROC and precision-recall curves generated for each anatomical entity during the network-based candidate gene prediction evaluations are given in electronic supplementary material, figures S1 and S2, respectively. According to the curves, all anatomical entities except the pelvic fin show high accuracies for network-based candidate gene predictions (the AUC values of ROC curves were higher than 0.85). This shows the high reliability of the network candidate gene predictions. The lower performance for the pelvic fin could be due to its low number of original gene annotations. It has been shown that the prediction accuracy improves with the size of the dataset/number of gene annotations, and anatomical entities with a low number of gene annotations can lead to lower AUC values [36].

The statistics for the extracted gene modules are given in electronic supplementary material, table S1. The genes with original annotations that were lost during the module extraction are listed in electronic supplementary material, table S2. A high precision threshold of 0.7 was used for candidate gene predictions for pectoral fin, forelimb, and hindlimb modules. The precision threshold for the pelvic fin was lowered to 0.05 to make the number of genes in the pelvic fin and the forelimb modules approximately similar.

The visualizations of the resulting modules for the pectoral fin, pelvic fin, forelimb, and hindlimb are given in electronic supplementary material, figures S3, S4, S5, and S6, respectively. The companion Cytoscape network files for these modules are available in electronic supplementary material, files S1, S2, S3, and S4. The genes in the pectoral fin, pelvic fin, forelimb, and hindlimb modules ranked based on the weighted degree are listed in electronic supplementary material, files S5, S6, S7, and S8, respectively.

### (c) Validation of the predicted genes

The list of predicted genes for pectoral fin, pelvic fin, forelimb, and hindlimb modules are given in electronic supplementary material, tables S3, S4, S5, and S6, respectively. Of the 45 predicted genes for the pectoral fin, 14 genes had mouse orthologs that were associated with the forelimb (9 direct annotations, 2 annotations only to the parts or the developmental precursors, and 3 predicted genes). Of the 605 predicted genes for the pelvic fin, 78 genes had mouse orthologs that were associated with the hindlimb (46 direct annotations, 20 annotations only to the parts or the developmental precursors, and 12 predicted genes). Of the 18 predicted genes for the forelimb, 6 genes had mouse orthologs that were associated with the pectoral fin (2 direct annotations, 1 annotation only to the parts or the developmental precursors, and 3 predicted genes). Of the 32 predicted genes for the hindlimb, 12 genes had mouse orthologs that were associated with the pelvic fin (all 12 were predicted genes). These results indicate that the orthologs of the predicted genes are annotated to homologous anatomical entities, providing a certain level of validation for the predicted genes.

The enriched GO-BP terms that are common to the predicted genes and genes with original annotations to pectoral fin, pelvic fin, forelimb, and hindlimb are listed in electronic supplementary material, tables S7, S8, S9, and S10, respectively. The enriched Uberon terms that are common to the predicted genes and genes with original annotations to pectoral fin, pelvic fin, forelimb, and hindlimb are listed in electronic supplementary material, tables S11, S12, S13, and S14, respectively. There were several common enriched GO-BP terms for all the modules, some of which were related to paired fins and limbs, such as pectoral fin development, fin development, embryonic limb morphogenesis, embryonic digit morphogenesis. Some of the common enriched Uberon terms, such as median fin fold, ventral fin fold, caudal fin, appendicular skeleton and limb, were related with fin or limb development.

The boxplot comparisons of the weighted degree distributions for the predicted genes *versus* genes with original annotations for the pectoral fin, pelvic fin, forelimb, and hindlimb modules are shown in figure 1. In all the modules, the weighted degree distributions of the predicted genes were higher than the genes with original annotations. This indicates that predicted genes as a group are important in the modules and central to the function of the modules, which supports the biological significance of the predicted genes.

**Figure 1.**
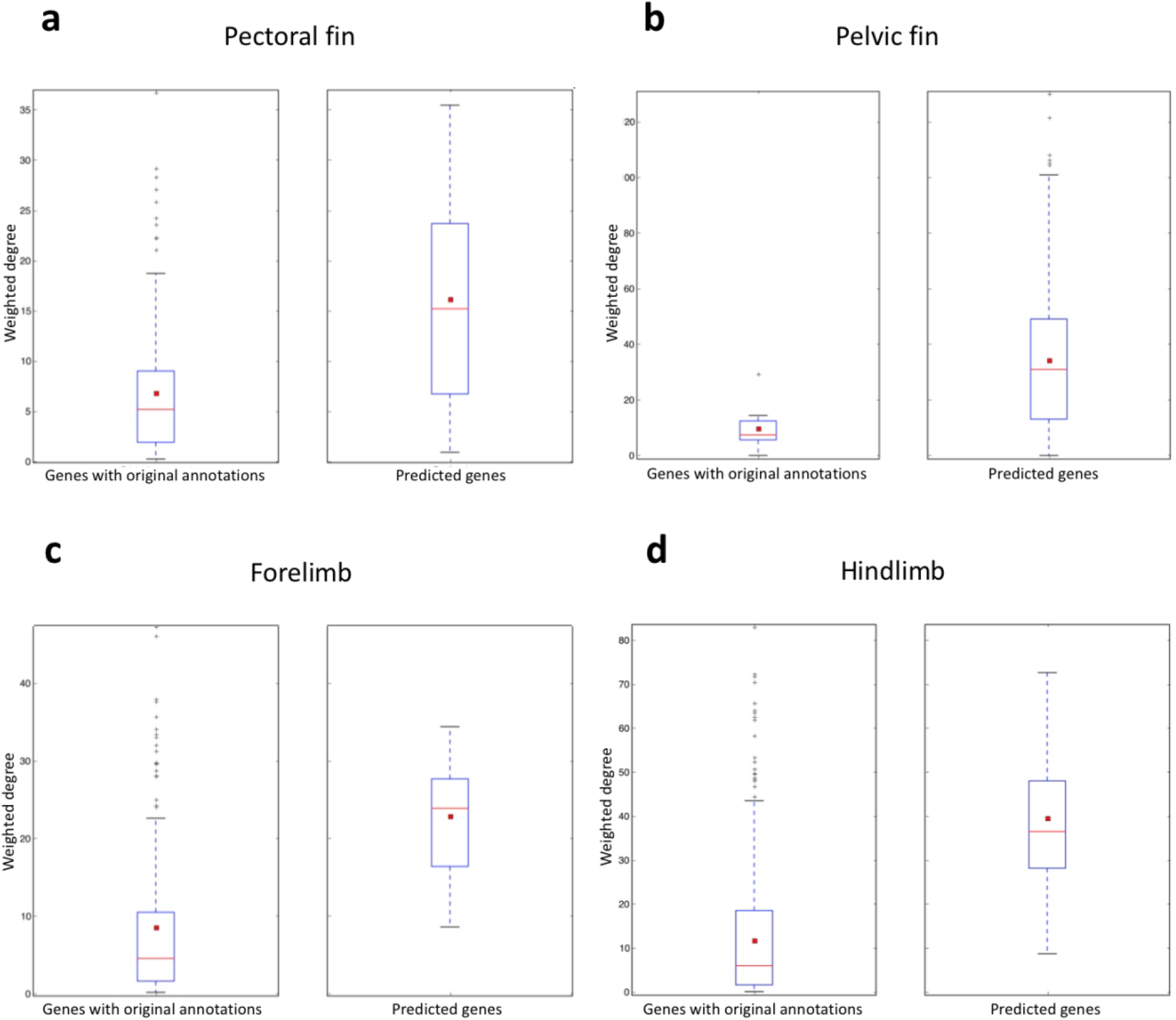
The boxplot comparisons of the weighted degree distributions for the predicted genes *versus* genes with original annotations for each module. In the boxplots, the red line and the square represent the median and mean, respectively.

### (d) Comparison of the network modules

#### (i) Pectoral fin and forelimb comparison

According to the comparison, 183 genes were specific to the pectoral fin module, 207 genes were specific to the forelimb module. 37 genes were shared (conserved genes) between the pectoral fin and forelimb (electronic supplementary material, table S15, figure 2).

**Figure 2.**
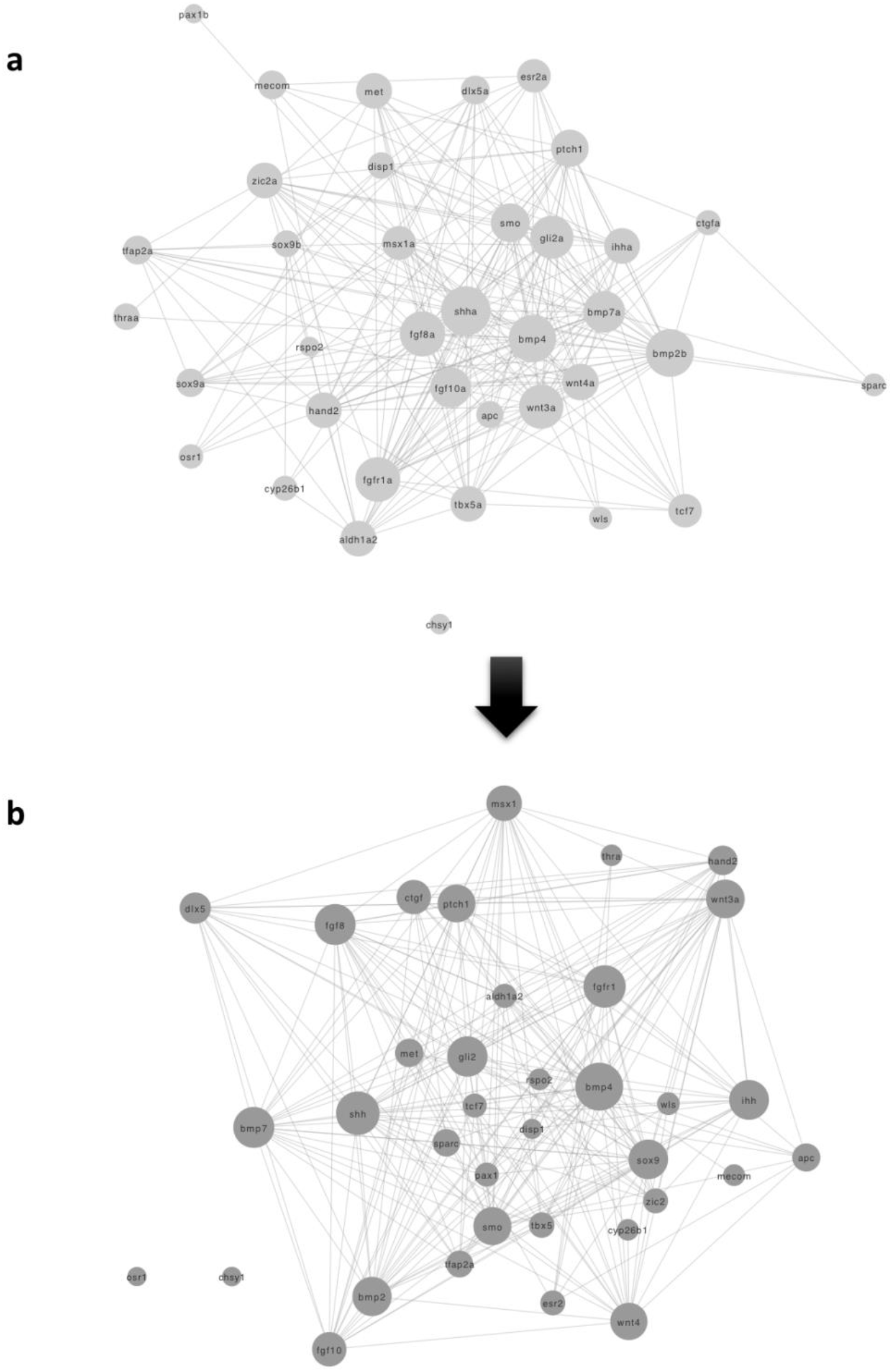
Networks of the 37 conserved genes that are common to and extracted from (a) the pectoral fin module and (b) the forelimb module. Node size is proportional to the degree (number of interactions) of the gene. Hub genes, such as *bmp4*, *shh*, *smo*, *bmp7, sox9*, and *gli2*, are shown in larger node sizes. The arrow represents the direction of modular evolution.

In the pectoral fin module, the top-ranked hub gene based on the weighted degree was *shha* (sonic hedgehog a) (electronic supplementary material, file S5), whose role has been well-documented in pectoral fin development [16]. Its ortholog, *Shh*, is important in the development and morphogenesis of limbs in tetrapods including humans [37], and it was also highly ranked in the forelimb module (4^th^, see electronic supplementary material, table S15). The loss or gain of activity in the sonic hedgehog signaling pathway in tetrapods results in lost, gained, or malformed limbs [37]. The *shh* gene has long been considered an important gene associated with fin to limb transition because it is important in the morphological patterning of paired fins and limbs [14].

The highest-ranking gene in the forelimb module was *bmp4* (bone morphogenetic protein 4), another gene closely associated with limb formation and morphogenesis in tetrapods [38]. Mutations in *bmp4* affect the *bmp4* signaling pathway to cause abnormalities in limb and digit formation in tetrapods [38]. *Bmp4* was ranked 2^nd^ in the pectoral fin module (electronic supplementary material, table S15) and was predicted during module detection.

When considering the conserved genes (figure 2), some of the important hub genes in the pectoral fin module, such as *shha*, *bmp4*, *bmp2b*, and *bmp7a*, had retained their importance demonstrated by their higher ranks based on the weighted degree in the forelimb module (electronic supplementary material, table S15). Other genes such as *sox9* were elevated in rank during the transition from pectoral fin to forelimb. In the pectoral fin module, *sox9a* and *sox9b* genes were ranked 83^rd^ and 104^th^, respectively, while in the mouse, the ortholog *sox9* was elevated to 15^th^ (electronic supplementary material, table S15). *Sox9* is well known to be involved with digit patterning in the limbs of tetrapods due to its participation in the a bmp-sox9-wnt Turing network [17, 39]. Because digits emerged after the transition from fins to limbs [13, 14], the involvement of *sox9* in a digit patterning pathway could have increased the number of interactions with other genes in the forelimb module, and hence, the increased importance.

A boxplot comparison of normalized weighted degree distributions for pectoral fin module-specific genes, pectoral fin conserved genes (genes of the pectoral fin in common with forelimb), forelimb conserved genes (genes of the forelimb in common with pectoral fin), and forelimb module-specific genes are given in figure 3. The conserved genes in both modules have higher normalized weighted degree distributions compared to the respective module-specific genes. This indicates that the conserved genes share more interactions within the module as a group and are more central to modular stability. From an evolutionary point of view, during the transition from pectoral fin to the forelimb, it appears that genes with higher degrees in the pectoral fin module, such as *shha*, *bmp4*, were conserved in the forelimb and new forelimb module-specific genes were recruited surrounding those conserved genes.

**Figure 3.**
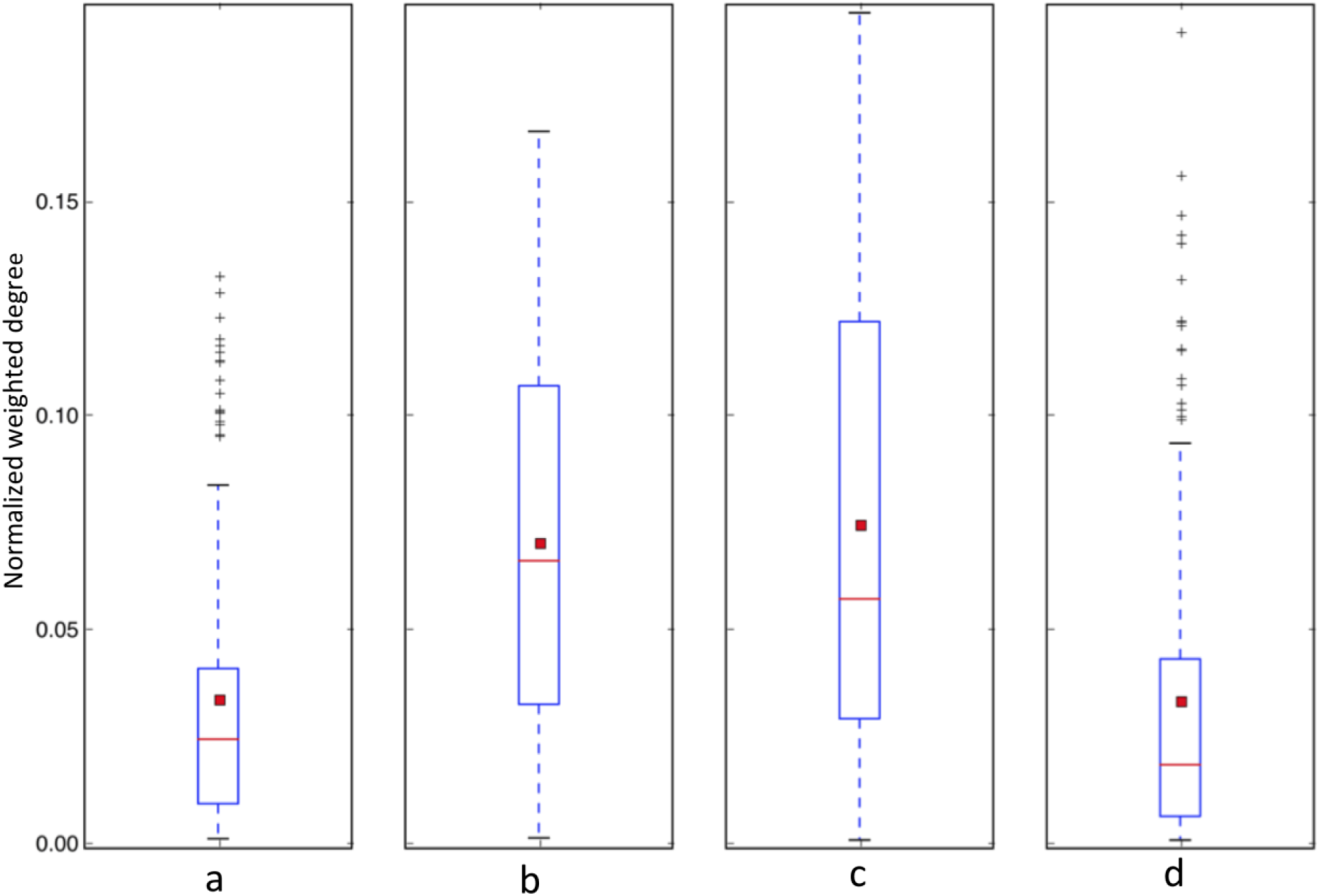
Boxplot comparison of normalized weighted degree distributions for (a) pectoral fin module-specific genes, (b) pectoral fin conserved genes, (c) forelimb conserved genes, and (d) forelimb module-specific genes. In the boxplots, the red line and the square represent the median and mean, respectively.

#### (ii) Pelvic fin and hindlimb comparison

According to the comparison, 536 genes were specific to the pelvic fin module, 601 genes were specific to the hindlimb module, and 81 genes were conserved between pectoral fin and forelimb modules. (electronic supplementary material, table S16 and figure 4).

**Figure 4.**
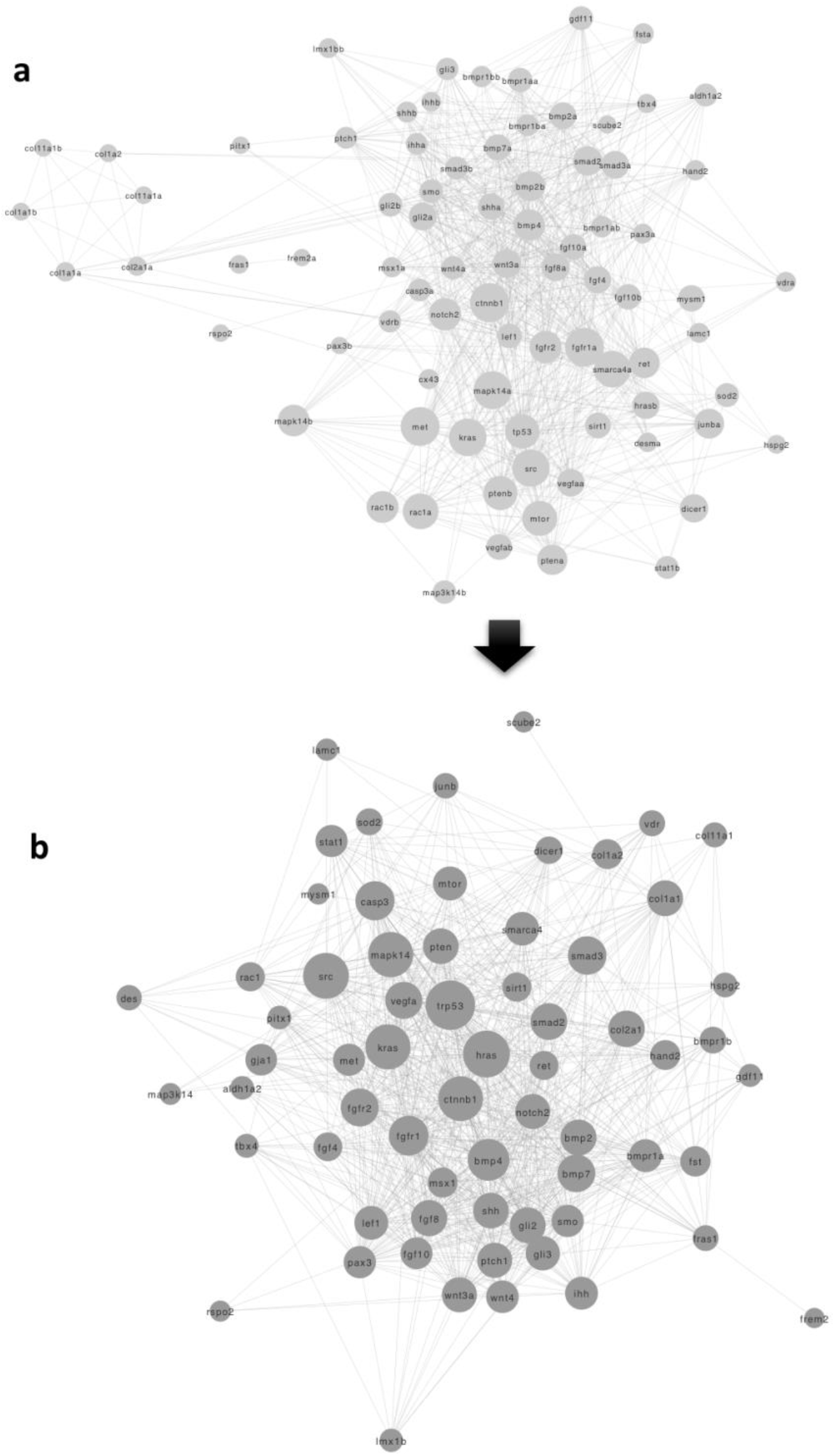
Networks of the 81 conserved genes common to and extracted from (a) the pelvic fin module and (b) the hindlimb module. Node size is proportional to the degree (number of interactions) of the gene. Hub genes, such as *bmp4*, *shh*, *ctnb1*, *bmp7, trp53*, and *hras*, are shown in larger node sizes. The arrow represents the direction of modular evolution.

In the pelvic fin module, the highest-ranking gene was *hsp90ab* (predicted) (electronic supplementary material, file S6). Although it is a heat shock protein and does not have known effects on the pelvic fin, studies have shown that the inhibition of its expression causes defects in zebrafish, especially in eye development [40]. Furthermore, the disruption of *hsp90ab* expression has been associated with caudal fin fold defects in the zebrafish [40], which is not recorded in the ZFIN or the Monarch Initiative repository. Our computational results, together with a noted effect on a fin, indicate that *hsp90ab* is a prime new candidate gene for pelvic fin development that may have a key role in the module stability.

The top ranked hub gene in the hindlimb module based on weighted degree was *trp53* (electronic supplementary material, file S8), which has been associated with embryonic hindlimb development in mouse [41]. When *trp53* is disrupted, mouse limbs are deformed [42]. *Trp53* was also found in the pelvic fin module (predicted gene) but it had a lower rank (24th) based on the weighted degree (figure 4 and electronic supplementary material, table S16).

When comparing the conserved genes between the pelvic fin and the hindlimb modules (electronic supplementary material, table S16 and figure 4), several that are central to the modular stability were identified. For example, the *ctnnb1* gene, predicted and ranked 4^th^ in the pelvic fin module, was also highly ranked (3^rd^) in the forelimb module. *Ctnnb1* is essential for the β-catenin pathway, which is necessary for the hindlimb initiation in the mouse [43]. Although it does not have known association to either of the paired fins in the zebrafish, it is known to be essential in fish development [44].

A boxplot comparison of normalized weighted degree distributions for pelvic fin module-specific genes, pelvic fin conserved genes, hindlimb conserved genes, and hindlimb module-specific genes is given in figure 5. The conserved genes in both modules show higher normalized weighted degree distributions compared to their respective module-specific genes. As observed for the pectoral fin, this indicates the higher importance of the conserved genes for the stability of the modules.

**Figure 5.**
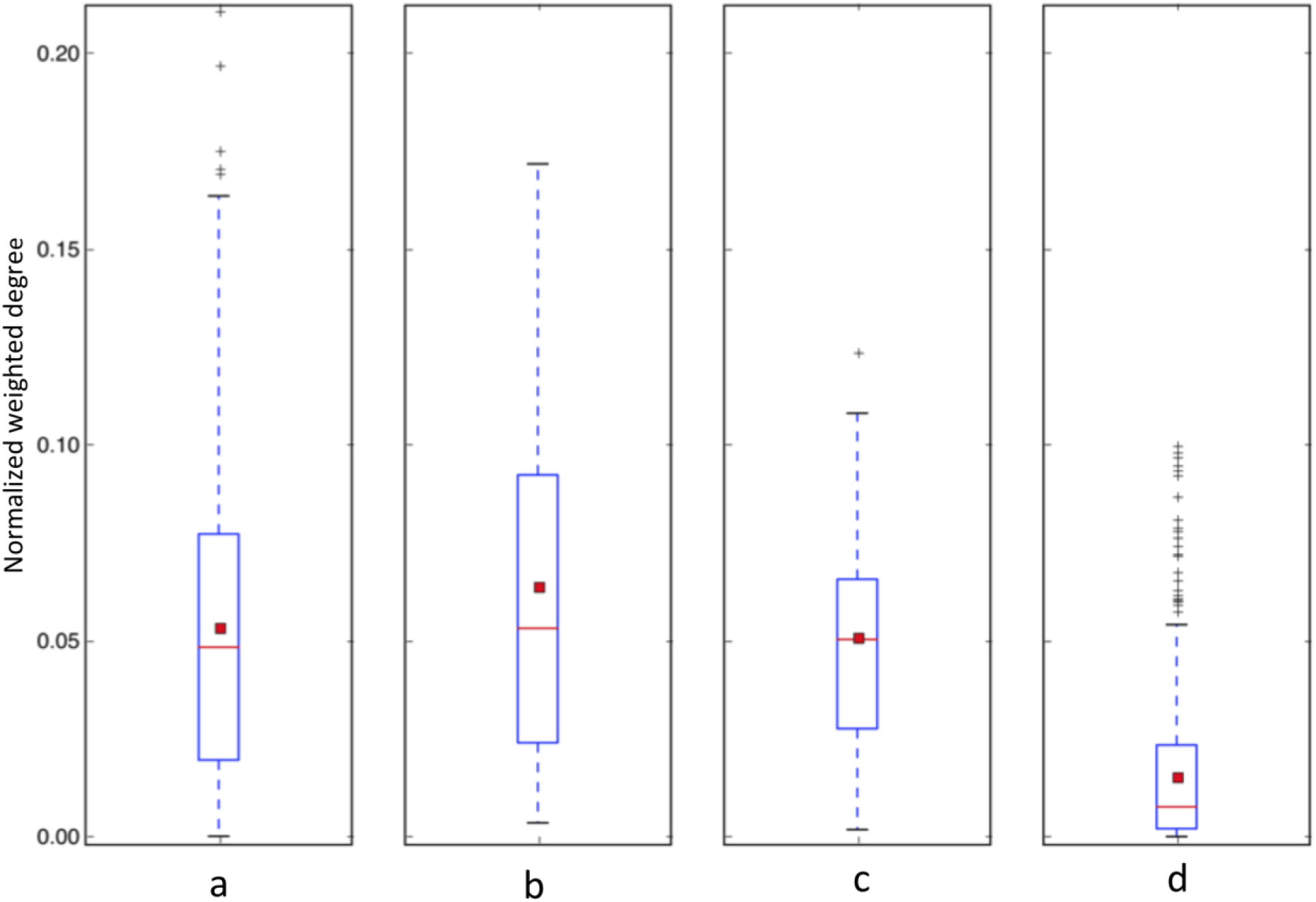
Boxplot comparison of normalized weighted degree distributions for (a) pelvic fin module-specific genes, (b) pelvic fin conserved genes, (c) hindlimb conserved genes, and (d) hindlimb module-specific genes. In the boxplots, the red line and the square represent the median and mean, respectively.

#### (iii) The fate of zebrafish paired fin module-specific genes in the mouse

A large number of zebrafish fin module genes (183 for pectoral fin and 536 for pelvic fin) were not included in the mouse limb modules (electronic supplementary material, files S5 and S6), implying these genes had not been maintained in limb development. To understand the roles of those zebrafish pectoral and pelvic fin module-specific genes in the mouse, the enriched GO-BP and Uberon terms for the mouse orthologs for these fin module-specific genes are given in electronic supplementary material, tables S17, S18, S19 and S20. They were enriched for a number of novel anatomical entities and related biological processes unique to tetrapods [13] (electronic supplementary material, tables S21 and S22).

For instance, the pectoral fin module-specific gene *lef1*, an important (ranked 7^th^) member in the pectoral fin module, is involved with palate development, trachea gland development, and associated with neck-related phenotypes [45, 46]. The neck evolved in tetrapods and allowed them to support the head, which was crucial for their success in land [47, 48]. In the pelvic fin module-specific genes, *mapk1* (ranked 12th) is also associated with neck-related phenotypes, such as thymus development and trachea formation [49, 50]. It is additionally involved with the lung phenotypes and the development of the lung [50], another structure which progressively evolved in tetrapods that enabled them to breath and thrive in terrestrial environments [51]. *Lama5*, a gene found in both the pectoral fin and pelvic fin modules, is an example of another module-specific gene that is involved with lung development in the mouse [52]. Furthermore, it is also involved with hair follicle development and hair-related phenotypes [53], which are other anatomical entities specific for mammals [54]. These examples point to the possibility that many of genes used in fin development were recruited in the development of novel anatomical entities that enabled tetrapods to thrive in a terrestrial environment.

#### (iv) The role of mouse module-specific limb genes in the zebrafish

A large number of module-specific genes for the forelimb and hindlimb (207 for forelimb and 601 for hindlimb) did not appear in pectoral fin or pelvic fin modules (electronic supplementary material, files S7 and S8), and the question of their developmental function in the zebrafish occurred. To understand the function of the limb module-specific genes in zebrafish, the enriched GO-BP and Uberon terms for the limb module-specific genes are given in electronic supplementary material, tables S23, S24, S25 and S26. According to the enrichment analyses, these mouse limb module-specific genes were enriched to the head of the zebrafish, specifically, the jaw skeleton and post-hyoid pharyngeal arch skeleton (electronic supplementary material, tables S27 and S28). The latter region includes the gill chamber and contains parts such as gill rakers [55] that have been lost in tetrapods. For instance, *fst* is a crucial forelimb module-specific gene, which has a zebrafish ortholog (*fsta*) with phenotypes related to splanchnocranium [56] and post-hyoid pharyngeal arch skeleton [57] that supports the gill chamber. Furthermore, *twist1* is module-specific for both forelimb and hindlimb, and it has two zebrafish orthologs (*twist1a* and *twist1b*) that are involved with pharyngeal system development [58].

There are some mouse module-specific genes, e.g., *tgfbr3*, which are involved in the development of both the forelimb and the hindlimb, that is associated with the development of the caudal fin in zebrafish [59]. Another example, *lep*, which is module-specific for both forelimb and hindlimb, is associated with otolith development in zebrafish [60]. Otoliths are located in the inner ear cavity of all teleost fishes where they aid in hearing and serve as balance organs [61]. The enrichment analyses point to the possibility that genes associated with various fish-specific structures such as gill arches and the caudal fin, were recruited for limb development as they were lost during the transition to tetrapods.

## 4. Conclusion

The goal of this work was to study the modular changes associated with the fin to limb transition using gene networks. This computational study expanded the number of genes that could be analyzed compared to wet lab methods and enabled the study of gene network structure rather than individual genes. Employing the quality-enhanced integrated networks ensured that the module detections, gene predictions, and identification of important genes in the modules were accurate, as evidenced from the results. To our knowledge, this is the first work that uses PPI networks to study the fin to limb transition. We discovered important information such as the hub genes responsible for the stability of paired fin and limb modules and changes in the importance of module genes associated with the transition. Some of the module genes were predicted during the module detection, with evidence confirming their involvement with the respective fins or limbs. This paper tabulates rankings of module genes based on their importance, predicted candidates, and comparisons of the importance of the module genes, which will be useful for future studies on fin to limb transition.

Furthermore, we discovered that the conserved genes were more likely to be hub genes than the module-specific genes. Thus, it appeared that during the fin to limb transition, most of the crucial hub genes of fin modules were conserved in the limb, and limb-specific genes were recruited to surround this conserved ‘appendage’ core network. Moreover, our data suggested that zebrafish fin module-specific genes were additionally employed in anatomical structures that emerged after the aquatic to terrestrial vertebrate transition, such as lung and neck. Furthermore, the evidence implied that mouse limb module-specific genes were involved with anatomical structures, such as the gill rakers in the zebrafish that were lost during the transition. These results provide the groundwork for evolutionary developmental biologists to experimentally investigate aforementioned hypotheses. Most importantly, this work demonstrates how gene networks can be used to study evolutionary phenotypic transitions and this computational workflow can be used to perform large-scale network analyses to study evolutionary transitions involving any model organism and anatomical entity with sufficient data, which is a valuable addition to evolutionary biology.

## Supporting information

Electronic supplementary material tables and figures

Electronic supplementary material files

## Data accessibility

The network files and the anatomy profiles used for the candidate gene predictions are available at https://doi.org/10.6084/m9.figshare.13589579.v1 and the Python scripts used for this analysis are available at https://doi.org/10.5281/zenodo.4445583.

## Authors’ contributions

All authors planned and designed the experiments. PCF wrote the Python scripts for the analysis and performed the experiments under the supervision of PMM and EZ. All authors analyzed the results, read and approved the final manuscript.

## Acknowledgements

The authors thank J. P. Balhoff for assisting with the retrieval of gene-anatomical entity relationships from the Monarch Initiative repository. The authors thank T. J. Vision, D. Goodman, B. Wone, W. M. Dahdul, and L. M. Jackson for their assistance and helpful comments to improve this research.

