## Supplementary material for "Gene network module changes associated with the vertebrate fin to limb transition": Electronic supplementary material tables and figures

**Supplementary tables and figures**

**Supplementary Tables**

Supplementary Table S1. The statistics for anatomical entities and the extracted modules

|  | Pectoral fin | Pelvic fin | Forelimb | Hindlimb |
| --- | --- | --- | --- | --- |
| Number of genes with direct annotations to the anatomical entity | 192 | 13 | 216 | 530 |
| Number of genes annotated only to the parts of the anatomical entity | 3 | 2 | 44 | 239 |
| Number of genes annotated only to the bud of the anatomical entity | 3 | 0 | 7 | 8 |
| The total number of genes used for the module detection | 198 | 15 | 267 | 777 |
| The precision threshold used for candidate gene prediction | 0.7 | 0.05 | 0.7 | 0.7 |
| Number of predicted genes in the module | 45 | 605 | 18 | 32 |
| Number of genes with original annotations to the anatomical entity, a part, or the bud in the module | 175 | 12 | 225 | 639 |
| Total number of genes in the module | 220 | 617 | 243 | 671 |
| Number of genes with original annotations that were lost due to the network cutoff | 17 | 3 | 25 | 101 |
| Number of genes with original annotations that were lost due to isolation in the network | 6 | 0 | 17 | 37 |

Supplementary Table S2. The genes with original annotations that were lost due to network cutoff or isolation in the network

| Pectoral fin | Pelvic fin | Forelimb | Hindlimb |
| --- | --- | --- | --- |
| **Lost due to network cutoff: 17**   1. *bcl2l16* 2. *bot* 3. *don* 4. *ful* 5. *gpatch3* 6. *hmx4* 7. *mko* 8. *not found* 9. *si:ch211-185a18.2* 10. *tmem260* 11. *unm au21* 12. *unm m572* 13. *unm ti205* 14. *unm tm136* 15. *unm to219* 16. *wan* 17. *zon*   **Lost due to isolation: 6**   1. *ap1g1* 2. *dul* 3. *gaz* 4. *nokr2* 5. *nrg2a* 6. *perp* | **Lost due to network cutoff: 3**   1. *mir223* 2. *sub* 3. *wan*   **Lost due to isolation: 0** | **Lost due to network cutoff: 25**   1. *am* 2. *ccd* 3. *cl* 4. *dbf* 5. *del(2hoxd1-hoxd10)26ddu* 6. *del(6dlx6-dlx5)1tlu* 7. *etn3* 8. *fts* 9. *is(in8b2-8b3.1;6c1)1tshir* 10. *lgl* 11. *mdga2* 12. *mhdaali18* 13. *mirc1* 14. *morc2a* 15. *os* 16. *pf* 17. *t(7;18)50h* 18. *tg(cag-mrfp1,-sox9,-egfp)1haak* 19. *tg(col2a1-mef2c/vp16)1eno* 20. *tg(hand2)#tshir* 21. *tg(pgk1-fgf2)15cofn* 22. *tg(plp1-lmnb1)1108qsp* 23. *tg(prrx1-sox9,-lacz)1haak* 24. *tg(tyr,col2a1-trpv4*r594h)#dhco* 25. *vsd*   **Lost due to isolation: 17**   1. *btd* 2. *cdk20* 3. *cpox* 4. *del(5d5mit73-d5mit351)5jcs* 5. *lgi4* 6. *nkx6-1* 7. *npat* 8. *ostm1* 9. *pappa2* 10. *pex10* 11. *rnf165* 12. *scx* 13. *tg(sod1*g127x)716mrkl* 14. *tram2* 15. *trpv4* 16. *uchl1* 17. *vps54* | **Lost due to network cutoff: 101**   1. *4933430i17rik* 2. *ano5* 3. *aspb* 4. *b2b1594clo* 5. *b2b2187clo* 6. *bolt* 7. *cby* 8. *cl* 9. *clec11a* 10. *dbf* 11. *del(6dlx6-dlx5)1tlu* 12. *dh* 13. *dmpy* 14. *dp(16cbr1-fam3b)1rhr* 15. *fts* 16. *gnd* 17. *hacd1* 18. *hxd* 19. *hydro* 20. *igf1sl1* 21. *inad* 22. *is(in8b2-8b3.1;6c1)1tshir* 23. *klhl41* 24. *lgl* 25. *lx* 26. *lz* 27. *map3k20* 28. *mhdaali18* 29. *mir140* 30. *mir92-1* 31. *mirc1* 32. *mpc234h* 33. *nad* 34. *nma* 35. *nmf419* 36. *not found* 37. *ogfod1* 38. *os* 39. *pf* 40. *pl* 41. *pma* 42. *ps* 43. *scarf2* 44. *skc3* 45. *srn* 46. *ssq* 47. *t(7;18)50h* 48. *tal* 49. *tenm4* 50. *tg(acta1-il15*)11650lsq* 51. *tg(ar*100q)c25als* 52. *tg(ar*100q)c32als* 53. *tg(b19-rnai:il3)241ckn* 54. *tg(cag-dsred2/rnai:tardbp)6zxu* 55. *tg(cd4-npm/alk)n1ingh* 56. *tg(ckmm-cav3)1ysu* 57. *tg(cnp-gpr17)1qrlu* 58. *tg(col11a2-npr2*)28keoz* 59. *tg(col1a1-ifitm5*)1brle* 60. *tg(col1a1)73prc* 61. *tg(ctnnb1)1efu* 62. *tg(eef1a1-gnas*r201c)184pabi* 63. *tg(epo*)458mym* 64. *tg(gfap-il6)g167lms* 65. *tg(gfap-il6)g16lms* 66. *tg(gfap-il6)g369lms* 67. *tg(h2-k-fosl1)1wag* 68. *tg(igkv3-5*-myc)#plbe* 69. *tg(lckil4)1315dbl* 70. *tg(mbp-cadm4*)#pele* 71. *tg(mt1-hgfsf)19lmb* 72. *tg(mt2a-tgfbr2)4rser* 73. *tg(myh7-pln)2egk* 74. *tg(nefl*e397k)#milg* 75. *tg(pgk1-fgf2)15cofn* 76. *tg(plp1*)4rsj* 77. *tg(prnp-ar*112q)#deme* 78. *tg(prnp-fus)wt3cshw* 79. *tg(prnp-mapt*p301l)jnpl3hlmc* 80. *tg(prnp-snca*a53t)83vle* 81. *tg(prnp-tardbp*a315t)23jlel* 82. *tg(prnp)c35cwe* 83. *tg(prnp*)#rgab* 84. *tg(s100b-v-erbb)4496waw* 85. *tg(sod1*)df7yaw* 86. *tg(sod1*g37r)106dpr* 87. *tg(sod1*g37r)29dpr* 88. *tg(sod1*g85r)148dwc* 89. *tg(sod1*g85r)74dwc* 90. *tg(sod1*g93a)<dl>1gur* 91. *tg(sod1*g93a)1gur* 92. *tg(sod1*h46r)lara* 93. *tg(tcrar28,tcrbr28)krndim* 94. *tg(teto-htr4*d100a)7niss* 95. *tg(thy1-mapt*p301l)2vln* 96. *tg(thy1-mapt*p301s)2541godt* 97. *tg(thy1-snca*a30p)18pjk* 98. *tg(thy1-ubqln2*p497s)3mont* 99. *tg(thy1-ubqln2*p506t)6mont* 100. *ts(17<16>)65dn* 101. *vsd*   **Lost due to isolation: 37**   1. *adgrf5* 2. *akap11* 3. *arid5b* 4. *col4a3bp* 5. *coq9* 6. *ctsf* 7. *dnase1l2* 8. *elmod1* 9. *epg5* 10. *hhipl1* 11. *hr* 12. *lgi4* 13. *lncpint* 14. *ltn1* 15. *lyst* 16. *mbd5* 17. *noto* 18. *nxn* 19. *pex10* 20. *pgam5* 21. *pole4* 22. *psph* 23. *qk* 24. *rnf13* 25. *sacs* 26. *skida1* 27. *slc38a2* 28. *sno* 29. *spata6* 30. *spg20* 31. *stard10* 32. *tg(pmp22)c22clh* 33. *tg(prnp-fus*r521c)3313ejh* 34. *tg(sod1*g85r)#roos* 35. *tg(sod1*g86r)m3jwg* 36. *tg(thy1-mapt*)30schd* 37. *zfp106* |

Supplementary Table S3. The 45 predicted genes of the pectoral fin module ranked according to their weighted degrees.

| Zebrafish gene name | ZFIN identifier | Weighted degree | Rank in the module | Mouse ortholog name | Mouse ortholog status |
| --- | --- | --- | --- | --- | --- |
| *bmp4* | zdb-gene-980528-2059 | 35.5652 | 2 | *bmp4* | direct annotation to the forelimb |
| *bmp2b* | zdb-gene-980526-474 | 34.3109 | 3 | *bmp2* | direct annotation to the forelimb |
| *wnt3a* | zdb-gene-001106-1 | 31.4927 | 4 | *wnt3a* | predicted |
| *fgf8a* | zdb-gene-990415-72 | 31.0094 | 5 | *fgf8* | direct annotation to the forelimb |
| *gli2a* | zdb-gene-990706-8 | 27.2862 | 8 | *gli2* | direct annotation to the forelimb |
| *wnt5b* | zdb-gene-980526-87 | 25.9169 | 10 | *wnt5b* | not associated with the forelimb |
| *fgfr1a* | zdb-gene-980526-255 | 25.6442 | 12 | *fgfr1* | direct annotation to the forelimb |
| *smad5* | zdb-gene-990603-9 | 25.5690 | 13 | *smad5* | not associated with the forelimb |
| *gli1* | zdb-gene-030321-1 | 25.2858 | 14 | *gli1* | not associated with the forelimb |
| *tcf7l1a* | zdb-gene-980605-30 | 24.8101 | 15 | *tcf7l1* | not associated with the forelimb |
| *foxd3* | zdb-gene-980526-143 | 24.7462 | 16 | *foxd3* | not associated with the forelimb |
| *ta* | zdb-gene-980526-437 | 23.7992 | 18 | *NA* | not associated with the forelimb |
| *cdx4* | zdb-gene-980526-330 | 23.1331 | 20 | *cdx4* | not associated with the forelimb |
| *pax2a* | zdb-gene-990415-8 | 22.2653 | 22 | *pax2* | not associated with the forelimb |
| *ctnnb2* | zdb-gene-040426-2575 | 22.1606 | 24 | *NA* | not associated with the forelimb |
| *ptch2* | zdb-gene-980526-44 | 21.7001 | 25 | *ptch2* | not associated with the forelimb |
| *isl1* | zdb-gene-980526-112 | 21.5418 | 26 | *isl1* | not associated with the forelimb |
| *fgf3* | zdb-gene-980526-178 | 20.9767 | 28 | *fgf3* | not associated with the forelimb |
| *wnt4a* | zdb-gene-980526-352 | 20.4183 | 29 | *wnt4* | predicted |
| *gpc4* | zdb-gene-011119-1 | 18.5224 | 31 | *gpc4* | not associated with the forelimb |
| *ihha* | zdb-gene-051010-1 | 18.1064 | 32 | *ihh* | direct annotation to the forelimb |
| *wnt11* | zdb-gene-990603-12 | 17.1389 | 34 | *wnt11* | not associated with the forelimb |
| *zic2a* | zdb-gene-000710-4 | 15.2976 | 40 | *zic2* | annotated to a part or bud |
| *nkx2.2a* | zdb-gene-980526-403 | 15.0866 | 41 | *nkx2-2* | not associated with the forelimb |
| *dlx2a* | zdb-gene-980526-212 | 14.8123 | 43 | *dlx2* | not associated with the forelimb |
| *pitx2* | zdb-gene-990714-27 | 14.7898 | 44 | *pitx2* | not associated with the forelimb |
| *msx1a* | zdb-gene-980526-312 | 14.6308 | 45 | *msx1* | predicted |
| *myf5* | zdb-gene-000616-6 | 13.7077 | 48 | *myf5* | not associated with the forelimb |
| *esr2a* | zdb-gene-030116-2 | 13.2000 | 51 | *esr2* | direct annotation to the forelimb |
| *dharma* | zdb-gene-990415-22 | 9.6753 | 59 | *NA* | not associated with the forelimb |
| *tfap2a* | zdb-gene-011212-6 | 9.2007 | 62 | *tfap2a* | direct annotation to the forelimb |
| *mecom* | NA | 8.4715 | 89 | *mecom* | annotated to a part or bud |
| *sox9b* | zdb-gene-001103-2 | 7.2558 | 104 | *sox9* | direct annotation to the forelimb |
| *grem2b* | zdb-gene-030911-9 | 6.8164 | 108 | *grem2* | not associated with the forelimb |
| *unm t31148* | zdb-gene-070117-1894 | 6.6863 | 109 | *NA* | not associated with the forelimb |
| *dzip1* | zdb-gene-040526-1 | 6.4409 | 112 | *dzip1* | not associated with the forelimb |
| *b3gat3* | zdb-gene-020419-3 | 6.0316 | 115 | *NA* | not associated with the forelimb |
| *scube2* | zdb-gene-050302-80 | 5.3349 | 123 | *scube2* | not associated with the forelimb |
| *hot* | zdb-gene-070117-2108 | 2.7511 | 160 | *NA* | not associated with the forelimb |
| *frem2b* | zdb-gene-081119-4 | 2.6947 | 163 | *frem2* | not associated with the forelimb |
| *unm s273* | zdb-gene-070117-864 | 2.4459 | 167 | *NA* | not associated with the forelimb |
| *unm s245* | zdb-gene-070117-865 | 2.4459 | 167 | *NA* | not associated with the forelimb |
| *eda* | zdb-gene-050107-6 | 2.3936 | 169 | *eda* | not associated with the forelimb |
| *zmp:0000001138* | zdb-gene-140106-98 | 2.1176 | 172 | *NA* | not associated with the forelimb |
| *mgt* | zdb-gene-070117-2188 | 1.0103 | 197 | *NA* | not associated with the forelimb |

*NA indicates ‘not available’ due to the ortholog not found in the mouse

Supplementary Table S4. The top 50 predicted genes of the pelvic fin module ranked according to their weighted degrees. The full predicted gene list is available on electronic material, supplementary file S9.

| Zebrafish gene name | ZFIN identifier | Weighted degree | Rank in the module | Mouse ortholog name | Mouse ortholog status |
| --- | --- | --- | --- | --- | --- |
| *hsp90ab1* | zdb-gene-990415-95 | 129.9415 | 1 | *hsp90ab1* | not associated with the hindlimb |
| *mapk3* | zdb-gene-040121-1 | 121.5328 | 2 | *mapk3* | not associated with the hindlimb |
| *rhoab* | zdb-gene-040322-2 | 108.0837 | 3 | *rhoa* | not associated with the hindlimb |
| *ctnnb1* | zdb-gene-980526-362 | 106.2749 | 4 | *ctnnb1* | direct annotation to the hindlimb |
| *hsp90aa1.2* | zdb-gene-031001-3 | 105.1273 | 5 | *hsp90aa1* | not associated with the hindlimb |
| *paics* | zdb-gene-030131-9762 | 104.4687 | 6 | *paics* | not associated with the hindlimb |
| *gsk3b* | zdb-gene-990714-4 | 101.2293 | 7 | *gsk3b* | not associated with the hindlimb |
| *cad* | zdb-gene-021030-4 | 97.8373 | 8 | *cad* | not associated with the hindlimb |
| *cdc42* | zdb-gene-030131-8783 | 97.7752 | 9 | *NA* | not associated with the hindlimb |
| *acta1b* | zdb-gene-030131-55 | 97.6682 | 10 | *acta1* | not associated with the hindlimb |
| *smarca4a* | zdb-gene-030605-1 | 97.4336 | 11 | *smarca4* | direct annotation to the hindlimb |
| *mapk1* | zdb-gene-030722-2 | 96.3956 | 12 | *mapk1* | not associated with the hindlimb |
| *jupa* | zdb-gene-991207-22 | 95.0796 | 13 | *jup* | not associated with the hindlimb |
| *cdk1* | zdb-gene-010320-1 | 93.9169 | 14 | *cdk1* | not associated with the hindlimb |
| *kras* | NA | 93.4030 | 15 | *kras* | predicted |
| *rac1a* | zdb-gene-030131-5415 | 92.9107 | 16 | *rac1* | direct annotation to the hindlimb |
| *mapk14a* | zdb-gene-010202-2 | 92.1789 | 17 | *mapk14* | direct annotation to the hindlimb |
| *src* | zdb-gene-030131-3809 | 91.7256 | 18 | *src* | direct annotation to the hindlimb |
| *fgfr1a* | zdb-gene-980526-255 | 91.2553 | 19 | *fgfr1* | direct annotation to the hindlimb |
| *met* | zdb-gene-041014-1 | 90.8735 | 20 | *met* | direct annotation to the hindlimb |
| *insrb* | zdb-gene-020503-4 | 90.6827 | 21 | *insr* | not associated with the hindlimb |
| *si:ch211-163m16.1* | NA | 84.6841 | 22 | *NA* | not associated with the hindlimb |
| *actl6a* | zdb-gene-020419-36 | 83.9023 | 23 | *actl6a* | not associated with the hindlimb |
| *tp53* | zdb-gene-990415-270 | 83.8035 | 24 | *trp53* | direct annotation to the hindlimb |
| *pak2a* | zdb-gene-021011-2 | 83.2637 | 25 | *pak2* | not associated with the hindlimb |
| *ehmt2* | zdb-gene-010501-6 | 82.9897 | 26 | *ehmt2* | not associated with the hindlimb |
| *kdrl* | zdb-gene-000705-1 | 81.0991 | 27 | *NA* | not associated with the hindlimb |
| *mtor* | zdb-gene-030131-2974 | 80.8121 | 28 | *mtor* | annotated to a part or bud |
| *pkn2* | zdb-gene-061207-42 | 79.5943 | 29 | *pkn2* | not associated with the hindlimb |
| *prkcbb* | zdb-gene-040426-1178 | 79.5529 | 30 | *prkcb* | not associated with the hindlimb |
| *hsp90aa1.1* | zdb-gene-990415-94 | 78.6325 | 31 | *hsp90aa1* | not associated with the hindlimb |
| *ptenb* | zdb-gene-030616-47 | 78.5310 | 32 | *pten* | annotated to a part or bud |
| *kita* | zdb-gene-980526-464 | 77.2229 | 33 | *kit* | not associated with the hindlimb |
| *akt2* | zdb-gene-031007-5 | 77.0797 | 34 | *akt2* | not associated with the hindlimb |
| *igf1ra* | zdb-gene-020503-1 | 76.5270 | 35 | *igf1r* | not associated with the hindlimb |
| *bmp4* | zdb-gene-980528-2059 | 76.2150 | 36 | *bmp4* | direct annotation to the hindlimb |
| *igf1rb* | zdb-gene-020503-2 | 75.0886 | 37 | *igf1r* | not associated with the hindlimb |
| *rap1b* | zdb-gene-030131-9662 | 73.3502 | 38 | *rap1b* | not associated with the hindlimb |
| *rac2* | zdb-gene-040625-27 | 73.3136 | 39 | *rac2* | not associated with the hindlimb |
| *hspa9* | zdb-gene-030828-12 | 72.3324 | 40 | *hspa9* | not associated with the hindlimb |
| *hdac1* | zdb-gene-020419-32 | 69.4428 | 41 | *hdac1* | not associated with the hindlimb |
| *pola1* | zdb-gene-030114-9 | 69.0229 | 42 | *pola1* | not associated with the hindlimb |
| *actc1a* | zdb-gene-040520-4 | 68.6027 | 43 | *actc1* | not associated with the hindlimb |
| *top2b* | zdb-gene-041008-136 | 68.4413 | 44 | *top2b* | not associated with the hindlimb |
| *bmp2b* | zdb-gene-980526-474 | 68.2429 | 45 | *bmp2* | predicted |
| *insra* | zdb-gene-020503-3 | 68.1565 | 46 | *insr* | not associated with the hindlimb |
| *flt1* | zdb-gene-050407-1 | 68.0761 | 47 | *flt1* | not associated with the hindlimb |
| *ralbb* | zdb-gene-040625-121 | 67.8323 | 48 | *ralb* | not associated with the hindlimb |
| *btk* | zdb-gene-070531-1 | 67.4762 | 49 | *NA* | not associated with the hindlimb |
| *rac1b* | zdb-gene-060312-45 | 67.0359 | 50 | *rac1* | direct annotation to the hindlimb |

*NA indicates ‘not available’ due to the ortholog not found in the mouse

Supplementary Table S5. The 18 predicted genes of the forelimb module ranked according to their weighted degrees.

| Mouse gene name | MGI identifier | Weighted degree | Rank in the module | Zebrafish ortholog names | Zebrafish ortholog status |
| --- | --- | --- | --- | --- | --- |
| *smad4* | mgi:894293 | 34.5294 | 6 | *smad4a, smad4b* | not associated with the pectoral fin |
| *bmp7* | mgi:103302 | 34.2132 | 7 | *bmp7a* | direct annotation to the pectoral fin |
| *wnt3a* | mgi:98956 | 30.9124 | 14 | *wnt3a* | predicted |
| *nog* | mgi:104327 | 29.3757 | 19 | *NA* | not associated with the pectoral fin |
| *wnt4* | mgi:98957 | 27.9668 | 23 | *wnt4a* | predicted |
| *ptch1* | mgi:105373 | 27.3374 | 24 | *ptch1* | direct annotation to the pectoral fin |
| *wnt1* | mgi:98953 | 26.4361 | 25 | *wnt1* | not associated with the pectoral fin |
| *bmpr1a* | mgi:1338938 | 26.0359 | 26 | *bmpr1ab, bmpr1aa* | not associated with the pectoral fin |
| *chrd* | mgi:1313268 | 24.6081 | 28 | *chrd* | not associated with the pectoral fin |
| *msx1* | mgi:97168 | 23.4172 | 31 | *msx1a* | predicted |
| *msx2* | mgi:97169 | 22.8361 | 32 | *msx2b, msx2a* | not associated with the pectoral fin |
| *fst* | mgi:95586 | 18.1062 | 46 | *fsta, fstb* | not associated with the pectoral fin |
| *wnt7b* | mgi:98962 | 17.7777 | 47 | *wnt7ba, wnt7bb* | not associated with the pectoral fin |
| *dlx5* | mgi:101926 | 16.1115 | 51 | *dlx5a* | annotated to a part or bud |
| *wnt9a* | mgi:2446084 | 15.9927 | 52 | *wnt9a* | not associated with the pectoral fin |
| *foxc2* | mgi:1347481 | 13.9395 | 61 | *NA* | not associated with the pectoral fin |
| *nkx3-2* | mgi:108015 | 13.6063 | 63 | *nkx3.2* | not associated with the pectoral fin |
| *tbx4* | mgi:102556 | 8.7400 | 84 | *tbx4* | not associated with the pectoral fin |

*NA indicates ‘not available’ due to the ortholog not found in the zebrafish

Supplementary Table S6. The 32 predicted genes of the hindlimb module ranked according to their weighted degrees.

| Mouse gene name | MGI identifier | Weighted degree | Rank in the module | Zebrafish ortholog names | Zebrafish ortholog status |
| --- | --- | --- | --- | --- | --- |
| *hras* | mgi:96224 | 72.8342 | 2 | *hrasb* | predicted |
| *kras* | mgi:96680 | 70.7748 | 5 | *kras* | predicted |
| *myc* | mgi:97250 | 66.8072 | 7 | *mycb, myca* | not associated with the pelvic fin |
| *fos* | mgi:95574 | 64.7904 | 9 | *fosab, fosaa* | not associated with the pelvic fin |
| *tgfb1* | mgi:98725 | 54.2763 | 15 | *tgfb1b, tgfb1a* | not associated with the pelvic fin |
| *igf1* | mgi:96432 | 52.9341 | 17 | *igf1* | not associated with the pelvic fin |
| *fgf2* | mgi:95516 | 51.2651 | 19 | *fgf2* | not associated with the pelvic fin |
| *casp3* | mgi:107739 | 49.0774 | 23 | *casp3a* | predicted |
| *bmp7* | mgi:103302 | 48.2316 | 26 | *bmp7a* | predicted |
| *wnt5a* | mgi:98958 | 48.1588 | 27 | *wnt5a* | not associated with the pelvic fin |
| *smad2* | mgi:108051 | 45.6514 | 30 | *smad2* | predicted |
| *pdgfra* | mgi:97530 | 45.2074 | 31 | *pdgfra* | not associated with the pelvic fin |
| *bmp2* | mgi:88177 | 45.0361 | 32 | *bmp2a, bmp2b* | predicted |
| *nog* | mgi:104327 | 39.7063 | 46 | *NA* | not associated with the pelvic fin |
| *ptch1* | mgi:105373 | 38.8191 | 48 | *ptch1* | predicted |
| *lef1* | mgi:96770 | 36.7074 | 53 | *lef1* | predicted |
| *wnt1* | mgi:98953 | 35.9468 | 60 | *wnt1* | not associated with the pelvic fin |
| *wnt4* | mgi:98957 | 35.4014 | 62 | *wnt4a* | predicted |
| *mmp2* | mgi:97009 | 35.3982 | 63 | *mmp2* | not associated with the pelvic fin |
| *spp1* | mgi:98389 | 33.6944 | 70 | *NA* | not associated with the pelvic fin |
| *dcn* | mgi:94872 | 33.0020 | 73 | *dcn* | not associated with the pelvic fin |
| *mmp14* | mgi:101900 | 30.9303 | 77 | *mmp14a, mmp14b* | not associated with the pelvic fin |
| *chrd* | mgi:1313268 | 29.8136 | 81 | *chrd* | not associated with the pelvic fin |
| *gja1* | mgi:95713 | 28.3709 | 84 | *cx43* | predicted |
| *msx1* | mgi:97168 | 26.6468 | 88 | *msx1a* | predicted |
| *twist1* | mgi:98872 | 25.5266 | 93 | *twist1a, twist1b* | not associated with the pelvic fin |
| *bglap* | mgi:88156 | 24.8621 | 99 | *NA* | not associated with the pelvic fin |
| *bglap2* | mgi:88157 | 24.8447 | 100 | *NA* | not associated with the pelvic fin |
| *sp7* | mgi:2153568 | 21.0093 | 119 | *sp7* | not associated with the pelvic fin |
| *foxc2* | mgi:1347481 | 17.4167 | 192 | *NA* | not associated with the pelvic fin |
| *nkx3-2* | mgi:108015 | 15.3314 | 207 | *nkx3.2* | not associated with the pelvic fin |
| *hapln1* | mgi:1337006 | 8.9023 | 279 | *hapln1a, hapln1b* | not associated with the pelvic fin |

*NA indicates ‘not available’ due to the ortholog not found in the zebrafish

Supplementary Table S7. The enriched Biological Process terms from the Gene Ontology that are common to the predicted genes and genes with original annotations for the pectoral fin. The enriched terms are sorted based on the p-value of those terms for the predicted genes.

| Term identifier | Term name | P-value for predicted genes | P-value for original genes |
| --- | --- | --- | --- |
| GO:0007275 | multicellular organism development | 2.41E-12 | 1.85E-09 |
| GO:0009953 | dorsal/ventral pattern formation | 6.32E-12 | 2.23E-06 |
| GO:0051216 | cartilage development | 2.28E-11 | 4.70E-17 |
| GO:0009880 | embryonic pattern specification | 4.19E-09 | 0.0160149 |
| GO:0006355 | regulation of transcription, DNA-templated | 4.82E-09 | 1.70E-07 |
| GO:0014032 | neural crest cell development | 2.91E-08 | 0.00016897 |
| GO:0001756 | somitogenesis | 6.37E-08 | 2.80E-07 |
| GO:0007368 | determination of left/right symmetry | 1.71E-07 | 4.78E-09 |
| GO:0030182 | neuron differentiation | 3.70E-07 | 9.35E-05 |
| GO:0007224 | smoothened signaling pathway | 6.03E-07 | 0.00117078 |
| GO:0048703 | embryonic viscerocranium morphogenesis | 3.24E-06 | 2.95E-16 |
| GO:2000223 | regulation of BMP signaling pathway involved in heart jogging | 3.31E-06 | 0.0037629 |
| GO:0042476 | odontogenesis | 4.29E-06 | 2.35E-06 |
| GO:0021984 | adenohypophysis development | 4.29E-06 | 0.00442453 |
| GO:0030902 | hindbrain development | 4.52E-06 | 4.34E-08 |
| GO:0009952 | anterior/posterior pattern specification | 7.59E-06 | 0.00060755 |
| GO:0001947 | heart looping | 8.61E-06 | 6.61E-15 |
| GO:0016055 | Wnt signaling pathway | 1.66E-05 | 6.53E-06 |
| GO:0048793 | pronephros development | 2.04E-05 | 1.01E-05 |
| GO:0003143 | embryonic heart tube morphogenesis | 3.81E-05 | 3.94E-08 |
| GO:0031018 | endocrine pancreas development | 5.91E-05 | 0.02264448 |
| GO:0042694 | muscle cell fate specification | 9.42E-05 | 0.04578302 |
| GO:0031290 | retinal ganglion cell axon guidance | 0.000111737 | 9.33E-06 |
| GO:0048701 | embryonic cranial skeleton morphogenesis | 0.000292195 | 5.06E-09 |
| GO:0060070 | canonical Wnt signaling pathway | 0.00042765 | 8.61E-05 |
| GO:0021508 | floor plate formation | 0.000564411 | 0.00513581 |
| GO:0048264 | determination of ventral identity | 0.001287915 | 0.00055075 |
| GO:0060037 | pharyngeal system development | 0.001287915 | 0.00055075 |
| GO:0060041 | retina development in camera-type eye | 0.001442926 | 5.42E-06 |
| GO:0071599 | otic vesicle development | 0.001684552 | 0.0148082 |
| GO:0030917 | midbrain-hindbrain boundary development | 0.002132019 | 0.00117078 |
| GO:0060059 | embryonic retina morphogenesis in camera-type eye | 0.002132019 | 0.01854958 |
| GO:0045893 | positive regulation of transcription, DNA-templated | 0.002141963 | 0.04413722 |
| GO:0008543 | fibroblast growth factor receptor signaling pathway | 0.003568028 | 0.03020605 |
| GO:0031016 | pancreas development | 0.003981011 | 9.33E-06 |
| GO:0001501 | skeletal system development | 0.003981011 | 0.00293934 |
| GO:0043010 | camera-type eye development | 0.007171621 | 0.00060755 |
| GO:0007507 | heart development | 0.010543118 | 1.44E-08 |
| GO:0003342 | proepicardium development | 0.012678336 | 0.00059083 |
| GO:0010002 | cardioblast differentiation | 0.022708189 | 3.72E-05 |
| GO:0045892 | negative regulation of transcription, DNA-templated | 0.024331089 | 0.00642797 |
| GO:0030166 | proteoglycan biosynthetic process | 0.025200109 | 0.00259184 |
| GO:0021986 | habenula development | 0.032638807 | 0.00442453 |
| GO:0048384 | retinoic acid receptor signaling pathway | 0.035106069 | 0.00015684 |
| GO:0001649 | osteoblast differentiation | 0.035106069 | 0.00015684 |
| GO:0048709 | oligodendrocyte differentiation | 0.035106069 | 3.28E-06 |
| GO:0030198 | extracellular matrix organization | 0.047350757 | 0.00940935 |

Supplementary Table S8. The enriched Biological Process terms from the Gene Ontology that are common to the predicted genes and genes with original annotations for the pelvic fin. The enriched terms are sorted based on the p-value of those terms for the predicted genes.

| Term identifier | Term name | P-value for the predicted genes | P-value for the original genes |
| --- | --- | --- | --- |
| GO:0033333 | fin development | 7.11E-10 | 9.72E-08 |

Supplementary Table S9. The enriched Biological Process terms from the Gene Ontology that are common to the predicted genes and genes with original annotations for the forelimb. The enriched terms are sorted based on the p-value of those terms for the predicted genes.

| Term identifier | Term name | P-value for the predicted genes | P-value for the original genes |
| --- | --- | --- | --- |
| GO:0030509 | BMP signaling pathway | 2.12E-17 | 2.19E-10 |
| GO:0007275 | multicellular organism development | 5.15E-12 | 5.81E-25 |
| GO:0007389 | pattern specification process | 1.71E-09 | 1.90E-15 |
| GO:0030326 | embryonic limb morphogenesis | 2.62E-09 | 1.44E-40 |
| GO:0045669 | positive regulation of osteoblast differentiation | 3.87E-09 | 2.15E-10 |
| GO:0045893 | positive regulation of transcription, DNA-templated | 1.91E-08 | 8.27E-22 |
| GO:0001701 | in utero embryonic development | 2.06E-07 | 3.67E-14 |
| GO:0042475 | odontogenesis of dentin-containing tooth | 3.08E-07 | 2.14E-16 |
| GO:0042733 | embryonic digit morphogenesis | 3.72E-07 | 6.50E-38 |
| GO:0051216 | cartilage development | 8.94E-07 | 1.05E-21 |
| GO:0060021 | palate development | 1.03E-06 | 1.88E-18 |
| GO:0060070 | canonical Wnt signaling pathway | 1.24E-06 | 1.80E-11 |
| GO:0045944 | positive regulation of transcription from RNA polymerase II promoter | 1.27E-06 | 5.94E-32 |
| GO:0000122 | negative regulation of transcription from RNA polymerase II promoter | 2.30E-06 | 3.42E-23 |
| GO:0001501 | skeletal system development | 2.60E-06 | 9.14E-36 |
| GO:0009952 | anterior/posterior pattern specification | 3.12E-06 | 1.71E-17 |
| GO:0001649 | osteoblast differentiation | 3.23E-06 | 3.89E-10 |
| GO:0001707 | mesoderm formation | 4.43E-06 | 4.00E-05 |
| GO:0071773 | cellular response to BMP stimulus | 4.43E-06 | 4.00E-05 |
| GO:0007411 | axon guidance | 9.70E-06 | 1.08E-07 |
| GO:0001658 | branching involved in ureteric bud morphogenesis | 1.16E-05 | 3.84E-12 |
| GO:0009953 | dorsal/ventral pattern formation | 1.24E-05 | 1.42E-21 |
| GO:0008285 | negative regulation of cell proliferation | 2.11E-05 | 1.63E-13 |
| GO:0003007 | heart morphogenesis | 2.78E-05 | 2.07E-09 |
| GO:0090263 | positive regulation of canonical Wnt signaling pathway | 3.19E-05 | 8.99E-12 |
| GO:0016055 | Wnt signaling pathway | 3.95E-05 | 2.85E-11 |
| GO:0001503 | ossification | 9.34E-05 | 1.17E-21 |
| GO:0090090 | negative regulation of canonical Wnt signaling pathway | 0.000111887 | 1.05E-10 |
| GO:0009887 | organ morphogenesis | 0.000136246 | 1.77E-11 |
| GO:0045892 | negative regulation of transcription, DNA-templated | 0.00014868 | 3.08E-08 |
| GO:0045596 | negative regulation of cell differentiation | 0.000155607 | 0.00172739 |
| GO:0048646 | anatomical structure formation involved in morphogenesis | 0.000226843 | 5.78E-06 |
| GO:0042474 | middle ear morphogenesis | 0.000226843 | 5.78E-06 |
| GO:0003148 | outflow tract septum morphogenesis | 0.000226843 | 0.000132268 |
| GO:0001822 | kidney development | 0.000229367 | 1.74E-17 |
| GO:0021983 | pituitary gland development | 0.00035634 | 1.84E-05 |
| GO:0001837 | epithelial to mesenchymal transition | 0.000380704 | 0.00036855 |
| GO:0060349 | bone morphogenesis | 0.00040586 | 4.87E-11 |
| GO:0035115 | embryonic forelimb morphogenesis | 0.000514369 | 6.35E-37 |
| GO:0030501 | positive regulation of bone mineralization | 0.000543461 | 1.62E-13 |
| GO:0060325 | face morphogenesis | 0.000635432 | 7.74E-05 |
| GO:0048568 | embryonic organ development | 0.000635432 | 4.91E-06 |
| GO:0007492 | endoderm development | 0.000635432 | 0.000990141 |
| GO:0030901 | midbrain development | 0.000635432 | 1.15E-08 |
| GO:0030514 | negative regulation of BMP signaling pathway | 0.000840378 | 0.000152854 |
| GO:0010862 | positive regulation of pathway-restricted SMAD protein phosphorylation | 0.000914879 | 1.46E-05 |
| GO:0001657 | ureteric bud development | 0.000914879 | 9.46E-07 |
| GO:0045599 | negative regulation of fat cell differentiation | 0.000914879 | 0.001967343 |
| GO:0003151 | outflow tract morphogenesis | 0.001156789 | 7.48E-09 |
| GO:0045668 | negative regulation of osteoblast differentiation | 0.001243528 | 0.00346626 |
| GO:0001756 | somitogenesis | 0.001243528 | 1.82E-07 |
| GO:0045165 | cell fate commitment | 0.002045776 | 2.19E-11 |
| GO:0042472 | inner ear morphogenesis | 0.002045776 | 1.01E-07 |
| GO:0050679 | positive regulation of epithelial cell proliferation | 0.002276056 | 1.05E-13 |
| GO:0006355 | regulation of transcription, DNA-templated | 0.0030741 | 1.70E-09 |
| GO:0007267 | cell-cell signaling | 0.004133083 | 1.62E-09 |
| GO:0030336 | negative regulation of cell migration | 0.004371326 | 0.030620272 |
| GO:0006351 | transcription, DNA-templated | 0.005714263 | 2.85E-08 |
| GO:0030324 | lung development | 0.005931081 | 6.63E-18 |
| GO:0048762 | mesenchymal cell differentiation | 0.010296133 | 0.00020907 |
| GO:0060272 | embryonic skeletal joint morphogenesis | 0.01122718 | 1.20E-07 |
| GO:0043616 | keratinocyte proliferation | 0.012157404 | 0.008847589 |
| GO:0061053 | somite development | 0.013086804 | 0.00045 |
| GO:0008284 | positive regulation of cell proliferation | 0.013313841 | 3.23E-18 |
| GO:0014032 | neural crest cell development | 0.015870067 | 8.96E-07 |
| GO:0003203 | endocardial cushion morphogenesis | 0.015870067 | 3.15E-05 |
| GO:0035137 | hindlimb morphogenesis | 0.015870067 | 1.96E-08 |
| GO:0032331 | negative regulation of chondrocyte differentiation | 0.016796179 | 2.91E-08 |
| GO:0051145 | smooth muscle cell differentiation | 0.016796179 | 4.01E-05 |
| GO:0060037 | pharyngeal system development | 0.016796179 | 0.016734602 |
| GO:0048856 | anatomical structure development | 0.016796179 | 4.01E-05 |
| GO:0021904 | dorsal/ventral neural tube patterning | 0.017721471 | 4.21E-08 |
| GO:0008283 | cell proliferation | 0.017771413 | 3.67E-05 |
| GO:0045879 | negative regulation of smoothened signaling pathway | 0.019569595 | 7.65E-05 |
| GO:0023019 | signal transduction involved in regulation of gene expression | 0.021414447 | 0.002033958 |
| GO:0060425 | lung morphogenesis | 0.022335647 | 0.002305628 |
| GO:0007498 | mesoderm development | 0.0241756 | 0.002913859 |
| GO:0007507 | heart development | 0.024473032 | 1.18E-18 |
| GO:0042476 | odontogenesis | 0.025094353 | 4.27E-07 |
| GO:0035108 | limb morphogenesis | 0.026012293 | 6.57E-17 |
| GO:0001702 | gastrulation with mouth forming second | 0.026012293 | 0.038461084 |
| GO:0071542 | dopaminergic neuron differentiation | 0.026929419 | 7.12E-10 |
| GO:0048663 | neuron fate commitment | 0.029675922 | 2.55E-05 |
| GO:0035116 | embryonic hindlimb morphogenesis | 0.030589801 | 1.87E-19 |
| GO:0048589 | developmental growth | 0.032415128 | 1.19E-10 |
| GO:0034504 | protein localization to nucleus | 0.032415128 | 0.000592382 |
| GO:0048754 | branching morphogenesis of an epithelial tube | 0.033326579 | 4.55E-12 |
| GO:0002053 | positive regulation of mesenchymal cell proliferation | 0.034237221 | 1.34E-18 |
| GO:0030154 | cell differentiation | 0.034609309 | 1.49E-12 |
| GO:0060412 | ventricular septum morphogenesis | 0.035147055 | 0.008576143 |
| GO:0030879 | mammary gland development | 0.035147055 | 6.02E-05 |
| GO:0001656 | metanephros development | 0.035147055 | 7.45E-09 |
| GO:0033077 | T cell differentiation in thymus | 0.036056083 | 0.000898871 |
| GO:0048706 | embryonic skeletal system development | 0.040589145 | 4.11E-11 |
| GO:0045597 | positive regulation of cell differentiation | 0.042396746 | 1.13E-05 |
| GO:0048705 | skeletal system morphogenesis | 0.049595145 | 1.36E-07 |

Supplementary Table S10. The enriched Biological Process terms from the Gene Ontology that are common to the predicted genes and genes with original annotations for the hindlimb. The enriched terms are sorted based on the p-value of those terms for the predicted genes.

| Term identifier | Term name | P-value for the predicted genes | P-value for the original genes |
| --- | --- | --- | --- |
| GO:0045944 | positive regulation of transcription from RNA polymerase II promoter | 4.34E-14 | 7.62E-37 |
| GO:0045893 | positive regulation of transcription, DNA-templated | 3.71E-12 | 4.70E-31 |
| GO:0001649 | osteoblast differentiation | 1.27E-11 | 5.39E-12 |
| GO:0050679 | positive regulation of epithelial cell proliferation | 4.24E-11 | 2.50E-09 |
| GO:0001837 | epithelial to mesenchymal transition | 1.74E-09 | 0.00157941 |
| GO:0060325 | face morphogenesis | 6.66E-09 | 0.00497598 |
| GO:0008285 | negative regulation of cell proliferation | 1.07E-08 | 1.18E-17 |
| GO:0010629 | negative regulation of gene expression | 1.13E-08 | 7.93E-09 |
| GO:0008284 | positive regulation of cell proliferation | 1.36E-08 | 8.36E-26 |
| GO:0010628 | positive regulation of gene expression | 1.49E-08 | 5.42E-17 |
| GO:0001658 | branching involved in ureteric bud morphogenesis | 1.72E-08 | 6.90E-07 |
| GO:0009887 | organ morphogenesis | 2.72E-08 | 2.99E-14 |
| GO:0001701 | in utero embryonic development | 2.89E-08 | 3.27E-14 |
| GO:0030335 | positive regulation of cell migration | 4.24E-08 | 1.74E-05 |
| GO:0030326 | embryonic limb morphogenesis | 6.92E-08 | 9.15E-40 |
| GO:0045669 | positive regulation of osteoblast differentiation | 1.02E-07 | 8.74E-17 |
| GO:0007507 | heart development | 2.36E-07 | 8.32E-18 |
| GO:0060021 | palate development | 3.14E-07 | 8.09E-23 |
| GO:0010718 | positive regulation of epithelial to mesenchymal transition | 3.16E-07 | 0.01455875 |
| GO:0030509 | BMP signaling pathway | 3.53E-07 | 2.76E-12 |
| GO:0045892 | negative regulation of transcription, DNA-templated | 3.57E-07 | 4.21E-07 |
| GO:0042060 | wound healing | 5.21E-07 | 1.23E-05 |
| GO:0048701 | embryonic cranial skeleton morphogenesis | 5.57E-07 | 8.42E-08 |
| GO:0007275 | multicellular organism development | 5.67E-07 | 1.89E-21 |
| GO:0001503 | ossification | 5.78E-07 | 1.14E-28 |
| GO:0001934 | positive regulation of protein phosphorylation | 6.10E-07 | 1.08E-08 |
| GO:0001501 | skeletal system development | 9.93E-07 | 4.60E-43 |
| GO:0001657 | ureteric bud development | 1.30E-06 | 0.00213229 |
| GO:0030324 | lung development | 2.06E-06 | 3.43E-11 |
| GO:0000122 | negative regulation of transcription from RNA polymerase II promoter | 2.43E-06 | 7.84E-20 |
| GO:0007389 | pattern specification process | 2.81E-06 | 4.16E-11 |
| GO:0042475 | odontogenesis of dentin-containing tooth | 3.92E-06 | 6.87E-14 |
| GO:0042733 | embryonic digit morphogenesis | 4.73E-06 | 5.30E-25 |
| GO:0030500 | regulation of bone mineralization | 5.94E-06 | 1.79E-05 |
| GO:0045165 | cell fate commitment | 6.70E-06 | 6.90E-07 |
| GO:0060395 | SMAD protein signal transduction | 8.76E-06 | 9.49E-06 |
| GO:0042474 | middle ear morphogenesis | 9.01E-06 | 0.00415561 |
| GO:0051216 | cartilage development | 1.13E-05 | 6.37E-26 |
| GO:0060070 | canonical Wnt signaling pathway | 1.56E-05 | 3.09E-09 |
| GO:0001843 | neural tube closure | 2.19E-05 | 3.09E-06 |
| GO:0090090 | negative regulation of canonical Wnt signaling pathway | 2.67E-05 | 4.37E-10 |
| GO:0071773 | cellular response to BMP stimulus | 2.88E-05 | 0.00038339 |
| GO:0001707 | mesoderm formation | 2.88E-05 | 4.37E-06 |
| GO:0048754 | branching morphogenesis of an epithelial tube | 3.14E-05 | 5.48E-06 |
| GO:0030501 | positive regulation of bone mineralization | 3.41E-05 | 3.34E-09 |
| GO:0030879 | mammary gland development | 3.70E-05 | 7.76E-05 |
| GO:0043066 | negative regulation of apoptotic process | 3.88E-05 | 3.04E-15 |
| GO:0045596 | negative regulation of cell differentiation | 4.13E-05 | 0.01495266 |
| GO:0030514 | negative regulation of BMP signaling pathway | 6.59E-05 | 0.03947855 |
| GO:0048705 | skeletal system morphogenesis | 0.00010666 | 2.20E-09 |
| GO:0043406 | positive regulation of MAP kinase activity | 0.00010666 | 0.00073993 |
| GO:0045668 | negative regulation of osteoblast differentiation | 0.00011893 | 0.00015728 |
| GO:0048762 | mesenchymal cell differentiation | 0.00015495 | 0.00017231 |
| GO:0009612 | response to mechanical stimulus | 0.00016899 | 6.14E-05 |
| GO:0061053 | somite development | 0.00025555 | 0.00048879 |
| GO:0007179 | transforming growth factor beta receptor signaling pathway | 0.00028339 | 1.47E-07 |
| GO:0070374 | positive regulation of ERK1 and ERK2 cascade | 0.00028588 | 3.19E-09 |
| GO:0016477 | cell migration | 0.00030363 | 1.90E-06 |
| GO:2000679 | positive regulation of transcription regulatory region DNA binding | 0.00033627 | 0.00934741 |
| GO:0003203 | endocardial cushion morphogenesis | 0.0003807 | 0.00108701 |
| GO:0016055 | Wnt signaling pathway | 0.00045893 | 1.58E-09 |
| GO:0008283 | cell proliferation | 0.00051842 | 4.12E-06 |
| GO:1902895 | positive regulation of pri-miRNA transcription from RNA polymerase II promoter | 0.00064319 | 0.02270901 |
| GO:0032355 | response to estradiol | 0.00067863 | 4.88E-06 |
| GO:0023019 | signal transduction involved in regulation of gene expression | 0.0007037 | 0.0035404 |
| GO:0007267 | cell-cell signaling | 0.00071851 | 1.79E-07 |
| GO:0031214 | biomineral tissue development | 0.00112204 | 0.00012938 |
| GO:0001822 | kidney development | 0.00140877 | 2.39E-08 |
| GO:0031016 | pancreas development | 0.00145299 | 2.71E-06 |
| GO:0040007 | growth | 0.00172815 | 5.48E-06 |
| GO:0002053 | positive regulation of mesenchymal cell proliferation | 0.00182495 | 2.88E-17 |
| GO:0030154 | cell differentiation | 0.00184529 | 1.77E-08 |
| GO:0007492 | endoderm development | 0.0021305 | 0.00497598 |
| GO:0030901 | midbrain development | 0.0021305 | 1.26E-06 |
| GO:0010595 | positive regulation of endothelial cell migration | 0.00223737 | 0.02729736 |
| GO:0043065 | positive regulation of apoptotic process | 0.00245285 | 4.03E-10 |
| GO:0045740 | positive regulation of DNA replication | 0.00245859 | 0.00679759 |
| GO:0048706 | embryonic skeletal system development | 0.00257293 | 0.00020375 |
| GO:0045597 | positive regulation of cell differentiation | 0.00280899 | 4.36E-06 |
| GO:0010862 | positive regulation of pathway-restricted SMAD protein phosphorylation | 0.00305488 | 6.90E-07 |
| GO:0045599 | negative regulation of fat cell differentiation | 0.00305488 | 6.90E-07 |
| GO:0007568 | aging | 0.00317746 | 8.86E-08 |
| GO:0048468 | cell development | 0.00318148 | 0.00040482 |
| GO:0009953 | dorsal/ventral pattern formation | 0.00318148 | 4.20E-16 |
| GO:0021915 | neural tube development | 0.00344197 | 1.25E-06 |
| GO:0006355 | regulation of transcription, DNA-templated | 0.0035698 | 3.72E-08 |
| GO:0001756 | somitogenesis | 0.00413529 | 3.60E-07 |
| GO:0001666 | response to hypoxia | 0.00425666 | 8.11E-06 |
| GO:0001541 | ovarian follicle development | 0.0042811 | 0.00102775 |
| GO:0043408 | regulation of MAPK cascade | 0.00457982 | 0.00022813 |
| GO:0071560 | cellular response to transforming growth factor beta stimulus | 0.00488793 | 4.73E-05 |
| GO:0048812 | neuron projection morphogenesis | 0.00520538 | 0.00186241 |
| GO:0003007 | heart morphogenesis | 0.00536758 | 1.28E-12 |
| GO:0071363 | cellular response to growth factor stimulus | 0.00536758 | 6.98E-05 |
| GO:0048839 | inner ear development | 0.0056989 | 8.92E-05 |
| GO:0090263 | positive regulation of canonical Wnt signaling pathway | 0.00586801 | 3.47E-11 |
| GO:0001938 | positive regulation of endothelial cell proliferation | 0.00586801 | 0.03967443 |
| GO:0050680 | negative regulation of epithelial cell proliferation | 0.00674768 | 1.02E-11 |
| GO:0000187 | activation of MAPK activity | 0.00749191 | 0.00128144 |
| GO:0060548 | negative regulation of cell death | 0.00827159 | 0.02477654 |
| GO:0048661 | positive regulation of smooth muscle cell proliferation | 0.00827159 | 1.38E-05 |
| GO:0007050 | cell cycle arrest | 0.008472 | 1.56E-05 |
| GO:0016485 | protein processing | 0.00950655 | 0.01045742 |
| GO:0007417 | central nervous system development | 0.01015302 | 2.40E-08 |
| GO:0061312 | BMP signaling pathway involved in heart development | 0.0102439 | 0.00041145 |
| GO:0051897 | positive regulation of protein kinase B signaling | 0.01196962 | 0.00160387 |
| GO:0050731 | positive regulation of peptidyl-tyrosine phosphorylation | 0.01292758 | 1.37E-07 |
| GO:1904948 | midbrain dopaminergic neuron differentiation | 0.0136359 | 0.01964074 |
| GO:0043410 | positive regulation of MAPK cascade | 0.01366739 | 7.62E-11 |
| GO:0051092 | positive regulation of NF-kappaB transcription factor activity | 0.0149405 | 0.01163548 |
| GO:0061384 | heart trabecula morphogenesis | 0.01532769 | 0.0247888 |
| GO:0008584 | male gonad development | 0.0157281 | 7.20E-05 |
| GO:0009952 | anterior/posterior pattern specification | 0.0157281 | 4.58E-17 |
| GO:0006468 | protein phosphorylation | 0.01621133 | 3.88E-05 |
| GO:0060039 | pericardium development | 0.01701666 | 0.03041854 |
| GO:0007165 | signal transduction | 0.01815958 | 0.01179635 |
| GO:0007435 | salivary gland morphogenesis | 0.01870283 | 0.00305644 |
| GO:0007219 | Notch signaling pathway | 0.01934164 | 0.00078917 |
| GO:0030308 | negative regulation of cell growth | 0.01934164 | 4.31E-07 |
| GO:0042493 | response to drug | 0.01991886 | 3.17E-11 |
| GO:0060272 | embryonic skeletal joint morphogenesis | 0.02038621 | 0.00025275 |
| GO:0060389 | pathway-restricted SMAD protein phosphorylation | 0.02206678 | 0.0050812 |
| GO:0006366 | transcription from RNA polymerase II promoter | 0.02326995 | 0.04072805 |
| GO:0007411 | axon guidance | 0.02682883 | 9.88E-07 |
| GO:0061036 | positive regulation of cartilage development | 0.02709179 | 0.00934741 |
| GO:0035567 | non-canonical Wnt signaling pathway | 0.02876123 | 0.01111749 |
| GO:0060038 | cardiac muscle cell proliferation | 0.02876123 | 0.01111749 |
| GO:0035137 | hindlimb morphogenesis | 0.02876123 | 1.99E-07 |
| GO:0060037 | pharyngeal system development | 0.0304279 | 0.01306762 |
| GO:0017015 | regulation of transforming growth factor beta receptor signaling pathway | 0.03209181 | 0.00169332 |
| GO:0046579 | positive regulation of Ras protein signal transduction | 0.03706695 | 0.02270901 |
| GO:0006351 | transcription, DNA-templated | 0.03711583 | 4.28E-09 |
| GO:0090190 | positive regulation of branching involved in ureteric bud morphogenesis | 0.03871983 | 3.17E-05 |
| GO:0045778 | positive regulation of ossification | 0.03871983 | 5.75E-09 |
| GO:0035050 | embryonic heart tube development | 0.03871983 | 0.02558431 |
| GO:0009880 | embryonic pattern specification | 0.04036995 | 4.13E-05 |
| GO:0048646 | anatomical structure formation involved in morphogenesis | 0.04036995 | 2.99E-06 |
| GO:0045216 | cell-cell junction organization | 0.04036995 | 0.02864555 |
| GO:0032967 | positive regulation of collagen biosynthetic process | 0.04366199 | 0.03532201 |
| GO:0035987 | endodermal cell differentiation | 0.04530391 | 8.45E-05 |
| GO:0042476 | odontogenesis | 0.04530391 | 5.02E-07 |
| GO:0042981 | regulation of apoptotic process | 0.04548052 | 8.66E-07 |
| GO:0035108 | limb morphogenesis | 0.0469431 | 1.11E-16 |
| GO:0055010 | ventricular cardiac muscle tissue morphogenesis | 0.0469431 | 0.00097892 |
| GO:0043392 | negative regulation of DNA binding | 0.04857957 | 0.00012938 |
| GO:0071542 | dopaminergic neuron differentiation | 0.04857957 | 6.06E-08 |

Supplementary Table S11. The enriched Uberon terms that are common to the predicted genes and genes with original annotations for the pectoral fin. The enriched terms are sorted based on the p-value of those terms for the predicted genes.

| Uberon term identifier | Uberon term name | P-value for the predicted genes | P-value for the original genes |
| --- | --- | --- | --- |
| uberon_0011610 | ceratohyal cartilage | 3.81E-13 | 1.90E-14 |
| uberon_0005886 | post-hyoid pharyngeal arch skeleton | 1.88E-12 | 2.56E-19 |
| uberon_2001516 | ceratobranchial cartilage | 9.63E-11 | 0.00102618 |
| uberon_0011242 | ethmoid cartilage | 9.63E-11 | 0.00029496 |
| uberon_0003107 | meckel's cartilage | 1.84E-10 | 3.21E-12 |
| uberon_0003079 | floor plate | 1.42E-09 | 0.03374939 |
| uberon_0002533 | post-anal tail bud | 9.84E-08 | 0.00338073 |
| uberon_0007215 | trabecula cranii | 1.50E-07 | 0.00035249 |
| uberon_0008896 | post-hyoid pharyngeal arch | 1.85E-07 | 2.03E-11 |
| uberon_0007812 | post-anal tail | 6.22E-07 | 0.00521186 |
| uberon_0002329 | somite | 7.57E-07 | 0.00590808 |
| uberon_0001016 | nervous system | 2.59E-06 | 0.03822373 |
| uberon_0002328 | notochord | 6.81E-06 | 0.00808243 |
| uberon_0011607 | hyomandibular cartilage | 1.03E-05 | 6.45E-06 |
| uberon_0001049 | neural tube | 2.10E-05 | 0.0227542 |
| uberon_0004752 | palatoquadrate cartilage | 2.14E-05 | 2.87E-09 |
| uberon_2000250 | opercle | 2.74E-05 | 1.51E-05 |
| uberon_0000165 | mouth | 3.80E-05 | 0.00014836 |
| uberon_0000468 | multicellular organism | 4.20E-05 | 1.09E-09 |
| uberon_0001894 | diencephalon | 5.15E-05 | 0.0201601 |
| uberon_0003901 | horizontal septum | 7.47E-05 | 0.00192572 |
| uberon_2001256 | lateral floor plate | 8.54E-05 | 0.03796748 |
| uberon_0003051 | ear vesicle | 0.00019633 | 2.14E-06 |
| uberon_0005945 | neurocranial trabecula | 0.0002281 | 0.00646992 |
| uberon_2001239 | ceratobranchial 5 bone | 0.00033971 | 0.00024377 |
| uberon_0003099 | cranial neural crest | 0.00040528 | 0.017947 |
| uberon_0002028 | hindbrain | 0.0005451 | 0.00387881 |
| uberon_0000044 | dorsal root ganglion | 0.00085431 | 0.03471745 |
| uberon_0000965 | lens of camera-type eye | 0.00096984 | 0.02462029 |
| uberon_0001890 | forebrain | 0.00100664 | 0.00629125 |
| uberon_0001708 | jaw skeleton | 0.00115871 | 7.84E-20 |
| uberon_2000694 | ceratobranchial 5 tooth | 0.00170211 | 7.74E-06 |
| uberon_2001089 | myoseptum | 0.00170211 | 0.01399613 |
| uberon_0002240 | spinal cord | 0.0017408 | 0.00254371 |
| uberon_0001264 | pancreas | 0.00181488 | 0.00023088 |
| uberon_0003936 | postoptic commissure | 0.00248307 | 0.00081772 |
| uberon_0007329 | pancreatic duct | 0.003566 | 0.00556203 |
| uberon_0011615 | basihyal cartilage | 0.00372936 | 0.0016559 |
| uberon_0003077 | paraxial mesoderm | 0.00425656 | 0.00706401 |
| uberon_2000040 | median fin fold | 0.0046312 | 1.80E-17 |
| uberon_2000558 | posterior macula | 0.00500389 | 0.0087837 |
| uberon_0000033 | head | 0.0061048 | 6.69E-08 |
| uberon_0001032 | sensory system | 0.0062966 | 4.43E-05 |
| uberon_0003072 | optic cup | 0.00666504 | 0.01290014 |
| uberon_0003098 | optic stalk | 0.00666504 | 0.00167394 |
| uberon_4000164 | caudal fin | 0.00707562 | 9.31E-11 |
| uberon_2001069 | ventral fin fold | 0.00853811 | 2.60E-07 |
| uberon_0009635 | parachordal cartilage | 0.00854196 | 3.23E-06 |
| uberon_0004741 | cleithrum | 0.01174598 | 8.41E-06 |
| uberon_0003278 | skeleton of lower jaw | 0.0124274 | 1.01E-06 |
| uberon_0001898 | hypothalamus | 0.01291407 | 0.0053699 |
| uberon_0003011 | facial motor nucleus | 0.01291407 | 0.0053699 |
| uberon_0000926 | mesoderm | 0.01413075 | 0.03471745 |
| uberon_0000948 | heart | 0.01508289 | 0.00333015 |
| uberon_0001891 | midbrain | 0.01623867 | 0.01553909 |
| uberon_0000019 | camera-type eye | 0.01756108 | 0.00298894 |
| uberon_0005884 | hyoid arch skeleton | 0.0224617 | 0.00079602 |
| uberon_0000935 | anterior commissure | 0.02393768 | 0.00050308 |
| uberon_0011085 | palatoquadrate arch | 0.02551321 | 0.01749 |
| uberon_0004375 | bone of free limb or fin | 0.03295212 | 0.0093632 |
| uberon_0001703 | neurocranium | 0.03765523 | 1.19E-07 |
| uberon_0001976 | epithelium of esophagus | 0.04102281 | 0.0137538 |
| uberon_0002348 | epicardium | 0.04102281 | 0.00099481 |
| uberon_2002193 | dorsolateral septum | 0.04102281 | 0.00099481 |
| uberon_0011004 | pharyngeal arch cartilage | 0.04146017 | 2.83E-10 |
| uberon_2001257 | medial floor plate | 0.04902763 | 0.0188574 |
| uberon_0005598 | trunk somite | 0.04902763 | 0.0188574 |
| uberon_0003114 | pharyngeal arch 3 | 0.04902763 | 0.0188574 |

Supplementary Table S12. The enriched Uberon terms that are common to the predicted genes and genes with original annotations for the pelvic fin. The enriched terms are sorted based on the p-value of those terms for the predicted genes.

| Uberon term identifier | Uberon term name | P-value for the predicted genes | P-value for the original genes |
| --- | --- | --- | --- |
| uberon_0000151 | pectoral fin | 3.26E-07 | 1.42E-09 |
| uberon_2000040 | median fin fold | 5.40E-06 | 8.70E-05 |
| uberon_4000163 | anal fin | 0.00016979 | 4.43E-11 |
| uberon_4000172 | lepidotrichium | 0.00351758 | 2.42E-06 |
| uberon_0003097 | dorsal fin | 0.02306165 | 1.03E-11 |

Supplementary Table S13. The top 100 enriched Uberon terms that are common to the predicted genes and genes with original annotations for the forelimb. The enriched terms are sorted based on the p-value of those terms for the predicted genes.

| Uberon term identifier | Uberon term name | P-value for the predicted genes | P-value for the original genes |
| --- | --- | --- | --- |
| uberon_0001703 | neurocranium | 9.49E-19 | 3.67E-52 |
| uberon_0011156 | facial skeleton | 2.21E-18 | 1.36E-65 |
| uberon_0003128 | cranium | 5.88E-16 | 9.57E-79 |
| uberon_0001676 | occipital bone | 7.46E-16 | 4.99E-20 |
| uberon_0001708 | jaw skeleton | 2.83E-15 | 2.61E-61 |
| uberon_0004716 | conceptus | 7.34E-14 | 2.32E-48 |
| uberon_0007811 | craniocervical region | 7.49E-14 | 4.41E-77 |
| uberon_0000209 | tetrapod frontal bone | 1.20E-13 | 1.61E-17 |
| uberon_0002091 | appendicular skeleton | 1.27E-13 | 1.39E-187 |
| uberon_0001434 | skeletal system | 2.87E-13 | 2.37E-122 |
| uberon_0005944 | axial skeleton plus cranial skeleton | 3.25E-13 | 1.13E-87 |
| uberon_0000165 | mouth | 1.11E-12 | 3.64E-61 |
| uberon_0002517 | basicranium | 1.36E-12 | 2.14E-27 |
| uberon_0001692 | basioccipital bone | 2.48E-12 | 0.00016976 |
| uberon_0003252 | thoracic rib cage | 2.49E-12 | 4.72E-74 |
| uberon_0001756 | middle ear | 3.11E-11 | 1.26E-24 |
| uberon_0001456 | face | 5.09E-11 | 6.78E-61 |
| uberon_0006428 | basisphenoid bone | 7.87E-11 | 5.22E-16 |
| uberon_0001049 | neural tube | 1.06E-10 | 3.88E-29 |
| uberon_0001684 | mandible | 1.12E-10 | 1.56E-51 |
| uberon_0000210 | tetrapod parietal bone | 1.25E-10 | 1.03E-16 |
| uberon_0003216 | hard palate | 1.35E-10 | 4.89E-27 |
| uberon_0004747 | supraoccipital bone | 4.05E-10 | 6.38E-08 |
| uberon_0000033 | head | 6.18E-10 | 2.24E-58 |
| uberon_0001690 | ear | 6.79E-10 | 9.92E-30 |
| uberon_0001716 | secondary palate | 1.22E-09 | 1.29E-40 |
| uberon_0001685 | hyoid bone | 1.35E-09 | 1.21E-10 |
| uberon_0001677 | sphenoid bone | 1.46E-09 | 8.95E-27 |
| uberon_0001689 | malleus bone | 1.90E-09 | 8.78E-13 |
| uberon_0002105 | vestibulo-auditory system | 2.07E-09 | 1.06E-28 |
| uberon_0001007 | digestive system | 3.28E-09 | 1.27E-40 |
| uberon_0002229 | interparietal bone | 4.28E-09 | 3.38E-18 |
| uberon_0002228 | rib | 4.82E-09 | 2.08E-59 |
| uberon_0001004 | respiratory system | 1.82E-08 | 9.08E-34 |
| uberon_0000955 | brain | 2.00E-08 | 1.48E-14 |
| uberon_0001678 | temporal bone | 2.39E-08 | 1.11E-14 |
| uberon_0006721 | alisphenoid bone | 3.22E-08 | 5.08E-06 |
| uberon_0003450 | upper jaw incisor | 6.27E-08 | 9.49E-07 |
| uberon_0003051 | ear vesicle | 9.95E-08 | 1.97E-06 |
| uberon_0008828 | presphenoid bone | 9.95E-08 | 1.88E-12 |
| uberon_0000922 | embryo | 1.55E-07 | 1.90E-25 |
| uberon_0003451 | lower jaw incisor | 1.66E-07 | 2.76E-08 |
| uberon_0002218 | tympanic ring | 1.66E-07 | 1.04E-15 |
| uberon_0002418 | cartilage tissue | 5.94E-07 | 8.02E-30 |
| uberon_0003655 | molar tooth | 6.91E-07 | 1.66E-12 |
| uberon_0003966 | gonial bone | 9.41E-07 | 0.00011778 |
| uberon_0010389 | pterygoid bone | 9.41E-07 | 6.42E-06 |
| uberon_0002510 | anterior fontanel | 1.34E-06 | 0.00254357 |
| uberon_0002328 | notochord | 1.35E-06 | 2.01E-07 |
| uberon_0001066 | intervertebral disk | 1.77E-06 | 2.31E-19 |
| uberon_0011933 | vibrissa unit | 2.02E-06 | 3.00E-05 |
| uberon_0001738 | thyroid cartilage | 2.79E-06 | 1.22E-07 |
| uberon_0001681 | nasal bone | 3.03E-06 | 6.39E-22 |
| uberon_0001890 | forebrain | 3.25E-06 | 3.76E-13 |
| uberon_0004649 | sphenoid bone pterygoid process | 4.50E-06 | 1.93E-08 |
| uberon_0002329 | somite | 4.85E-06 | 6.28E-09 |
| uberon_0004660 | mandible coronoid process | 5.01E-06 | 4.14E-07 |
| uberon_0005871 | palatine process of maxilla | 5.01E-06 | 1.37E-09 |
| uberon_0001695 | squamous part of temporal bone | 6.79E-06 | 7.79E-07 |
| uberon_0005942 | hair outer root sheath | 6.79E-06 | 0.01132058 |
| uberon_0003075 | neural plate | 8.18E-06 | 0.00165765 |
| uberon_0000924 | ectoderm | 8.52E-06 | 6.64E-17 |
| uberon_0018242 | palatine bone horizontal plate | 9.74E-06 | 1.20E-07 |
| uberon_0001694 | petrous part of temporal bone | 9.74E-06 | 2.04E-05 |
| uberon_0002073 | hair follicle | 1.01E-05 | 7.57E-10 |
| uberon_0002224 | thoracic cavity | 1.06E-05 | 9.88E-09 |
| uberon_0000401 | mandibular ramus | 1.15E-05 | 4.55E-11 |
| uberon_0001737 | larynx | 1.80E-05 | 1.91E-10 |
| uberon_0006772 | long bone epiphyseal plate hypertrophic zone | 1.97E-05 | 2.91E-40 |
| uberon_0001687 | stapes bone | 2.19E-05 | 6.84E-08 |
| uberon_0003861 | neural arch | 2.26E-05 | 2.25E-18 |
| uberon_0002470 | autopod region | 2.28E-05 | 5.54E-91 |
| uberon_0002103 | hindlimb | 2.43E-05 | 2.57E-138 |
| uberon_0001752 | enamel | 2.98E-05 | 0.00014902 |
| uberon_0003107 | meckel's cartilage | 2.98E-05 | 4.98E-17 |
| uberon_0002413 | cervical vertebra | 3.01E-05 | 1.16E-14 |
| uberon_0001894 | diencephalon | 3.06E-05 | 5.38E-11 |
| uberon_0001706 | nasal septum | 3.53E-05 | 1.69E-09 |
| uberon_0001092 | vertebral bone 1 | 3.73E-05 | 1.52E-10 |
| uberon_0001682 | palatine bone | 3.93E-05 | 1.83E-10 |
| uberon_0002416 | integumental system | 3.96E-05 | 2.80E-16 |
| uberon_0000004 | nose | 4.01E-05 | 2.04E-24 |
| uberon_0002028 | hindbrain | 4.59E-05 | 3.77E-08 |
| uberon_0005354 | malleus processus brevis | 4.75E-05 | 0.00721838 |
| uberon_0005619 | secondary palatal shelf | 4.82E-05 | 3.00E-17 |
| uberon_0001130 | vertebral column | 5.03E-05 | 5.26E-27 |
| uberon_0001691 | external ear | 5.83E-05 | 3.87E-15 |
| uberon_0001232 | collecting duct of renal tubule | 6.68E-05 | 8.78E-05 |
| uberon_0003982 | mature ovarian follicle | 6.68E-05 | 0.00364138 |
| uberon_0001075 | bony vertebral centrum | 6.98E-05 | 6.89E-13 |
| uberon_0001675 | trigeminal ganglion | 6.98E-05 | 0.000653 |
| uberon_0001998 | sternocostal joint | 7.60E-05 | 1.87E-09 |
| uberon_0003461 | shoulder bone | 8.04E-05 | 3.33E-19 |
| uberon_0002129 | cerebellar cortex | 8.27E-05 | 0.00314678 |
| uberon_0002414 | lumbar vertebra | 8.67E-05 | 6.55E-10 |
| uberon_0005867 | mandibular prominence | 8.70E-05 | 0.00080115 |
| uberon_0001037 | strand of hair | 0.00010094 | 3.10E-07 |
| uberon_0003941 | cerebellum anterior vermis | 0.00010272 | 0.01504756 |
| uberon_0002217 | synovial joint | 0.00010272 | 0.00100864 |
| uberon_0001043 | esophagus | 0.00011262 | 6.33E-07 |

Supplementary Table S14. The top 100 enriched Uberon terms that are common to the predicted genes and genes with original annotations for the hindlimb. The enriched terms are sorted based on the p-value of those terms for the predicted genes.

| Uberon term identifier | Uberon term name | P-value for the predicted genes | P-value for the original genes |
| --- | --- | --- | --- |
| uberon_0003128 | cranium | 4.01E-24 | 5.82E-57 |
| uberon_0011156 | facial skeleton | 1.11E-22 | 4.74E-53 |
| uberon_0001434 | skeletal system | 1.42E-21 | 1.19E-223 |
| uberon_0005944 | axial skeleton plus cranial skeleton | 4.39E-21 | 3.58E-88 |
| uberon_0007811 | craniocervical region | 3.88E-19 | 2.99E-57 |
| uberon_0001456 | face | 4.19E-19 | 2.62E-45 |
| uberon_0000033 | head | 1.62E-18 | 5.98E-44 |
| uberon_0001708 | jaw skeleton | 2.11E-17 | 1.39E-53 |
| uberon_0001703 | neurocranium | 3.94E-17 | 2.49E-44 |
| uberon_0000165 | mouth | 3.90E-16 | 1.59E-47 |
| uberon_0002091 | appendicular skeleton | 1.44E-15 | 0 |
| uberon_0002105 | vestibulo-auditory system | 1.25E-14 | 1.85E-25 |
| uberon_0001690 | ear | 3.73E-14 | 1.27E-25 |
| uberon_0001684 | mandible | 4.59E-14 | 8.20E-43 |
| uberon_0001007 | digestive system | 1.41E-13 | 1.04E-28 |
| uberon_0006333 | snout | 2.45E-13 | 8.50E-22 |
| uberon_0002483 | trabecular bone tissue | 1.84E-12 | 4.59E-72 |
| uberon_0000210 | tetrapod parietal bone | 1.91E-12 | 6.03E-14 |
| uberon_0001756 | middle ear | 1.96E-12 | 9.35E-20 |
| uberon_0001004 | respiratory system | 4.07E-12 | 8.92E-32 |
| uberon_0000209 | tetrapod frontal bone | 3.47E-11 | 3.65E-13 |
| uberon_0000383 | musculature of body | 4.96E-11 | 1.02E-56 |
| uberon_0001630 | muscle organ | 1.56E-10 | 0.00011844 |
| uberon_0002517 | basicranium | 3.84E-10 | 1.66E-17 |
| uberon_0001049 | neural tube | 7.14E-10 | 2.88E-19 |
| uberon_0001676 | occipital bone | 1.39E-09 | 1.36E-12 |
| uberon_0002229 | interparietal bone | 1.95E-09 | 1.81E-10 |
| uberon_0001677 | sphenoid bone | 2.11E-09 | 1.50E-13 |
| uberon_0000955 | brain | 3.27E-09 | 2.55E-14 |
| uberon_0006428 | basisphenoid bone | 4.44E-09 | 2.35E-07 |
| uberon_0002228 | rib | 4.70E-09 | 4.14E-37 |
| uberon_0003252 | thoracic rib cage | 9.23E-09 | 5.47E-50 |
| uberon_0004716 | conceptus | 1.03E-08 | 3.81E-21 |
| uberon_0003450 | upper jaw incisor | 1.09E-08 | 2.32E-05 |
| uberon_0006772 | long bone epiphyseal plate hypertrophic zone | 2.06E-08 | 6.26E-55 |
| uberon_0002416 | integumental system | 3.26E-08 | 9.53E-16 |
| uberon_0003451 | lower jaw incisor | 3.59E-08 | 2.90E-09 |
| uberon_0000014 | zone of skin | 4.54E-08 | 9.99E-19 |
| uberon_0001689 | malleus bone | 5.05E-08 | 2.31E-12 |
| uberon_0001130 | vertebral column | 5.84E-08 | 2.46E-37 |
| uberon_0003107 | meckel's cartilage | 1.03E-07 | 2.75E-12 |
| uberon_0004535 | cardiovascular system | 2.17E-07 | 5.30E-28 |
| uberon_0002113 | kidney | 2.72E-07 | 3.61E-13 |
| uberon_0006721 | alisphenoid bone | 4.21E-07 | 6.91E-06 |
| uberon_0000474 | female reproductive system | 4.37E-07 | 7.29E-19 |
| uberon_0003216 | hard palate | 4.55E-07 | 3.86E-17 |
| uberon_0000019 | camera-type eye | 4.68E-07 | 1.67E-23 |
| uberon_0002516 | epiphyseal plate | 4.92E-07 | 4.29E-72 |
| uberon_0001678 | temporal bone | 6.21E-07 | 4.05E-12 |
| uberon_0002418 | cartilage tissue | 6.60E-07 | 3.22E-14 |
| uberon_0001890 | forebrain | 6.60E-07 | 2.57E-12 |
| uberon_0005070 | anterior neuropore | 6.63E-07 | 0.04722767 |
| uberon_0001692 | basioccipital bone | 9.19E-07 | 0.00016117 |
| uberon_0002101 | limb | 9.91E-07 | 4.55E-08 |
| uberon_0001008 | renal system | 1.01E-06 | 5.55E-21 |
| uberon_0002104 | visual system | 1.18E-06 | 1.15E-25 |
| uberon_0002397 | maxilla | 1.25E-06 | 8.65E-29 |
| uberon_0001681 | nasal bone | 1.28E-06 | 2.59E-17 |
| uberon_0002370 | thymus | 1.86E-06 | 5.49E-13 |
| uberon_0002405 | immune system | 1.88E-06 | 1.05E-06 |
| uberon_0003697 | abdominal wall | 2.07E-06 | 1.51E-07 |
| uberon_0002218 | tympanic ring | 2.14E-06 | 5.31E-11 |
| uberon_0001716 | secondary palate | 2.86E-06 | 2.15E-25 |
| uberon_0000948 | heart | 2.92E-06 | 1.31E-18 |
| uberon_0002390 | hematopoietic system | 3.26E-06 | 3.56E-07 |
| uberon_0002544 | digit | 3.57E-06 | 2.75E-48 |
| uberon_0003461 | shoulder bone | 4.45E-06 | 2.81E-14 |
| uberon_0001752 | enamel | 5.00E-06 | 1.17E-06 |
| uberon_0006849 | scapula | 5.00E-06 | 1.09E-18 |
| uberon_0002491 | lambdoid suture | 5.12E-06 | 0.00065007 |
| uberon_0003975 | internal female genitalia | 5.36E-06 | 5.15E-14 |
| uberon_0000323 | late embryo | 8.00E-06 | 4.11E-10 |
| uberon_0002428 | limb bone | 8.82E-06 | 0.00139297 |
| uberon_0000309 | body wall | 9.97E-06 | 7.79E-08 |
| uberon_0002048 | lung | 1.23E-05 | 3.26E-20 |
| uberon_0002470 | autopod region | 1.37E-05 | 1.95E-55 |
| uberon_0001229 | renal corpuscle | 1.41E-05 | 4.22E-05 |
| uberon_0001075 | bony vertebral centrum | 1.53E-05 | 1.30E-09 |
| uberon_0002328 | notochord | 1.71E-05 | 2.82E-07 |
| uberon_0001285 | nephron | 1.79E-05 | 1.62E-06 |
| uberon_0002073 | hair follicle | 1.82E-05 | 3.40E-07 |
| uberon_0001225 | cortex of kidney | 2.68E-05 | 1.80E-05 |
| uberon_0001987 | placenta | 2.68E-05 | 1.80E-05 |
| uberon_0006861 | diaphysis proper | 3.28E-05 | 3.71E-12 |
| uberon_0006771 | long bone epiphyseal plate proliferative zone | 3.45E-05 | 2.52E-29 |
| uberon_0001016 | nervous system | 3.73E-05 | 5.57E-07 |
| uberon_0005956 | outflow part of left ventricle | 3.78E-05 | 0.00017418 |
| uberon_0001037 | strand of hair | 4.12E-05 | 3.95E-06 |
| uberon_0002137 | aortic valve | 4.12E-05 | 0.00021436 |
| uberon_0001695 | squamous part of temporal bone | 4.43E-05 | 0.00058499 |
| uberon_0010166 | coat of hair | 5.19E-05 | 1.65E-06 |
| uberon_0002329 | somite | 5.99E-05 | 0.00015746 |
| uberon_0001758 | periodontium | 6.34E-05 | 3.57E-09 |
| uberon_0003051 | ear vesicle | 6.89E-05 | 0.00027719 |
| uberon_0008828 | presphenoid bone | 6.89E-05 | 3.51E-08 |
| uberon_0001683 | jugal bone | 6.89E-05 | 8.05E-11 |
| uberon_0002224 | thoracic cavity | 6.89E-05 | 5.32E-05 |
| uberon_0002412 | vertebra | 7.08E-05 | 2.93E-18 |
| uberon_0000945 | stomach | 7.42E-05 | 0.00021632 |
| uberon_0000012 | somatic nervous system | 7.46E-05 | 1.69E-27 |

Supplementary Table S15. Comparison of the 37 genes that are common to the pectoral fin and forelimb modules (conserved genes). The genes are ordered according to the rank in the pectoral fin module.

| Zebrafish gene name | Annotation type | Weighted degree | Rank in the module | Mouse gene name | Annotation type | Weighted degree | Rank in the module |
| --- | --- | --- | --- | --- | --- | --- | --- |
| *shha* | direct annotation to the pectoral fin | 36.6983 | 1 | *shh* | direct annotation to the forelimb | 37.6249 | 4 |
| *bmp4* | predicted | 35.5652 | 2 | *bmp4* | direct annotation to the forelimb | 47.2641 | 1 |
| *bmp2b* | predicted | 34.3109 | 3 | *bmp2* | direct annotation to the forelimb | 31.3004 | 13 |
| *wnt3a* | predicted | 31.4927 | 4 | *wnt3a* | predicted | 30.9124 | 14 |
| *fgf8a* | predicted | 31.0094 | 5 | *fgf8* | direct annotation to the forelimb | 34.0672 | 8 |
| *gli2a* | predicted | 27.2862 | 8 | *gli2* | direct annotation to the forelimb | 33.0563 | 11 |
| *bmp7a* | direct annotation to the pectoral fin | 25.8141 | 11 | *bmp7* | predicted | 34.2132 | 7 |
| *fgfr1a* | predicted | 25.6442 | 12 | *fgfr1* | direct annotation to the forelimb | 33.4360 | 10 |
| *fgf10a* | direct annotation to the pectoral fin | 24.2622 | 17 | *fgf10* | direct annotation to the forelimb | 22.7474 | 33 |
| *smo* | direct annotation to the pectoral fin | 23.6222 | 19 | *smo* | direct annotation to the forelimb | 28.7826 | 20 |
| *wnt4a* | predicted | 20.4183 | 29 | *wnt4* | predicted | 27.9668 | 23 |
| *ptch1* | direct annotation to the pectoral fin | 18.8304 | 30 | *ptch1* | predicted | 27.3374 | 24 |
| *ihha* | predicted | 18.1064 | 32 | *ihh* | direct annotation to the forelimb | 29.6653 | 17 |
| *tbx5a* | direct annotation to the pectoral fin | 16.7968 | 35 | *tbx5* | direct annotation to the forelimb | 8.6229 | 86 |
| *aldh1a2* | direct annotation to the pectoral fin | 16.6865 | 36 | *aldh1a2* | annotated to a part or bud | 7.1983 | 100 |
| *hand2* | direct annotation to the pectoral fin | 16.4984 | 38 | *hand2* | direct annotation to the forelimb | 13.8461 | 62 |
| *zic2a* | predicted | 15.2976 | 40 | *zic2* | annotated to a part or bud | 7.3701 | 99 |
| *tcf7* | direct annotation to the pectoral fin | 15.0392 | 42 | *tcf7* | direct annotation to the forelimb | 7.7495 | 93 |
| *msx1a* | predicted | 14.6308 | 45 | *msx1* | predicted | 23.4172 | 31 |
| *met* | direct annotation to the pectoral fin | 14.5234 | 46 | *met* | direct annotation to the forelimb | 13.9973 | 60 |
| *esr2a* | predicted | 13.2000 | 51 | *esr2* | direct annotation to the forelimb | 7.6748 | 95 |
| *apc* | direct annotation to the pectoral fin | 9.3808 | 61 | *apc* | direct annotation to the forelimb | 12.6243 | 67 |
| *tfap2a* | predicted | 9.2007 | 62 | *tfap2a* | direct annotation to the forelimb | 9.3891 | 78 |
| *dlx5a* | annotated to a part or bud | 8.9299 | 81 | *dlx5* | predicted | 16.1115 | 51 |
| *sox9a* | direct annotation to the pectoral fin | 8.8290 | 83 | *sox9* | direct annotation to the forelimb | 29.7370 | 15 |
| *disp1* | direct annotation to the pectoral fin | 8.6768 | 85 | *disp1* | direct annotation to the forelimb | 1.8115 | 183 |
| *mecom* | predicted | 8.4715 | 89 | *mecom* | annotated to a part or bud | 3.3452 | 155 |
| *sox9b* | predicted | 7.2558 | 104 | *sox9* | direct annotation to the forelimb | 29.7370 | 15 |
| *thraa* | direct annotation to the pectoral fin | 7.0911 | 105 | *thra* | direct annotation to the forelimb | 3.2473 | 157 |
| *cyp26b1* | direct annotation to the pectoral fin | 6.5056 | 111 | *cyp26b1* | direct annotation to the forelimb | 3.5223 | 151 |
| *ctgfa* | direct annotation to the pectoral fin | 5.3072 | 124 | *ctgf* | direct annotation to the forelimb | 20.3372 | 41 |
| *wls* | direct annotation to the pectoral fin | 4.8573 | 133 | *wls* | direct annotation to the forelimb | 6.6655 | 107 |
| *osr1* | direct annotation to the pectoral fin | 4.6429 | 136 | *osr1* | annotated to a part or bud | 0.3046 | 241 |
| *sparc* | direct annotation to the pectoral fin | 3.7395 | 148 | *sparc* | direct annotation to the forelimb | 12.4871 | 69 |
| *chsy1* | direct annotation to the pectoral fin | 1.9657 | 176 | *chsy1* | annotated to a part or bud | 1.0702 | 210 |
| *rspo2* | direct annotation to the pectoral fin | 0.7583 | 200 | *rspo2* | direct annotation to the forelimb | 3.6481 | 149 |
| *pax1b* | direct annotation to the pectoral fin | 0.3795 | 209 | *pax1* | direct annotation to the forelimb | 6.2887 | 111 |

Supplementary Table S16. Comparison of the 81 genes that are common to the pelvic fin and hindlimb modules (conserved genes). The genes are ordered according to the rank in the pelvic fin module.

| Zebrafish gene name | Annotation type | Weighted degree | Rank in the module | Mouse gene name | Annotation type | Weighted degree | Rank in the module |
| --- | --- | --- | --- | --- | --- | --- | --- |
| *ctnnb1* | predicted | 106.2749 | 4 | *ctnnb1* | direct annotation to the hindlimb | 72.1808 | 3 |
| *smarca4a* | predicted | 97.4336 | 11 | *smarca4* | direct annotation to the hindlimb | 36.4034 | 56 |
| *kras* | predicted | 93.403 | 15 | *kras* | predicted | 70.7748 | 5 |
| *rac1a* | predicted | 92.9107 | 16 | *rac1* | direct annotation to the hindlimb | 24.2559 | 103 |
| *mapk14a* | predicted | 92.1789 | 17 | *mapk14* | direct annotation to the hindlimb | 71.7856 | 4 |
| *src* | predicted | 91.7256 | 18 | *src* | direct annotation to the hindlimb | 70.3811 | 6 |
| *fgfr1a* | predicted | 91.2553 | 19 | *fgfr1* | direct annotation to the hindlimb | 53.223 | 16 |
| *met* | predicted | 90.8735 | 20 | *met* | direct annotation to the hindlimb | 34.8637 | 64 |
| *tp53* | predicted | 83.8035 | 24 | *trp53* | direct annotation to the hindlimb | 82.9324 | 1 |
| *mtor* | predicted | 80.8121 | 28 | *mtor* | annotated to a part or bud | 38.6216 | 49 |
| *ptenb* | predicted | 78.531 | 32 | *pten* | annotated to a part or bud | 43.6208 | 35 |
| *bmp4* | predicted | 76.215 | 36 | *bmp4* | direct annotation to the hindlimb | 63.9989 | 10 |
| *bmp2b* | predicted | 68.2429 | 45 | *bmp2* | predicted | 45.0361 | 32 |
| *rac1b* | predicted | 67.0359 | 50 | *rac1* | direct annotation to the hindlimb | 24.2559 | 103 |
| *notch2* | predicted | 66.3735 | 54 | *notch2* | direct annotation to the hindlimb | 39.0854 | 47 |
| *fgfr2* | predicted | 63.0559 | 67 | *fgfr2* | direct annotation to the hindlimb | 48.7644 | 24 |
| *ptena* | predicted | 60.7765 | 80 | *pten* | annotated to a part or bud | 43.6208 | 35 |
| *mapk14b* | predicted | 59.24 | 89 | *mapk14* | direct annotation to the hindlimb | 71.7856 | 4 |
| *smad2* | predicted | 58.5213 | 92 | *smad2* | predicted | 45.6514 | 30 |
| *wnt3a* | predicted | 57.9542 | 95 | *wnt3a* | direct annotation to the hindlimb | 42.6564 | 39 |
| *ret* | predicted | 57.2523 | 98 | *ret* | annotated to a part or bud | 23.5326 | 110 |
| *vegfaa* | predicted | 57.0152 | 100 | *vegfa* | annotated to a part or bud | 50.7476 | 20 |
| *junba* | predicted | 54.8789 | 109 | *junb* | direct annotation to the hindlimb | 15.5932 | 205 |
| *fgf8a* | predicted | 52.5593 | 122 | *fgf8* | direct annotation to the hindlimb | 46.6654 | 29 |
| *gli2a* | predicted | 52.1448 | 130 | *gli2* | direct annotation to the hindlimb | 42.8396 | 38 |
| *dicer1* | predicted | 50.473 | 140 | *dicer1* | direct annotation to the hindlimb | 18.3287 | 187 |
| *smad3a* | predicted | 49.5403 | 150 | *smad3* | annotated to a part or bud | 49.4434 | 22 |
| *hrasb* | predicted | 48.0853 | 162 | *hras* | predicted | 72.8342 | 2 |
| *shha* | predicted | 48.0501 | 163 | *shh* | direct annotation to the hindlimb | 44.4006 | 33 |
| *bmp2a* | predicted | 47.8327 | 165 | *bmp2* | predicted | 45.0361 | 32 |
| *fgf4* | predicted | 47.5358 | 167 | *fgf4* | annotated to a part or bud | 24.3609 | 102 |
| *bmp7a* | predicted | 46.1288 | 178 | *bmp7* | predicted | 48.2316 | 26 |
| *vegfab* | predicted | 45.8605 | 183 | *vegfa* | annotated to a part or bud | 50.7476 | 20 |
| *mysm1* | predicted | 43.0713 | 209 | *mysm1* | direct annotation to the hindlimb | 4.6408 | 394 |
| *fgf10a* | predicted | 42.2153 | 212 | *fgf10* | direct annotation to the hindlimb | 33.825 | 67 |
| *fgf10b* | predicted | 40.7886 | 219 | *fgf10* | direct annotation to the hindlimb | 33.825 | 67 |
| *sirt1* | predicted | 40.1693 | 232 | *sirt1* | annotated to a part or bud | 25.4897 | 94 |
| *bmpr1aa* | predicted | 36.7892 | 251 | *bmpr1a* | direct annotation to the hindlimb | 35.5242 | 61 |
| *lef1* | predicted | 35.665 | 259 | *lef1* | predicted | 36.7074 | 53 |
| *gli2b* | predicted | 33.6578 | 277 | *gli2* | direct annotation to the hindlimb | 42.8396 | 38 |
| *wnt4a* | predicted | 33.0982 | 286 | *wnt4* | predicted | 35.4014 | 62 |
| *smo* | predicted | 32.2549 | 293 | *smo* | direct annotation to the hindlimb | 34.086 | 66 |
| *sod2* | predicted | 30.6061 | 308 | *sod2* | annotated to a part or bud | 18.8196 | 178 |
| *gdf11* | direct annotation to the pelvic fin | 29.3173 | 321 | *gdf11* | direct annotation to the hindlimb | 8.6569 | 283 |
| *stat1b* | predicted | 28.9557 | 323 | *stat1* | direct annotation to the hindlimb | 33.3038 | 72 |
| *smad3b* | predicted | 27.6166 | 333 | *smad3* | annotated to a part or bud | 49.4434 | 22 |
| *casp3a* | predicted | 27.2208 | 340 | *casp3* | predicted | 49.0774 | 23 |
| *map3k14b* | predicted | 26.1706 | 350 | *map3k14* | annotated to a part or bud | 7.1186 | 312 |
| *aldh1a2* | predicted | 25.4504 | 354 | *aldh1a2* | annotated to a part or bud | 9.8933 | 263 |
| *gli3* | predicted | 25.0939 | 359 | *gli3* | direct annotation to the hindlimb | 38.3084 | 52 |
| *bmpr1ba* | predicted | 24.8781 | 361 | *bmpr1b* | annotated to a part or bud | 20.0326 | 131 |
| *ihha* | predicted | 24.0223 | 366 | *ihh* | direct annotation to the hindlimb | 33.7444 | 68 |
| *ptch1* | predicted | 23.8507 | 369 | *ptch1* | predicted | 38.8191 | 48 |
| *bmpr1ab* | predicted | 22.5899 | 386 | *bmpr1a* | direct annotation to the hindlimb | 35.5242 | 61 |
| *vdrb* | predicted | 22.4408 | 388 | *vdr* | direct annotation to the hindlimb | 15.8149 | 202 |
| *shhb* | predicted | 20.6403 | 404 | *shh* | direct annotation to the hindlimb | 44.4006 | 33 |
| *bmpr1bb* | predicted | 20.1116 | 408 | *bmpr1b* | annotated to a part or bud | 20.0326 | 131 |
| *fsta* | predicted | 17.8232 | 422 | *fst* | annotated to a part or bud | 27.0851 | 86 |
| *hand2* | predicted | 15.7932 | 437 | *hand2* | direct annotation to the hindlimb | 24.7719 | 101 |
| *msx1a* | predicted | 15.3589 | 442 | *msx1* | predicted | 26.6468 | 88 |
| *cx43* | predicted | 15.0787 | 444 | *gja1* | predicted | 28.3709 | 84 |
| *vdra* | predicted | 13.7382 | 454 | *vdr* | direct annotation to the hindlimb | 15.8149 | 202 |
| *pax3a* | predicted | 13.4122 | 458 | *pax3* | direct annotation to the hindlimb | 31.1909 | 76 |
| *ihhb* | predicted | 13.389 | 460 | *ihh* | direct annotation to the hindlimb | 33.7444 | 68 |
| *lamc1* | predicted | 13.3297 | 461 | *lamc1* | direct annotation to the hindlimb | 7.8573 | 299 |
| *desma* | predicted | 13.1643 | 462 | *des* | annotated to a part or bud | 14.3164 | 217 |
| *col1a1a* | predicted | 12.9905 | 465 | *col1a1* | direct annotation to the hindlimb | 43.3734 | 36 |
| *lmx1bb* | predicted | 11.9354 | 481 | *lmx1b* | annotated to a part or bud | 9.1733 | 271 |
| *col2a1a* | predicted | 11.8949 | 483 | *col2a1* | direct annotation to the hindlimb | 43.3479 | 37 |
| *tbx4* | direct annotation to the pelvic fin | 10.3408 | 496 | *tbx4* | direct annotation to the hindlimb | 10.3871 | 259 |
| *hspg2* | predicted | 9.6841 | 500 | *hspg2* | direct annotation to the hindlimb | 14.1006 | 220 |
| *fras1* | predicted | 9.0362 | 507 | *fras1* | annotated to a part or bud | 14.535 | 212 |
| *col1a2* | predicted | 6.6976 | 539 | *col1a2* | direct annotation to the hindlimb | 28.4794 | 83 |
| *scube2* | predicted | 6.4974 | 543 | *scube2* | direct annotation to the hindlimb | 3.4888 | 434 |
| *col11a1b* | predicted | 6.2965 | 545 | *col11a1* | direct annotation to the hindlimb | 14.3918 | 215 |
| *col11a1a* | predicted | 5.9988 | 550 | *col11a1* | direct annotation to the hindlimb | 14.3918 | 215 |
| *col1a1b* | predicted | 5.9171 | 554 | *col1a1* | direct annotation to the hindlimb | 43.3734 | 36 |
| *rspo2* | annotated to a part or bud | 4.0927 | 570 | *rspo2* | direct annotation to the hindlimb | 3.6481 | 429 |
| *frem2a* | predicted | 3.5366 | 577 | *frem2* | annotated to a part or bud | 1.5047 | 537 |
| *pitx1* | predicted | 2.8035 | 592 | *pitx1* | direct annotation to the hindlimb | 9.5149 | 267 |
| *pax3b* | predicted | 2.4374 | 595 | *pax3* | direct annotation to the hindlimb | 31.1909 | 76 |

Supplementary Table S17. The enriched Biological Process terms from the Gene Ontology for the mouse orthologs of the pectoral fin module-specific genes.

| Term identifier | Term name | P-value |
| --- | --- | --- |
| GO:0007275 | multicellular organism development | 5.68E-14 |
| GO:0045944 | positive regulation of transcription from RNA polymerase II promoter | 3.32E-10 |
| GO:0009887 | organ morphogenesis | 6.91E-09 |
| GO:0006355 | regulation of transcription, DNA-templated | 4.00E-08 |
| GO:0006351 | transcription, DNA-templated | 3.72E-07 |
| GO:0060070 | canonical Wnt signaling pathway | 4.94E-07 |
| GO:0043588 | skin development | 6.59E-07 |
| GO:0042475 | odontogenesis of dentin-containing tooth | 1.08E-06 |
| GO:0006357 | regulation of transcription from RNA polymerase II promoter | 1.41E-06 |
| GO:0045893 | positive regulation of transcription, DNA-templated | 1.22E-05 |
| GO:0030154 | cell differentiation | 1.67E-05 |
| GO:0016055 | Wnt signaling pathway | 1.91E-05 |
| GO:0021766 | hippocampus development | 4.51E-05 |
| GO:0010628 | positive regulation of gene expression | 5.57E-05 |
| GO:0000122 | negative regulation of transcription from RNA polymerase II promoter | 1.37E-04 |
| GO:0090090 | negative regulation of canonical Wnt signaling pathway | 2.21E-04 |
| GO:0035904 | aorta development | 3.56E-04 |
| GO:0006024 | glycosaminoglycan biosynthetic process | 4.45E-04 |
| GO:0021522 | spinal cord motor neuron differentiation | 4.94E-04 |
| GO:0007507 | heart development | 5.13E-04 |
| GO:0007224 | smoothened signaling pathway | 5.74E-04 |
| GO:0030182 | neuron differentiation | 6.07E-04 |
| GO:0043010 | camera-type eye development | 8.19E-04 |
| GO:0060021 | palate development | 0.00113011 |
| GO:0007281 | germ cell development | 0.00128033 |
| GO:0001843 | neural tube closure | 0.00184155 |
| GO:0002062 | chondrocyte differentiation | 0.00192196 |
| GO:0001657 | ureteric bud development | 0.00217299 |
| GO:0008284 | positive regulation of cell proliferation | 0.00252077 |
| GO:0001756 | somitogenesis | 0.00337659 |
| GO:0030111 | regulation of Wnt signaling pathway | 0.00412055 |
| GO:0015012 | heparan sulfate proteoglycan biosynthetic process | 0.00412055 |
| GO:0030324 | lung development | 0.00446791 |
| GO:0071300 | cellular response to retinoic acid | 0.00513915 |
| GO:0001701 | in utero embryonic development | 0.00546615 |
| GO:0045669 | positive regulation of osteoblast differentiation | 0.00582864 |
| GO:0001568 | blood vessel development | 0.00657098 |
| GO:0045165 | cell fate commitment | 0.00683035 |
| GO:0048844 | artery morphogenesis | 0.00788494 |
| GO:0045665 | negative regulation of neuron differentiation | 0.00792833 |
| GO:0030334 | regulation of cell migration | 0.00821812 |
| GO:0051216 | cartilage development | 0.00976019 |
| GO:0043066 | negative regulation of apoptotic process | 0.01109612 |
| GO:0021983 | pituitary gland development | 0.0112363 |
| GO:0001837 | epithelial to mesenchymal transition | 0.01196962 |
| GO:0007417 | central nervous system development | 0.01218445 |
| GO:0045892 | negative regulation of transcription, DNA-templated | 0.01259202 |
| GO:0035116 | embryonic hindlimb morphogenesis | 0.01349739 |
| GO:0010718 | positive regulation of epithelial to mesenchymal transition | 0.01429137 |
| GO:0007399 | nervous system development | 0.01594534 |
| GO:0061153 | trachea gland development | 0.01600806 |
| GO:0060976 | coronary vasculature development | 0.01679091 |
| GO:0001656 | metanephros development | 0.01766254 |
| GO:0003281 | ventricular septum development | 0.01766254 |
| GO:0071787 | endoplasmic reticulum tubular network assembly | 0.02128751 |
| GO:0034653 | retinoic acid catabolic process | 0.02128751 |
| GO:0008543 | fibroblast growth factor receptor signaling pathway | 0.02229888 |
| GO:0001649 | osteoblast differentiation | 0.02292059 |
| GO:0048706 | embryonic skeletal system development | 0.02328026 |
| GO:0002051 | osteoblast fate commitment | 0.02653893 |
| GO:0048755 | branching morphogenesis of a nerve | 0.02653893 |
| GO:0001755 | neural crest cell migration | 0.02844596 |
| GO:0001942 | hair follicle development | 0.03174493 |
| GO:0072177 | mesonephric duct development | 0.03176245 |
| GO:2000343 | positive regulation of chemokine (C-X-C motif) ligand 2 production | 0.03176245 |
| GO:0060662 | salivary gland cavitation | 0.03176245 |
| GO:0060789 | hair follicle placode formation | 0.03176245 |
| GO:0042127 | regulation of cell proliferation | 0.0339043 |
| GO:0060348 | bone development | 0.03636502 |
| GO:0009913 | epidermal cell differentiation | 0.03695824 |
| GO:1901522 | positive regulation of transcription from RNA polymerase II promoter involved in cellular response to chemical stimulus | 0.03695824 |
| GO:0021554 | optic nerve development | 0.03695824 |
| GO:0071168 | protein localization to chromatin | 0.03695824 |
| GO:0048752 | semicircular canal morphogenesis | 0.03695824 |
| GO:0048793 | pronephros development | 0.03695824 |
| GO:0048341 | paraxial mesoderm formation | 0.04212643 |
| GO:0048702 | embryonic neurocranium morphogenesis | 0.04212643 |
| GO:0030853 | negative regulation of granulocyte differentiation | 0.04212643 |
| GO:0001947 | heart looping | 0.04374438 |
| GO:0016567 | protein ubiquitination | 0.04533511 |

Supplementary Table S18. The top 100 enriched Uberon terms for the mouse orthologs of the pectoral fin module-specific genes.

| Uberon identifier | Term name | P-value |
| --- | --- | --- |
| uberon 0000922 | embryo | 6.92E-12 |
| uberon 0004716 | conceptus | 2.01E-11 |
| uberon 0007811 | craniocervical region | 8.10E-11 |
| uberon 0001049 | neural tube | 3.87E-10 |
| uberon 0005944 | axial skeleton plus cranial skeleton | 1.06E-09 |
| uberon 0003128 | cranium | 1.35E-09 |
| uberon 0000033 | head | 1.99E-09 |
| uberon 0001456 | face | 5.43E-08 |
| uberon 0001434 | skeletal system | 5.81E-07 |
| uberon 0000165 | mouth | 5.91E-07 |
| uberon 0001703 | neurocranium | 1.11E-06 |
| uberon 0001007 | digestive system | 2.10E-06 |
| uberon 0000926 | mesoderm | 2.55E-06 |
| uberon 0001711 | eyelid | 2.75E-06 |
| uberon 0004341 | primitive streak | 3.74E-06 |
| uberon 0011156 | facial skeleton | 5.55E-06 |
| uberon 0003457 | head bone | 7.51E-06 |
| uberon 0005070 | anterior neuropore | 1.28E-05 |
| uberon 0001819 | palpebral fissure | 2.93E-05 |
| uberon 0001708 | jaw skeleton | 2.93E-05 |
| uberon 0000478 | extraembryonic structure | 3.52E-05 |
| uberon 0000947 | aorta | 4.31E-05 |
| uberon 0003655 | molar tooth | 5.25E-05 |
| uberon 0004022 | germinal neuroepithelium | 7.58E-05 |
| uberon 0000948 | heart | 9.02E-05 |
| uberon 0000019 | camera-type eye | 9.15E-05 |
| uberon 0002104 | visual system | 9.22E-05 |
| uberon 0005600 | crus commune | 9.42E-05 |
| uberon 0001860 | endolymphatic duct | 0.0001457 |
| uberon 0003451 | lower jaw incisor | 0.00017578 |
| uberon 0002470 | autopod region | 0.00019574 |
| uberon 0001508 | arch of aorta | 0.0002267 |
| uberon 0002167 | right lung | 0.00023623 |
| uberon 0001004 | respiratory system | 0.0002535 |
| uberon 0001690 | ear | 0.00026285 |
| uberon 0002105 | vestibulo-auditory system | 0.00028019 |
| uberon 0003216 | hard palate | 0.00029064 |
| uberon 0010190 | pair of dorsal aortae | 0.00029282 |
| uberon 0002384 | connective tissue | 0.00030192 |
| uberon 0004044 | anterior visceral endoderm | 0.00043905 |
| uberon 0001818 | tarsal gland | 0.00049939 |
| uberon 0035077 | lateral nasal gland | 0.00062633 |
| uberon 0000014 | zone of skin | 0.0006311 |
| uberon 0002544 | digit | 0.00066645 |
| uberon 0002827 | vestibulocochlear ganglion | 0.00071149 |
| uberon 0003955 | molar crown | 0.0007931 |
| uberon 0001756 | middle ear | 0.00080729 |
| uberon 0004535 | cardiovascular system | 0.00087226 |
| uberon 0000423 | eccrine sweat gland | 0.00087311 |
| uberon 0001601 | extra-ocular muscle | 0.00087311 |
| uberon 0000004 | nose | 0.00090415 |
| uberon 0002539 | pharyngeal arch | 0.00091495 |
| uberon 0000084 | ureteric bud | 0.0009682 |
| uberon 0001751 | dentine | 0.00097362 |
| uberon 0003066 | pharyngeal arch 2 | 0.00097362 |
| uberon 0002168 | left lung | 0.00120309 |
| uberon 0001688 | incus bone | 0.00128991 |
| uberon 0000091 | bilaminar disc | 0.00140356 |
| uberon 0011864 | tendon collagen fibril | 0.00148402 |
| uberon 0001716 | secondary palate | 0.00151298 |
| uberon 0003544 | brain white matter | 0.0016391 |
| uberon 0000955 | brain | 0.00172361 |
| uberon 0003051 | ear vesicle | 0.00180251 |
| uberon 0003068 | axial mesoderm | 0.00184711 |
| uberon 0007833 | osseus semicircular canal | 0.00187241 |
| uberon 0000309 | body wall | 0.00193054 |
| uberon 0002091 | appendicular skeleton | 0.00207267 |
| uberon 0004573 | systemic artery | 0.00222527 |
| uberon 0010513 | strand of zigzag hair | 0.00225969 |
| uberon 0000945 | stomach | 0.0023366 |
| uberon 0010409 | ocular surface region | 0.00233819 |
| uberon 0004043 | semicircular canal ampulla | 0.00242659 |
| uberon 0005356 | rathke's pouch | 0.00242659 |
| uberon 0003950 | inner ear canal | 0.00254497 |
| uberon 0001890 | forebrain | 0.00257106 |
| uberon 0003252 | thoracic rib cage | 0.00267041 |
| uberon 0008854 | root of molar tooth | 0.00268605 |
| uberon 0008799 | transverse palatine fold | 0.00268605 |
| uberon 0002418 | cartilage tissue | 0.00271496 |
| uberon 0001199 | mucosa of stomach | 0.00287821 |
| uberon 0005291 | embryonic tissue | 0.00300689 |
| uberon 0001167 | wall of stomach | 0.00324016 |
| uberon 0000007 | pituitary gland | 0.00332109 |
| uberon 0001862 | vestibular labyrinth | 0.003632 |
| uberon 0002413 | cervical vertebra | 0.00380992 |
| uberon 0004362 | pharyngeal arch 1 | 0.00403845 |
| uberon 0001872 | parietal lobe | 0.00403845 |
| uberon 0002329 | somite | 0.00405486 |
| uberon 0001908 | optic tract | 0.00421892 |
| uberon 0001274 | ischium | 0.00421892 |
| uberon 0001853 | utricle of membranous labyrinth | 0.00477616 |
| uberon 0001752 | enamel | 0.00477616 |
| uberon 0002196 | adenohypophysis | 0.00516843 |
| uberon 0005176 | tooth enamel organ | 0.00541814 |
| uberon 0002487 | tooth cavity | 0.00541814 |
| uberon 0010197 | trunk of common carotid artery | 0.00541814 |
| uberon 0004090 | periorbital region | 0.00541814 |
| uberon 0001894 | diencephalon | 0.00586178 |
| uberon 0001854 | saccule of membranous labyrinth | 0.00617638 |
| uberon 0004212 | glomerular capillary | 0.00617638 |

Supplementary Table S19. The top 100 enriched Biological Process terms from the Gene Ontology for the mouse orthologs of the pelvic fin module-specific genes.

| Term identifier | Term name | P-value |
| --- | --- | --- |
| GO:0006468 | protein phosphorylation | 5.19E-34 |
| GO:0016310 | phosphorylation | 1.24E-30 |
| GO:0046777 | protein autophosphorylation | 9.64E-22 |
| GO:0018105 | peptidyl-serine phosphorylation | 1.63E-15 |
| GO:0045893 | positive regulation of transcription, DNA-templated | 5.38E-15 |
| GO:0018108 | peptidyl-tyrosine phosphorylation | 1.80E-14 |
| GO:0045944 | positive regulation of transcription from RNA polymerase II promoter | 3.24E-14 |
| GO:0007169 | transmembrane receptor protein tyrosine kinase signaling pathway | 1.41E-13 |
| GO:0010628 | positive regulation of gene expression | 1.67E-13 |
| GO:0009887 | organ morphogenesis | 7.65E-12 |
| GO:0030335 | positive regulation of cell migration | 1.14E-11 |
| GO:0042493 | response to drug | 7.06E-11 |
| GO:0018107 | peptidyl-threonine phosphorylation | 1.51E-10 |
| GO:0043066 | negative regulation of apoptotic process | 4.82E-10 |
| GO:0043627 | response to estrogen | 5.96E-10 |
| GO:0001934 | positive regulation of protein phosphorylation | 6.88E-10 |
| GO:0006351 | transcription, DNA-templated | 1.09E-09 |
| GO:0038083 | peptidyl-tyrosine autophosphorylation | 1.75E-09 |
| GO:0048565 | digestive tract development | 2.26E-09 |
| GO:0035690 | cellular response to drug | 7.15E-09 |
| GO:0008284 | positive regulation of cell proliferation | 9.34E-09 |
| GO:0035556 | intracellular signal transduction | 1.19E-08 |
| GO:0030154 | cell differentiation | 1.47E-08 |
| GO:0035970 | peptidyl-threonine dephosphorylation | 2.43E-08 |
| GO:0007275 | multicellular organism development | 4.69E-08 |
| GO:0045665 | negative regulation of neuron differentiation | 8.95E-08 |
| GO:0071773 | cellular response to BMP stimulus | 1.34E-07 |
| GO:0007264 | small GTPase mediated signal transduction | 2.17E-07 |
| GO:0000122 | negative regulation of transcription from RNA polymerase II promoter | 2.52E-07 |
| GO:0030182 | neuron differentiation | 3.94E-07 |
| GO:0060070 | canonical Wnt signaling pathway | 4.66E-07 |
| GO:0071407 | cellular response to organic cyclic compound | 7.62E-07 |
| GO:0016477 | cell migration | 1.56E-06 |
| GO:0006355 | regulation of transcription, DNA-templated | 1.98E-06 |
| GO:0007178 | transmembrane receptor protein serine/threonine kinase signaling pathway | 2.25E-06 |
| GO:0001942 | hair follicle development | 3.29E-06 |
| GO:0043525 | positive regulation of neuron apoptotic process | 3.31E-06 |
| GO:0007049 | cell cycle | 4.64E-06 |
| GO:0043524 | negative regulation of neuron apoptotic process | 5.40E-06 |
| GO:0007179 | transforming growth factor beta receptor signaling pathway | 6.68E-06 |
| GO:0006357 | regulation of transcription from RNA polymerase II promoter | 8.40E-06 |
| GO:0048485 | sympathetic nervous system development | 8.61E-06 |
| GO:0045740 | positive regulation of DNA replication | 9.30E-06 |
| GO:0071363 | cellular response to growth factor stimulus | 1.61E-05 |
| GO:0007507 | heart development | 1.70E-05 |
| GO:0048663 | neuron fate commitment | 1.77E-05 |
| GO:0030509 | BMP signaling pathway | 2.27E-05 |
| GO:0001701 | in utero embryonic development | 2.45E-05 |
| GO:0008584 | male gonad development | 2.93E-05 |
| GO:0006470 | protein dephosphorylation | 3.41E-05 |
| GO:0071333 | cellular response to glucose stimulus | 3.48E-05 |
| GO:0030513 | positive regulation of BMP signaling pathway | 3.59E-05 |
| GO:0031175 | neuron projection development | 3.65E-05 |
| GO:0045165 | cell fate commitment | 3.86E-05 |
| GO:0003151 | outflow tract morphogenesis | 4.33E-05 |
| GO:0043406 | positive regulation of MAP kinase activity | 4.33E-05 |
| GO:0042127 | regulation of cell proliferation | 5.34E-05 |
| GO:0010629 | negative regulation of gene expression | 7.71E-05 |
| GO:0016569 | covalent chromatin modification | 8.05E-05 |
| GO:0051496 | positive regulation of stress fiber assembly | 1.15E-04 |
| GO:0060041 | retina development in camera-type eye | 1.17E-04 |
| GO:0010468 | regulation of gene expression | 1.23E-04 |
| GO:0003007 | heart morphogenesis | 1.31E-04 |
| GO:0045429 | positive regulation of nitric oxide biosynthetic process | 1.31E-04 |
| GO:0007399 | nervous system development | 1.38E-04 |
| GO:0045597 | positive regulation of cell differentiation | 1.49E-04 |
| GO:0006366 | transcription from RNA polymerase II promoter | 1.71E-04 |
| GO:0008285 | negative regulation of cell proliferation | 1.73E-04 |
| GO:0007165 | signal transduction | 2.05E-04 |
| GO:0045931 | positive regulation of mitotic cell cycle | 2.26E-04 |
| GO:0032212 | positive regulation of telomere maintenance via telomerase | 2.26E-04 |
| GO:0045471 | response to ethanol | 2.30E-04 |
| GO:0007224 | smoothened signaling pathway | 2.52E-04 |
| GO:0007167 | enzyme linked receptor protein signaling pathway | 2.85E-04 |
| GO:0007204 | positive regulation of cytosolic calcium ion concentration | 3.02E-04 |
| GO:0048709 | oligodendrocyte differentiation | 3.03E-04 |
| GO:0001525 | angiogenesis | 3.37E-04 |
| GO:0030324 | lung development | 3.55E-04 |
| GO:0001764 | neuron migration | 3.55E-04 |
| GO:0048015 | phosphatidylinositol-mediated signaling | 3.99E-04 |
| GO:0048538 | thymus development | 4.03E-04 |
| GO:0016055 | Wnt signaling pathway | 4.15E-04 |
| GO:0090090 | negative regulation of canonical Wnt signaling pathway | 4.48E-04 |
| GO:0010001 | glial cell differentiation | 4.49E-04 |
| GO:0071260 | cellular response to mechanical stimulus | 4.50E-04 |
| GO:0030501 | positive regulation of bone mineralization | 4.54E-04 |
| GO:0009790 | embryo development | 4.87E-04 |
| GO:0070301 | cellular response to hydrogen peroxide | 4.90E-04 |
| GO:0045909 | positive regulation of vasodilation | 5.16E-04 |
| GO:0060412 | ventricular septum morphogenesis | 5.16E-04 |
| GO:0070374 | positive regulation of ERK1 and ERK2 cascade | 5.21E-04 |
| GO:0048661 | positive regulation of smooth muscle cell proliferation | 5.26E-04 |
| GO:0008283 | cell proliferation | 5.53E-04 |
| GO:0051091 | positive regulation of sequence-specific DNA binding transcription factor activity | 6.17E-04 |
| GO:0001666 | response to hypoxia | 6.21E-04 |
| GO:0060045 | positive regulation of cardiac muscle cell proliferation | 6.46E-04 |
| GO:0007623 | circadian rhythm | 6.56E-04 |
| GO:0048646 | anatomical structure formation involved in morphogenesis | 7.64E-04 |
| GO:0048854 | brain morphogenesis | 7.64E-04 |
| GO:0000082 | G1/S transition of mitotic cell cycle | 7.71E-04 |

Supplementary Table S20. The top 100 Uberon terms for the mouse orthologs of the pelvic fin module-specific genes.

| Uberon identifier | Term name | P-value |
| --- | --- | --- |
| uberon 0000922 | embryo | 6.50E-16 |
| uberon 0004716 | conceptus | 1.53E-13 |
| uberon 0004365 | vitelline blood vessel | 2.69E-09 |
| uberon 0000478 | extraembryonic structure | 3.51E-09 |
| uberon 0001007 | digestive system | 8.53E-09 |
| uberon 0000014 | zone of skin | 2.47E-08 |
| uberon 0010190 | pair of dorsal aortae | 4.77E-08 |
| uberon 0001049 | neural tube | 1.02E-07 |
| uberon 0008852 | visceral yolk sac | 1.23E-07 |
| uberon 0000948 | heart | 1.60E-07 |
| uberon 0004535 | cardiovascular system | 4.34E-07 |
| uberon 0000468 | multicellular organism | 5.93E-07 |
| uberon 0000016 | endocrine pancreas | 8.87E-07 |
| uberon 0002067 | dermis | 1.91E-06 |
| uberon 0001004 | respiratory system | 3.10E-06 |
| uberon 0000358 | blastocyst | 3.19E-06 |
| uberon 0000033 | head | 3.57E-06 |
| uberon 0002012 | pulmonary artery | 4.49E-06 |
| uberon 0002407 | pericardium | 1.34E-05 |
| uberon 0002048 | lung | 1.66E-05 |
| uberon 0004374 | vitelline vasculature | 2.27E-05 |
| uberon 0001987 | placenta | 2.69E-05 |
| uberon 0003087 | anterior cardinal vein | 3.00E-05 |
| uberon 0000006 | islet of langerhans | 3.49E-05 |
| uberon 0001264 | pancreas | 4.05E-05 |
| uberon 0000947 | aorta | 4.50E-05 |
| uberon 0001456 | face | 6.01E-05 |
| uberon 0003512 | lung blood vessel | 6.31E-05 |
| uberon 0002240 | spinal cord | 7.35E-05 |
| uberon 0003946 | placenta labyrinth | 0.00010402 |
| uberon 0002368 | endocrine gland | 0.00011026 |
| uberon 0001792 | ganglionic layer of retina | 0.00012935 |
| uberon 0002062 | endocardial cushion | 0.00014493 |
| uberon 0001508 | arch of aorta | 0.00015353 |
| uberon 0001083 | myocardium of ventricle | 0.0001601 |
| uberon 0007811 | craniocervical region | 0.00016979 |
| uberon 0001809 | enteric ganglion | 0.0001807 |
| uberon 0000084 | ureteric bud | 0.00020974 |
| uberon 0002087 | atrioventricular canal | 0.00022946 |
| uberon 0002315 | gray matter of spinal cord | 0.00024038 |
| uberon 0004647 | liver lobule | 0.00026764 |
| uberon 0001818 | tarsal gland | 0.00026935 |
| uberon 0000165 | mouth | 0.00027721 |
| uberon 0001675 | trigeminal ganglion | 0.00027999 |
| uberon 0006207 | aortico-pulmonary spiral septum | 0.00028524 |
| uberon 0002416 | integumental system | 0.00028781 |
| uberon 0010172 | bulb of aorta | 0.00030071 |
| uberon 0002005 | enteric nervous system | 0.00030515 |
| uberon 0006524 | alveolar system | 0.00031454 |
| uberon 0008870 | pulmonary alveolar parenchyma | 0.00034866 |
| uberon 0000088 | trophoblast | 0.00039593 |
| uberon 0002094 | interventricular septum | 0.000402 |
| uberon 0000383 | musculature of body | 0.00042772 |
| uberon 0001711 | eyelid | 0.00043278 |
| uberon 0003216 | hard palate | 0.00044958 |
| uberon 0001280 | liver parenchyma | 0.00046467 |
| uberon 0000011 | parasympathetic nervous system | 0.0004649 |
| uberon 0002073 | hair follicle | 0.00052472 |
| uberon 0002370 | thymus | 0.00057905 |
| uberon 0008856 | stomach muscularis externa | 0.0006099 |
| uberon 0000019 | camera-type eye | 0.00061682 |
| uberon 0002408 | parietal serous pericardium | 0.00062547 |
| uberon 0001496 | ascending aorta | 0.00063348 |
| uberon 0000117 | respiratory tube | 0.00063848 |
| uberon 0002165 | endocardium | 0.00073687 |
| uberon 0003066 | pharyngeal arch 2 | 0.00073687 |
| uberon 0002511 | trabecula carnea | 0.00077776 |
| uberon 0000013 | sympathetic nervous system | 0.0008281 |
| uberon 0000012 | somatic nervous system | 0.00090485 |
| uberon 0002342 | neural crest | 0.00093289 |
| uberon 0003618 | aorta tunica media | 0.00093289 |
| uberon 0002080 | heart right ventricle | 0.00100621 |
| uberon 0001637 | artery | 0.0010228 |
| uberon 0001708 | jaw skeleton | 0.00113097 |
| uberon 0002104 | visual system | 0.00120939 |
| uberon 0002069 | stratum granulosum of epidermis | 0.00121228 |
| uberon 0001074 | pericardial cavity | 0.00142689 |
| uberon 0001041 | foregut | 0.00145277 |
| uberon 0008874 | pulmonary acinus | 0.00148651 |
| uberon 0004493 | cardiac muscle tissue of myocardium | 0.00151061 |
| uberon 0002410 | autonomic nervous system | 0.00154176 |
| uberon 0001135 | smooth muscle tissue | 0.00168825 |
| uberon 0001716 | secondary palate | 0.00180283 |
| uberon 0003501 | retina blood vessel | 0.00195196 |
| uberon 0002025 | stratum basale of epidermis | 0.00203946 |
| uberon 0011156 | facial skeleton | 0.0021246 |
| uberon 0003823 | hindlimb zeugopod | 0.00219172 |
| uberon 0001534 | right subclavian artery | 0.00222747 |
| uberon 0005970 | brain commissure | 0.00223824 |
| uberon 0003073 | lens placode | 0.00223824 |
| uberon 0003128 | cranium | 0.00240653 |
| uberon 0001003 | skin epidermis | 0.0024443 |
| uberon 0010513 | strand of zigzag hair | 0.00258297 |
| uberon 0001213 | intestinal villus | 0.00269833 |
| uberon 0001806 | sympathetic ganglion | 0.00286072 |
| uberon 0005870 | olfactory pit | 0.00286072 |
| uberon 0005343 | cortical plate | 0.00286898 |
| uberon 0005062 | neural fold | 0.00286898 |
| uberon 0004663 | aorta wall | 0.0028703 |
| uberon 0010512 | strand of guard hair | 0.00311381 |

Supplementary Table S21. Some of the enriched GO-BP and Uberon terms for the mouse orthologs of the pectoral fin module-specific genes that are related to novel anatomical entities emerged in tetrapods during fin to limb transition. The terms are organized into specific anatomical regions.

| Anatomical region | Term Identifier | Term name | P-value |
| --- | --- | --- | --- |
| Lung | uberon 0002167 | right lung | 0.00023623 |
|  | uberon 0002168 | left lung | 0.00120309 |
|  | uberon 0003512 | lung blood vessel | 0.04982657 |
|  | GO:0030324 | lung development | 0.00446791 |
| Neck and lower jaw | uberon 0001708 | jaw skeleton | 2.93E-05 |
|  | uberon 0003451 | lower jaw incisor | 0.00017578 |
|  | uberon 0002413 | cervical vertebra | 0.00380992 |
|  | uberon 0003216 | hard palate | 0.00029064 |
|  | uberon 0001716 | secondary palate | 0.00151298 |
|  | GO:0060021 | palate development | 0.00113011 |
|  | GO:0061153 | trachea gland development | 0.01600806 |
| Face and hair | uberon 0001456 | face | 5.43E-08 |
|  | uberon 0005600 | crus commune | 9.42E-05 |
|  | uberon 0001690 | ear | 0.00026285 |
|  | uberon 0000004 | nose | 0.00090415 |
|  | uberon 0001711 | eyelid | 2.75E-06 |
|  | uberon 0034772 | margin of eyelid | 0.03232832 |
|  | uberon 0002073 | hair follicle | 0.02808325 |
|  | uberon 0010514 | strand of duvet hair | 0.02594653 |
|  | GO:0001942 | hair follicle development | 0.03174493 |
|  | GO:0060789 | hair follicle placode formation | 0.03176245 |
|  | GO:0061029 | eyelid development in camera-type eye | 0.07754317 |
| Other | uberon 0002544 | digit | 0.00066645 |

Supplementary Table S22. Some of the enriched GO-BP and Uberon terms for the mouse orthologs of the pelvic fin module-specific genes that are related to novel anatomical entities emerged in tetrapods during fin to limb transition. The terms are organized into specific anatomical regions.

| Anatomical region | Term Identifier | Term name | P-value |
| --- | --- | --- | --- |
| Lung | uberon 0001004 | respiratory system | 3.10E-06 |
|  | uberon 0002012 | pulmonary artery | 4.49E-06 |
|  | uberon 0002048 | lung | 1.66E-05 |
|  | uberon 0003512 | lung blood vessel | 6.31E-05 |
|  | uberon 0006524 | alveolar system | 0.00031454 |
|  | uberon 0008870 | pulmonary alveolar parenchyma | 0.00034866 |
|  | uberon 0000117 | respiratory tube | 0.00063848 |
|  | uberon 0004785 | respiratory system mucosa | 0.00465107 |
|  | GO:0030324 | lung development | 3.55E-04 |
|  | GO:0060425 | lung morphogenesis | 0.0083659 |
| Neck and lower jaw | uberon 0003216 | hard palate | 0.00044958 |
|  | uberon 0002370 | thymus | 0.00057905 |
|  | GO:0048538 | thymus development | 4.03E-04 |
|  | GO:0060021 | palate development | 0.00391519 |
|  | GO:0060440 | trachea formation | 0.00614484 |
|  | GO:0060017 | parathyroid gland development | 0.00614484 |
| Face and hair | uberon 0001818 | tarsal gland | 0.00026935 |
|  | uberon 0001711 | eyelid | 0.00043278 |
|  | uberon 0000004 | nose | 0.00707171 |
|  | uberon 0001681 | nasal bone | 0.01712347 |
|  | uberon 0002073 | hair follicle | 0.00052472 |
|  | uberon 0010512 | strand of guard hair | 0.00311381 |
|  | GO:0061029 | eyelid development in camera-type eye | 0.00211434 |
|  | GO:0001942 | hair follicle development | 3.29E-06 |
| Other | uberon 0001987 | placenta | 2.69E-05 |
|  | uberon 0003946 | placenta labyrinth | 0.00010402 |

Supplementary Table S23. The enriched Biological Process terms from the Gene Ontology for the zebrafish orthologs of the forelimb module-specific genes.

| Term identifier | Term name | P-value |
| --- | --- | --- |
| GO:0006355 | regulation of transcription, DNA-templated | 2.74E-11 |
| GO:0007275 | multicellular organism development | 2.01E-09 |
| GO:0009953 | dorsal/ventral pattern formation | 1.36E-08 |
| GO:0016055 | Wnt signaling pathway | 6.90E-08 |
| GO:0006351 | transcription, DNA-templated | 2.18E-07 |
| GO:0030509 | BMP signaling pathway | 5.94E-07 |
| GO:0030182 | neuron differentiation | 7.04E-07 |
| GO:0035118 | embryonic pectoral fin morphogenesis | 1.60E-05 |
| GO:0045165 | cell fate commitment | 5.38E-05 |
| GO:0030510 | regulation of BMP signaling pathway | 1.57E-04 |
| GO:0048701 | embryonic cranial skeleton morphogenesis | 4.43E-04 |
| GO:0002072 | optic cup morphogenesis involved in camera-type eye development | 4.70E-04 |
| GO:0007178 | transmembrane receptor protein serine/threonine kinase signaling pathway | 0.00117724 |
| GO:0040007 | growth | 0.00129193 |
| GO:0060027 | convergent extension involved in gastrulation | 0.00135252 |
| GO:0010862 | positive regulation of pathway-restricted SMAD protein phosphorylation | 0.0015685 |
| GO:0042074 | cell migration involved in gastrulation | 0.00165586 |
| GO:0048468 | cell development | 0.00172114 |
| GO:0060395 | SMAD protein signal transduction | 0.00188379 |
| GO:0030903 | notochord development | 0.00205679 |
| GO:0043408 | regulation of MAPK cascade | 0.00205679 |
| GO:0007179 | transforming growth factor beta receptor signaling pathway | 0.00414164 |
| GO:0021592 | fourth ventricle development | 0.00421122 |
| GO:0045743 | positive regulation of fibroblast growth factor receptor signaling pathway | 0.00536967 |
| GO:0048703 | embryonic viscerocranium morphogenesis | 0.00585852 |
| GO:0060070 | canonical Wnt signaling pathway | 0.0061092 |
| GO:0001944 | vasculature development | 0.00649015 |
| GO:0015012 | heparan sulfate proteoglycan biosynthetic process | 0.00665665 |
| GO:0002062 | chondrocyte differentiation | 0.00665665 |
| GO:0030097 | hemopoiesis | 0.00670985 |
| GO:0008543 | fibroblast growth factor receptor signaling pathway | 0.0095973 |
| GO:0007417 | central nervous system development | 0.01064297 |
| GO:0061386 | closure of optic fissure | 0.01302316 |
| GO:0006024 | glycosaminoglycan biosynthetic process | 0.01302316 |
| GO:0071910 | determination of liver left/right asymmetry | 0.01490327 |
| GO:0007064 | mitotic sister chromatid cohesion | 0.01689251 |
| GO:0009948 | anterior/posterior axis specification | 0.01898788 |
| GO:0051216 | cartilage development | 0.01918023 |
| GO:0007155 | cell adhesion | 0.01955582 |
| GO:0061371 | determination of heart left/right asymmetry | 0.02012978 |
| GO:0045595 | regulation of cell differentiation | 0.02118641 |
| GO:0030198 | extracellular matrix organization | 0.0234852 |
| GO:0060026 | convergent extension | 0.02377358 |
| GO:0038108 | negative regulation of appetite by leptin-mediated signaling pathway | 0.02505648 |
| GO:2000366 | positive regulation of STAT protein import into nucleus | 0.02505648 |
| GO:0060912 | cardiac cell fate specification | 0.02505648 |
| GO:1900745 | positive regulation of p38MAPK cascade | 0.02505648 |
| GO:2000583 | regulation of platelet-derived growth factor receptor-alpha signaling pathway | 0.02505648 |
| GO:0002076 | osteoblast development | 0.02505648 |
| GO:0070587 | regulation of cell-cell adhesion involved in gastrulation | 0.02505648 |
| GO:1990051 | activation of protein kinase C activity | 0.02505648 |
| GO:0043010 | camera-type eye development | 0.02506223 |
| GO:0060041 | retina development in camera-type eye | 0.02513516 |
| GO:0060828 | regulation of canonical Wnt signaling pathway | 0.02588139 |
| GO:0001503 | ossification | 0.02588139 |
| GO:0060037 | pharyngeal system development | 0.02837219 |
| GO:0050767 | regulation of neurogenesis | 0.02837219 |
| GO:0002138 | retinoic acid biosynthetic process | 0.03734953 |
| GO:0032481 | positive regulation of type I interferon production | 0.03734953 |
| GO:0010332 | response to gamma radiation | 0.03734953 |
| GO:0032965 | regulation of collagen biosynthetic process | 0.03734953 |
| GO:0006978 | DNA damage response, signal transduction by p53 class mediator resulting in transcription of p21 class mediator | 0.03734953 |
| GO:0010470 | regulation of gastrulation | 0.03734953 |
| GO:0071733 | transcriptional activation by promoter-enhancer looping | 0.03734953 |
| GO:0030917 | midbrain-hindbrain boundary development | 0.04515412 |
| GO:0001568 | blood vessel development | 0.04877064 |
| GO:0048730 | epidermis morphogenesis | 0.0494884 |
| GO:0032868 | response to insulin | 0.0494884 |

Supplementary Table S24. The enriched Uberon terms for the zebrafish orthologs of the forelimb module-specific genes.

| Uberon identifier | Term name | P-value |
| --- | --- | --- |
| uberon 0008823 | neural tube derived brain | 2.29E-07 |
| uberon 0000033 | head | 2.96E-07 |
| uberon 0000468 | multicellular organism | 3.10E-07 |
| uberon 0000019 | camera-type eye | 2.07E-05 |
| uberon 0001555 | digestive tract | 9.84E-05 |
| uberon 0001016 | nervous system | 0.00082865 |
| uberon 0002107 | liver | 0.00093463 |
| uberon 0001017 | central nervous system | 0.00141338 |
| uberon 2000084 | yolk | 0.00145089 |
| uberon 4000164 | caudal fin | 0.00325469 |
| uberon 0000948 | heart | 0.00876155 |
| uberon 0008895 | splanchnocranium | 0.01593721 |
| uberon 0002280 | otolith | 0.01730733 |
| uberon 0005281 | ventricular system of central nervous system | 0.02130098 |
| uberon 2000033 | intermediate cell mass of mesoderm | 0.02183289 |
| uberon 0001782 | pigmented layer of retina | 0.02256819 |
| uberon 0001231 | nephron tubule | 0.02976965 |
| uberon 0001155 | colon | 0.02976965 |
| uberon 0001708 | jaw skeleton | 0.0342243 |
| uberon 0005886 | post-hyoid pharyngeal arch skeleton | 0.03545473 |
| uberon 0001647 | facial nerve | 0.03865184 |
| uberon 0002407 | pericardium | 0.04825439 |

Supplementary Table S25. The enriched Biological Process terms from the Gene Ontology for the zebrafish orthologs of the hindlimb module-specific genes.

| Term identifier | Term name | P-value |
| --- | --- | --- |
| GO:0006355 | regulation of transcription, DNA-templated | 1.34E-09 |
| GO:0007275 | multicellular organism development | 5.81E-09 |
| GO:0040007 | growth | 7.61E-07 |
| GO:0010862 | positive regulation of pathway-restricted SMAD protein phosphorylation | 1.22E-06 |
| GO:0060395 | SMAD protein signal transduction | 1.89E-06 |
| GO:0043408 | regulation of MAPK cascade | 2.33E-06 |
| GO:0048701 | embryonic cranial skeleton morphogenesis | 1.03E-05 |
| GO:0048468 | cell development | 1.49E-05 |
| GO:0051216 | cartilage development | 1.50E-05 |
| GO:0030182 | neuron differentiation | 3.49E-05 |
| GO:0016055 | Wnt signaling pathway | 5.61E-05 |
| GO:0007417 | central nervous system development | 1.10E-04 |
| GO:0060070 | canonical Wnt signaling pathway | 1.97E-04 |
| GO:0006351 | transcription, DNA-templated | 2.70E-04 |
| GO:0008543 | fibroblast growth factor receptor signaling pathway | 3.76E-04 |
| GO:0001501 | skeletal system development | 5.14E-04 |
| GO:0007155 | cell adhesion | 6.49E-04 |
| GO:0048703 | embryonic viscerocranium morphogenesis | 9.92E-04 |
| GO:0043401 | steroid hormone mediated signaling pathway | 0.00114721 |
| GO:0030903 | notochord development | 0.00117063 |
| GO:0005975 | carbohydrate metabolic process | 0.00129702 |
| GO:0009612 | response to mechanical stimulus | 0.00135076 |
| GO:0030510 | regulation of BMP signaling pathway | 0.00160837 |
| GO:0030513 | positive regulation of BMP signaling pathway | 0.00204619 |
| GO:0001503 | ossification | 0.00204619 |
| GO:0045165 | cell fate commitment | 0.00207662 |
| GO:0000165 | MAPK cascade | 0.00225021 |
| GO:0002062 | chondrocyte differentiation | 0.00225021 |
| GO:0007626 | locomotory behavior | 0.0023328 |
| GO:0031101 | fin regeneration | 0.00239174 |
| GO:0043010 | camera-type eye development | 0.0025896 |
| GO:0007517 | muscle organ development | 0.00308006 |
| GO:0009953 | dorsal/ventral pattern formation | 0.00369912 |
| GO:0005978 | glycogen biosynthetic process | 0.0039571 |
| GO:0030916 | otic vesicle formation | 0.0039571 |
| GO:0035118 | embryonic pectoral fin morphogenesis | 0.00553237 |
| GO:0007179 | transforming growth factor beta receptor signaling pathway | 0.00553237 |
| GO:0030282 | bone mineralization | 0.00628086 |
| GO:0060349 | bone morphogenesis | 0.00737634 |
| GO:0030902 | hindbrain development | 0.00739625 |
| GO:0070654 | sensory epithelium regeneration | 0.00769009 |
| GO:0031099 | regeneration | 0.00927093 |
| GO:0007519 | skeletal muscle tissue development | 0.0106517 |
| GO:0043627 | response to estrogen | 0.01086062 |
| GO:0061024 | membrane organization | 0.01086062 |
| GO:0021587 | cerebellum morphogenesis | 0.01086062 |
| GO:0043697 | cell dedifferentiation | 0.01086062 |
| GO:0009948 | anterior/posterior axis specification | 0.01102739 |
| GO:0042074 | cell migration involved in gastrulation | 0.01173397 |
| GO:0045595 | regulation of cell differentiation | 0.01296272 |
| GO:0048839 | inner ear development | 0.01440797 |
| GO:0008045 | motor neuron axon guidance | 0.01440797 |
| GO:0061300 | cerebellum vasculature development | 0.01492536 |
| GO:0030198 | extracellular matrix organization | 0.01507944 |
| GO:0060536 | cartilage morphogenesis | 0.01737942 |
| GO:0060021 | palate development | 0.01953552 |
| GO:0046716 | muscle cell cellular homeostasis | 0.01953552 |
| GO:0060538 | skeletal muscle organ development | 0.01953552 |
| GO:0048884 | neuromast development | 0.01986392 |
| GO:0060037 | pharyngeal system development | 0.01986392 |
| GO:0035567 | non-canonical Wnt signaling pathway | 0.02253359 |
| GO:0042981 | regulation of apoptotic process | 0.02458938 |
| GO:0072661 | protein targeting to plasma membrane | 0.02465756 |
| GO:0006874 | cellular calcium ion homeostasis | 0.02703075 |
| GO:0055113 | epiboly involved in gastrulation with mouth forming second | 0.02923793 |
| GO:0016203 | muscle attachment | 0.0302594 |
| GO:0030500 | regulation of bone mineralization | 0.0302594 |
| GO:0048840 | otolith development | 0.03165263 |
| GO:0007267 | cell-cell signaling | 0.03311103 |
| GO:0001889 | liver development | 0.03497684 |
| GO:0051482 | positive regulation of cytosolic calcium ion concentration involved in phospholipase C-activating G-protein coupled signaling pathway | 0.03505962 |
| GO:0060027 | convergent extension involved in gastrulation | 0.03507928 |
| GO:0001568 | blood vessel development | 0.03507928 |
| GO:0055001 | muscle cell development | 0.03631032 |
| GO:0030199 | collagen fibril organization | 0.03631032 |
| GO:0060041 | retina development in camera-type eye | 0.03668877 |
| GO:0006357 | regulation of transcription from RNA polymerase II promoter | 0.03680213 |
| GO:0060059 | embryonic retina morphogenesis in camera-type eye | 0.03864788 |
| GO:0045944 | positive regulation of transcription from RNA polymerase II promoter | 0.0397757 |
| GO:0006508 | proteolysis | 0.04168701 |
| GO:0007631 | feeding behavior | 0.04278097 |

Supplementary Table S26. The enriched Uberon terms for the zebrafish orthologs of the hindlimb module-specific genes.

| Uberon identifier | Term name | P-value |
| --- | --- | --- |
| uberon 0000019 | camera-type eye | 7.22E-18 |
| uberon 0000468 | multicellular organism | 5.69E-17 |
| uberon 0000033 | head | 2.90E-15 |
| uberon 0008823 | neural tube derived brain | 1.67E-12 |
| uberon 0001016 | nervous system | 5.42E-10 |
| uberon 0001017 | central nervous system | 1.81E-08 |
| uberon 0002107 | liver | 4.19E-07 |
| uberon 0000948 | heart | 6.07E-07 |
| uberon 0001555 | digestive tract | 1.70E-06 |
| uberon 0001032 | sensory system | 1.93E-06 |
| uberon 0007812 | post-anal tail | 2.28E-06 |
| uberon 0002407 | pericardium | 1.47E-05 |
| uberon 0005886 | post-hyoid pharyngeal arch skeleton | 3.02E-05 |
| uberon 2000084 | yolk | 3.05E-05 |
| uberon 0005281 | ventricular system of central nervous system | 6.07E-05 |
| uberon 0010314 | structure with developmental contribution from neural crest | 0.00074386 |
| uberon 0001846 | internal ear | 0.0008888 |
| uberon 0002329 | somite | 0.00089648 |
| uberon 0001003 | skin epidermis | 0.00170376 |
| uberon 0003102 | surface structure | 0.00190061 |
| uberon 0002100 | trunk | 0.00252997 |
| uberon 0001708 | jaw skeleton | 0.00287797 |
| uberon 0004141 | heart tube | 0.00345967 |
| uberon 0001782 | pigmented layer of retina | 0.00380629 |
| uberon 0000017 | exocrine pancreas | 0.00384968 |
| uberon 4000164 | caudal fin | 0.0046263 |
| uberon 0001945 | superior colliculus | 0.00488419 |
| uberon 0012438 | blastema of regenerating fin/limb | 0.00591299 |
| uberon 0000965 | lens of camera-type eye | 0.00652003 |
| uberon 0000966 | retina | 0.0085354 |
| uberon 0002280 | otolith | 0.01008715 |
| uberon 0008897 | fin | 0.01130278 |
| uberon 0002028 | hindbrain | 0.01447067 |
| uberon 0001890 | forebrain | 0.01678628 |
| uberon 0014907 | intersomitic vessel | 0.01866955 |
| uberon 0005884 | hyoid arch skeleton | 0.01978595 |
| uberon 2000106 | extension | 0.02390583 |
| uberon 0005310 | pronephric nephron tubule | 0.02958931 |
| uberon 2001456 | pectoral fin endoskeletal disc | 0.0328624 |
| uberon 0011611 | ceratohyal bone | 0.0328624 |
| uberon 0003052 | midbrain-hindbrain boundary | 0.03515936 |
| uberon 0002328 | notochord | 0.03668385 |
| uberon 0002082 | cardiac ventricle | 0.03785239 |
| uberon 0006283 | future cardiac ventricle | 0.04193983 |
| uberon 0005087 | tooth placode | 0.04412975 |
| uberon 0002422 | fourth ventricle | 0.04604822 |

Supplementary Table S27. Some of the enriched GO-BP and Uberon terms that are related to fin to limb transition for the zebrafish orthologs of the forelimb module-specific genes. The terms are organized into specific anatomical regions.

| Anatomical region | Term Identifier | Term name | P-value |
| --- | --- | --- | --- |
| Fin related | uberon 4000164 | caudal fin | 0.00325469 |
|  | GO:0035118 | embryonic pectoral fin morphogenesis | 1.60E-05 |
| Other | uberon 0000033 | head | 2.96E-07 |
|  | uberon 0005886 | post-hyoid pharyngeal arch skeleton | 0.03545473 |
|  | uberon 0001708 | jaw skeleton | 0.0342243 |
|  | uberon 0008895 | splanchnocranium | 0.01593721 |
|  | uberon 0002280 | otolith | 0.01730733 |
|  | GO:0060037 | pharyngeal system development | 0.02837219 |

Supplementary Table S28. Some of the enriched GO-BP and Uberon terms that are related to fin to limb transition for the zebrafish orthologs of the hindlimb module-specific genes. The terms are organized into specific anatomical regions.

| Anatomical region | Term Identifier | Term name | P-value |
| --- | --- | --- | --- |
| Fin related | uberon 4000164 | caudal fin | 0.0046263 |
|  | uberon 0012438 | blastema of regenerating fin/limb | 0.00591299 |
|  | uberon 2001456 | pectoral fin endoskeletal disc | 0.0328624 |
|  | GO:0035118 | embryonic pectoral fin morphogenesis | 0.00553237 |
|  | GO:0031101 | fin regeneration | 0.00239174 |
| Other | uberon 0000033 | head | 2.90E-15 |
|  | uberon 0005886 | post-hyoid pharyngeal arch skeleton | 3.02E-05 |
|  | uberon 0001708 | jaw skeleton | 0.00287797 |
|  | uberon 0005884 | hyoid arch skeleton | 0.01978595 |
|  | uberon 0011611 | ceratohyal bone | 0.0328624 |
|  | uberon 0002280 | otolith | 0.01008715 |
|  | GO:0060037 | pharyngeal system development | 0.02837219 |
|  | GO:0048701 | embryonic cranial skeleton morphogenesis | 1.03E-05 |
|  | GO:0048703 | embryonic viscerocranium morphogenesis | 9.92E-04 |
|  | GO:0060021 | palate development | 0.01953552 |
|  | GO:0060037 | pharyngeal system development | 0.01986392 |
|  | GO:0048840 | otolith development | 0.03165263 |

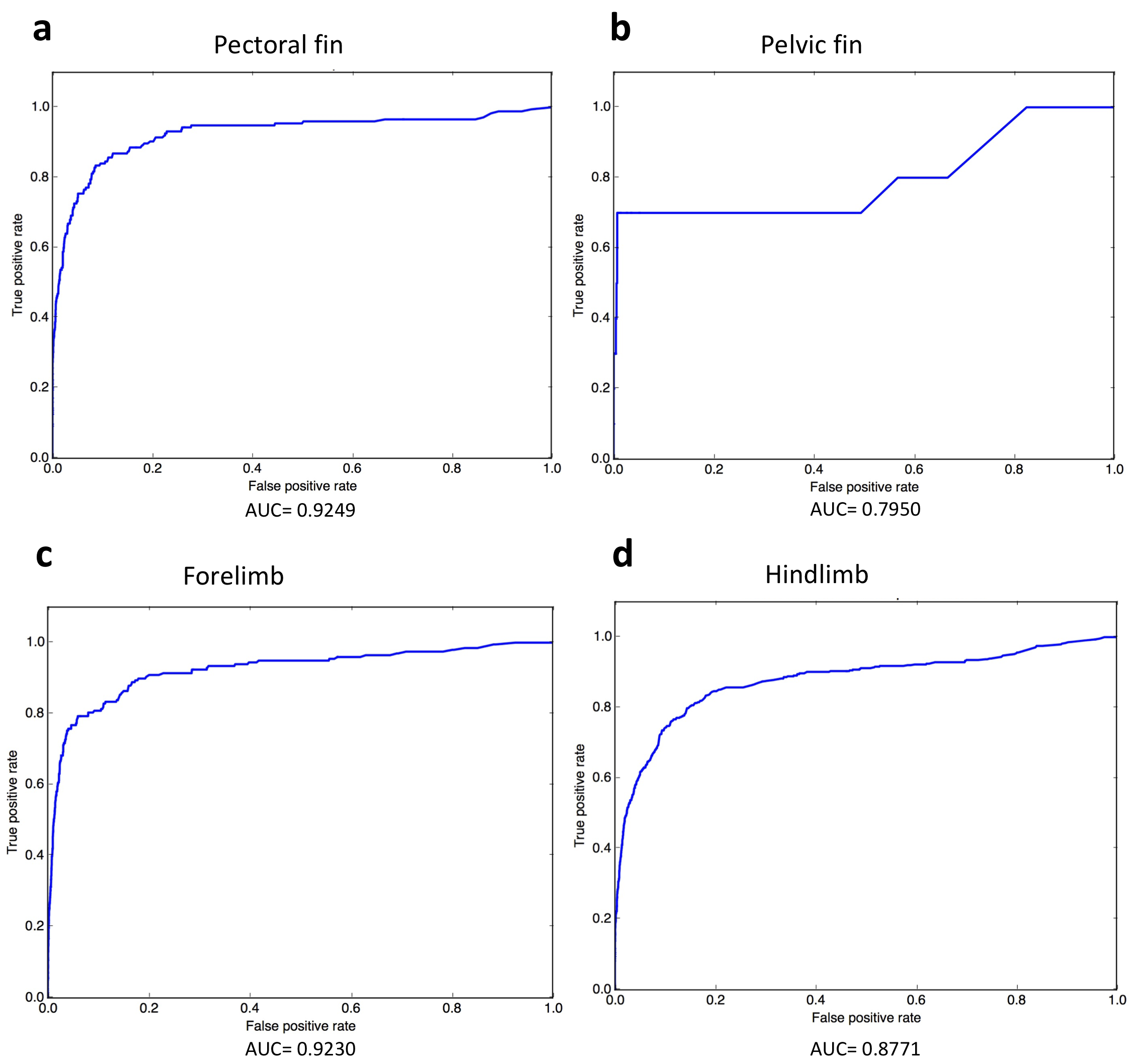

Supplementary figure S1. The ROC curves for the four anatomical entities generated during network-based candidate gene prediction evaluations.

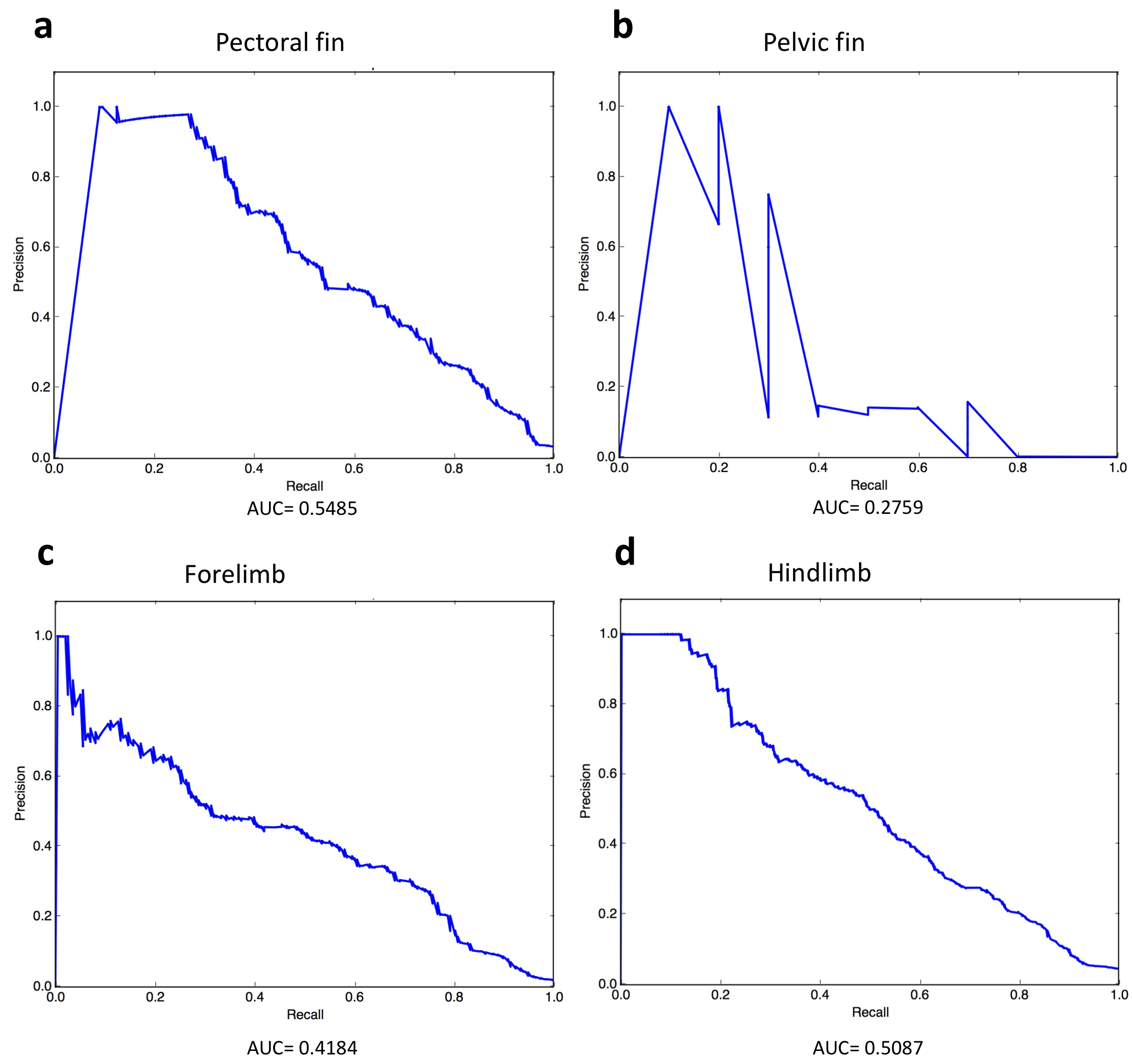

Supplementary figure S2. The precision-recall curves for the four anatomical entities generated during network-based candidate gene prediction evaluations.

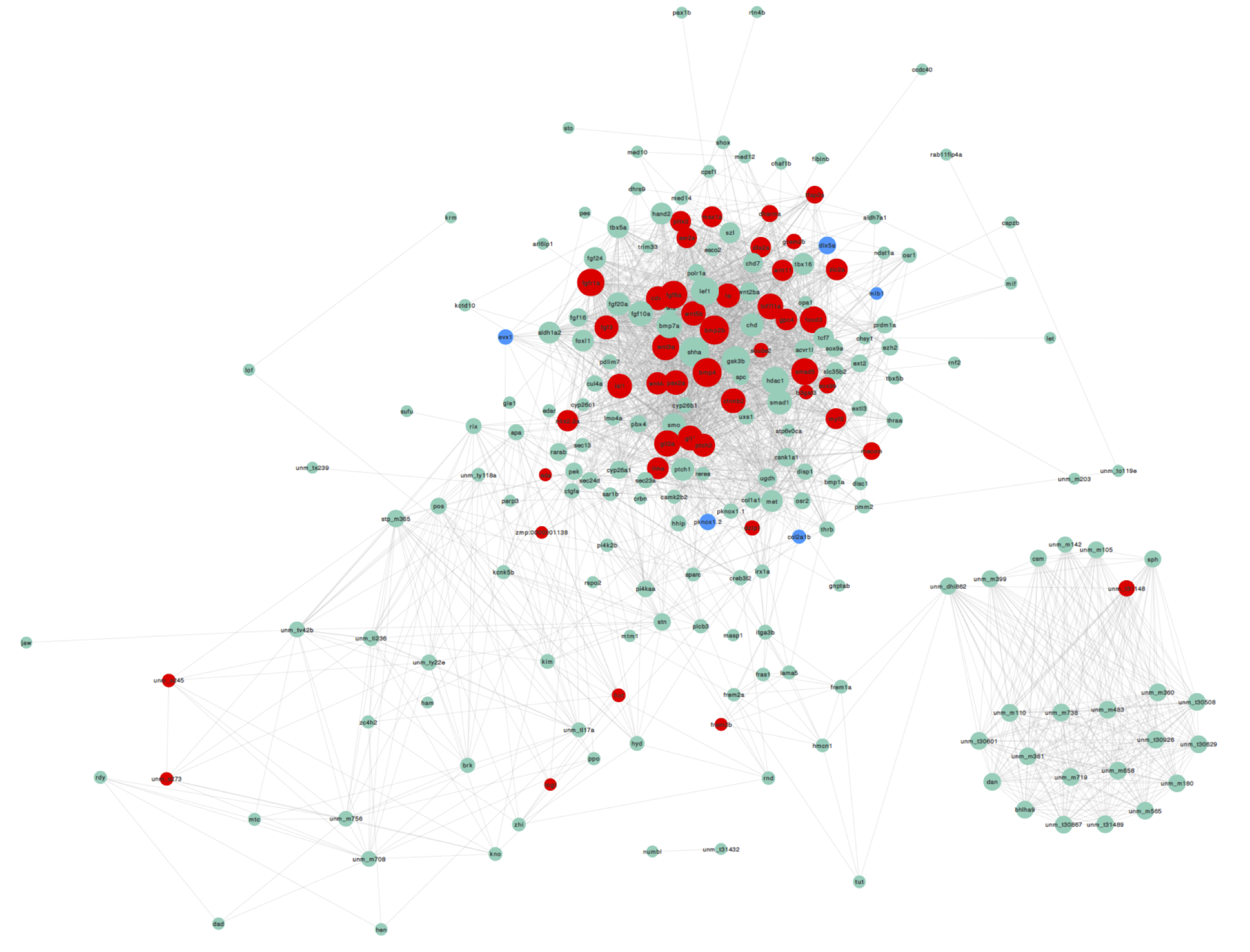

Supplementary figure S3. Visualization of the pectoral fin module, including genes with direct annotations to the pectoral fin (green), genes annotated only to the pectoral fin parts or developmental precursors (blue), and predicted genes (red). Node size is proportional to the degree (number of interactions) of the gene. An interactive version of this module is available in electoronic supplementary material, file S1 as a Cytoscape network file.

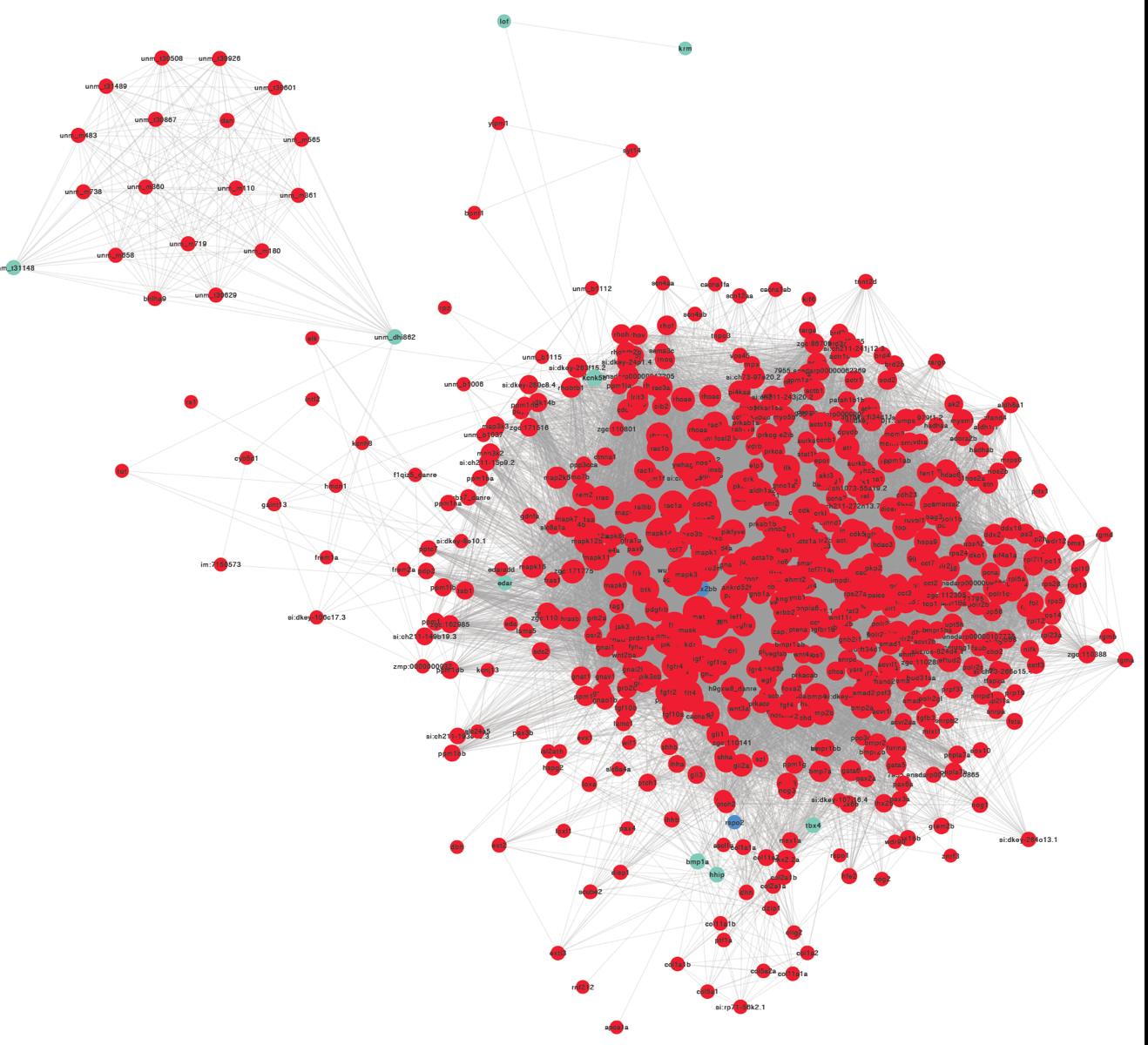

Supplementary figure S4. Visualization of the pelvic fin module, including genes with direct annotations to the pelvic fin (green), genes annotated only to the pelvic fin parts or developmental precursors (blue), and predicted genes (red). Node size is proportional to the degree (number of interactions) of the gene. An interactive version of this module is available in electoronic supplementary material, file S2 as a Cytoscape network file.

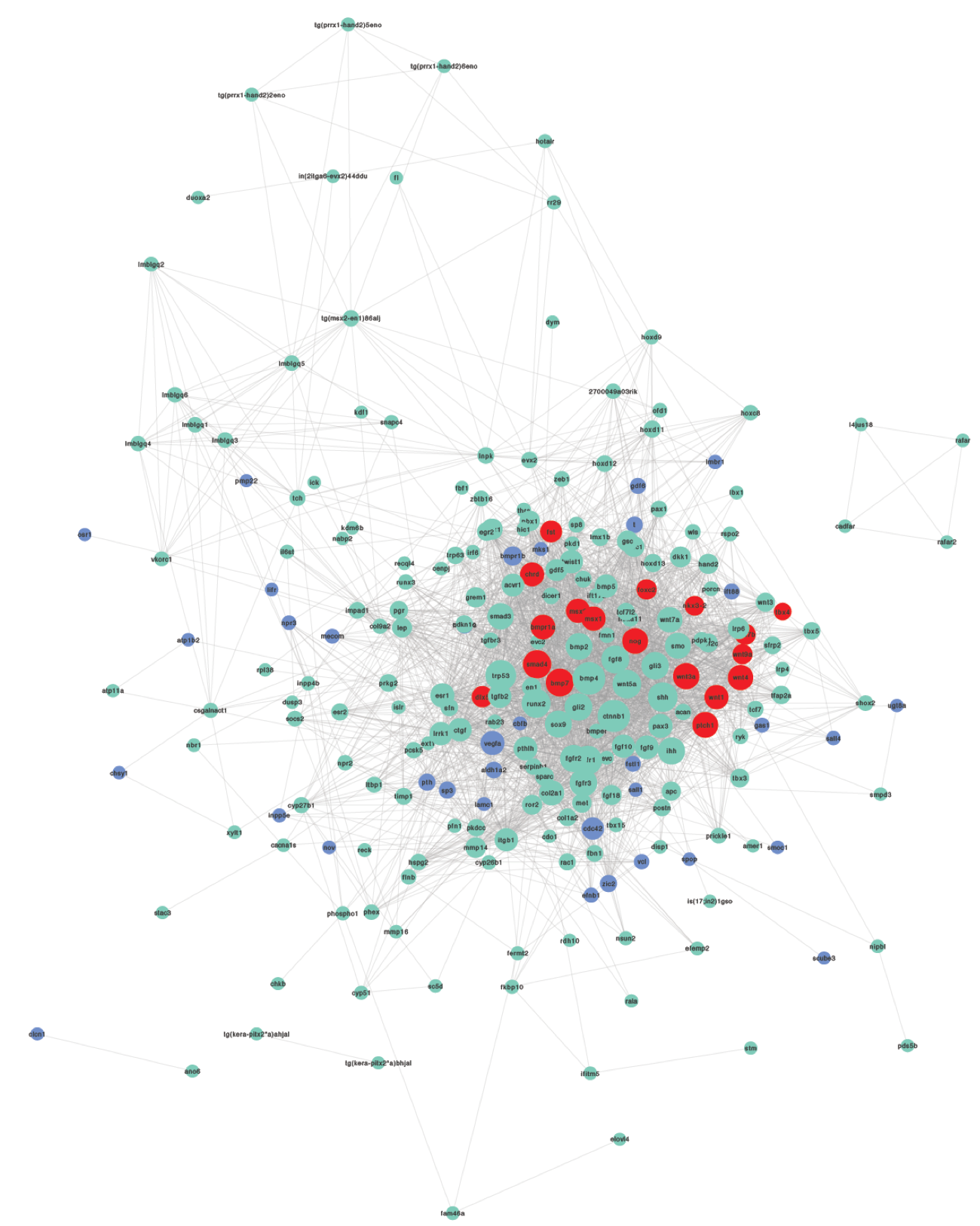

Supplementary figure S5. Visualization of the forelimb module, including genes with direct annotations to the forelimb (green), genes annotated only to the forelimb parts or developmental precursors (blue), and predicted genes (red). Node size is proportional to the degree (number of interactions) of the gene. An interactive version of this module is available in electoronic supplementary material, file S3 as a Cytoscape network file.

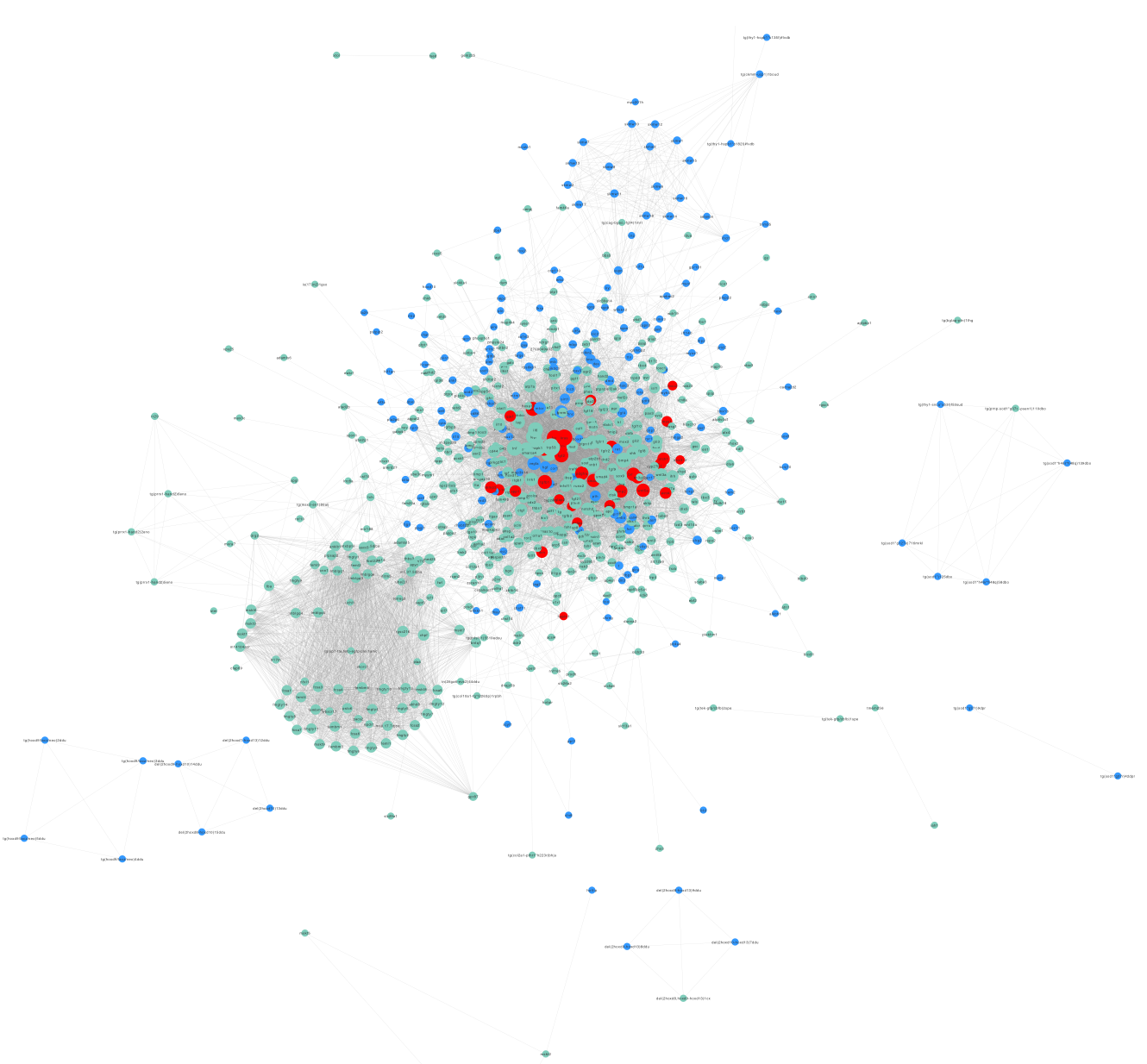

Supplementary figure S6. Visualization of the hindlimb module, including genes with direct annotations to the hindlimb (green), genes annotated only to the hindlimb parts or developmental precursors (blue), and predicted genes (red). Node size is proportional to the degree (number of interactions) of the gene. An interactive version of this module is available in electoronic supplementary material, file S4 as a Cytoscape network file.
